# MOSAIC: A Unified Trait Database to Complement Structured Population Models

**DOI:** 10.1101/2022.03.09.483599

**Authors:** Connor Bernard, Gabriel Silva Santos, Jacques A. Deere, Roberto Rodriguez-Caro, Pol Capdevila, Erik Kusch, Samuel J L Gascoigne, John Jackson, Roberto Salguero-Gómez

## Abstract

1. The ecological sciences have joined the big data revolution. However, despite exponential growth in data availability, broader interoperability amongst datasets is still needed to unlock the potential of open access. The interface of demography and functional traits is well-positioned to benefit from said interoperability. Trait-based ecological approaches have been criticised because of their inability to predict fitness components, the core of demography; likewise, demographic approaches are data-hungry, and so using traits as ecological *shortcuts* to understanding and forecasting population viability could offer great value.
2. Here, we introduce MOSAIC, an open-access trait database that unlocks the demographic potential stored in the COMADRE, COMPADRE, and PADRINO open-access databases. MOSAIC data have been digitised and curated through a combination of existing datasets and additional taxonomic and/or trait records sourced from primary literature. In its first release, MOSAIC (v. 1.0.0) includes 14 trait fields for 300 animal and plant species: biomass, height, growth determination, regeneration, sexual dimorphism, mating system, hermaphrodism, sequential hermaphrodism, dispersal capacity, type of dispersal, mode of dispersal, dispersal classes, volancy, and aquatic habitat dependency. MOSAIC also includes species-level phylogenies for 1,359 species and population-specific climate data where locations are recorded.
3. Using MOSAIC, we highlight a taxonomic mismatch of widely used trait databases with existing structured population models. Despite millions of trait records available in open-access databases, taxonomic overlap between open-access demographic and trait databases is <5%. We identify where traits of interest to ecologists can benefit from database integration and start to quantify traits that are poorly quantified (*e.g.,* growth determination, modularity).
4. The MOSAIC database evidences the importance of improving interoperability in open-access efforts in ecology as well as the need for complementary digitisation to fill targeted taxonomic gaps. In addition, MOSAIC highlights emerging challenges associated with the disparity between locations where different trait records are sourced.

---

> “I’m not interested in your data; I’m interested in merging your data with other data.

> Your data will never be as exciting as what I can merge it with.”

> —Sir Tim Berners-Lee (co-founder of the World Wide Web)

## Introduction

The ecological sciences have recently joined the big data revolution (Farley, Dawson, Goring, & Williams, 2018; Marx, 2013; Reichman, Jones, & Schildhauer, 2011). Total species distribution records measure in the hundreds of millions (Maldonado et al., 2015; Troia & McManamay, 2016). Functional trait data exist for tens of thousands of species across the globe (Díaz et al., 2016; Enquist et al., 2019; Maitner et al., 2018). Similar gains have taken place in related fields, from big omics (Field et al., 2009) to global state-of-the-art climate models (*e.g.*, ERA-5 [https://www.ecmwf.int/en/forecasts/dataset/ecmwf-reanalysis-v5]) becoming available at fine spatial and temporal resolutions suitable for biological analyses (Davy & Kusch, 2021; Hersbach et al., 2020), and from behavioural trait data (Rowcliffe, Jansen, Kays, Kranstauber, & Carbone, 2016) to population data (Global Population Dynamics Database [https://knb.ecoinformatics.org/view/doi:10.5063/AA/nceas.167.15]; Human Mortality Database (J.R. Wilmoth, K. Andreev, D. Jdanov, 2007; mortality.org); Human Fertility Database (Jasilioniene et al., 2015; humanfertility.org); AnAge Database (De Magalhᾶes & Costa, 2009); DATLife [Max Plank Institute of Demographic Research 2022; https://datlife.org/]). We have also seen growth datasets characterising phenomena at levels of biological complexity above the organismal level, including community ecology databases (BioTime (Dornelas et al., 2018); Web of Life (Fortuna, Ortega, & Bascompote, 2014); metaCommunity Ecology: Species, Traits, Environment and Space (CESTES) (Jeliazkov et al., 2020); Environmental Data Initiative [https://environmentaldatainitiative.org/]). The growth of records in ecology datasets and complementary environmental datasets is enabling us now to test ecological theory at larger and more complex scales. In line with open-access growth, however, there is a recognised need to improve and coordinate data standards to guide the systematic collection and management of data across different trait collection programmes (Gallagher et al., 2020; Kissling et al., 2018; Wilkinson et al., 2016).

Despite the increasing availability of biological data, combining different datasets into common analyses is often hampered by the lack of complementarity between databases. Interoperable systems reflect the transformation of constituent data types into a standardised format of spatial, temporal, and measurement scales, accounting for differences in methods (Nadrowski et al., 2013). The need to improve interoperability across datasets is demonstrated by the widescale emergence of data harmonisation initiatives across fields of ecology (*e.g.,* Culina et al., 2018; Kattge et al., 2020; Schneider et al., 2019). In the past decade, dozens of initiatives have taken shape to both centralise data from existing datasets and to improve data interoperability: standardising units, scales, and terminology for comparative purposes (Edwards, Lane, & Nielsen, 2000; Maurer, Firestone, & Scriver, 2000). Unifying data formats and streamlining their integration unlocks the potential for existing datasets to answer questions that cut across levels of biological complexity. Linking together levels of biological complexity is critical for identifying how phenomena emerge and transmit across different levels of biological organisation, upscaling and downscaling through biological systems. For example, the critical linkages between genetics and biochemistry (The ENCODE Project Consortium, 2012; http://bigg.ucsd.edu/), biochemistry and physiology (Vitousek, Johnson, & Husak, 2018), and physiology and demography (Laughlin, Gremer, Adler, Mitchell, & Moore, 2020; Swenson et al., 2020) have benefitted from dataset integration.

The limitations of global-scale datasets are shifting away from data availability and toward data interoperability. The global-scale integration of functional traits is headed in this direction. Momentum toward data integration is evidenced by recent database initiatives (*e.g.,* Guerrero-Ramírez et al., 2021) and global networks that, like the Open Trait Network (https://opentraits.org/), aim to standardise and integrate trait data across taxa (Gallagher et al., 2020). Despite major improvements in the consolidation and accessibility of trait data, there is not yet a single-source centralised database spanning behaviour, physiology, habitat, and other trait data for a wide range of species. Existing databases are often linked by taxonomy (*e.g.,* FishBase (Froese & Pauly, 2010), CoralTraits (Madin et al., 2016), MammalBase (Lintulaakso, 2013), AmphiBio (Oliveira, São-Pedro, Santos-Barrera, Penone, & Costa, 2017)); trait type (*e.g.,* TreeOfSex (The Tree of Sex Consortium 2014), TreeBase (Boettiger & Temple Lang, 2012), Xylum Functional Traits (Borghetti, Gentilesca, Colangelo, Ripullone, & Rita, 2020)); data type (*e.g.,* GBIF (https://www.gbif.org/), MOL (Jetz, Thomas, Joy, Hartmann, & Mooers, 2012), TetraDensity (Santini, Isaac, & Ficetola, 2018)); or a combination of the taxonomies and traits (WooDiv (Monnet et al., 2021), CarniDiet (Middleton, Svensson, Scharlemann, Faurby, & Sandom, 2021)). A number of other databases take a more general approach in their thematic scope, but are still constrained to a limited set of traits and taxonomy (*e.g.,* Amniote (Myhrvold et al., 2015), Pantheria (Jones et al., 2009), BIEN (Enquist, Condit, Peet, Schildhauer, & Thiers, 2016; Maitner et al., 2018), TRY (Kattge et al., 2020)). MOSAIC offers a platform that integrates databases in the remit of species with structured population models across their ecological traits.

Here, we introduce MOSAIC, a centralized database of trait data across the Tree of Life. MOSAIC is an open-access database that complements the existing demographic data available in the COMPADRE Plant Matrix Database (Salguero-Gómez et al., 2015), COMADRE Animal Matrix Database (Salguero-Gómez et al., 2016), and the new PADRINO IPM Database (Levin et al., BioRxiv; doi: 10.1101/2022.03.02.482673; https://levisc8.github.io/RPadrino/). MOSAIC v. 1.0.0 includes 14 frequently used traits that encompass morphological, reproductive, dispersal, and habitat type attributes for some 300 species of animals and plants. Additional traits will be added in the future (see *Future Direction, below*). MOSAIC allows users to integrate structured population data to probe larger questions through the collection, curation, and complementarity of relevant contextual data.

## Methods

### Scope and Coverage of MOSAIC

The MOSAIC database (Version 1.0.0) includes 14 key trait records across 300 species (Fig. 1). MOSAIC is designed to provide complementary data for analysis in connection with structured population models: matrix population models (MPMs, Caswell, 2000), where state variables are discrete (*e.g.*, age (Morris et al., 2008), ontogeny/development (Smallegange, Deere, & Coulson, 2014), discrete classes of size (Crouse, Crowder, & Caswell, 1987)), and integral projection models (IPMs, Easterling, Ellner, & Dixon, 2000), where the state variables are continuous (*e.g.,* size (Jongejans, Shea, Skarpaas, Kelly, & Ellner, 2011), mass (Ozgul et al., 2010), parasite load (Metcalf, Graham, Martinez-Bakker, & Childs, 2016)). The traits included in MOSAIC 1.0.0 were identified by the MOSAIC team as a set of physical, physiological, geographic, and behavioural attributes of most immediate relevance to demographic research (see Table 1; more information at https://mosaicdatabase.web.ox.ac.uk). Traits were also selected in consideration of the lack of standardised and centralised databases for certain traits (*e.g.,* volancy, modularity, and growth indeterminacy; see Fig. 2 for trait variance and taxonomic structure of select traits excluded from existing databases). Importantly, we note that MOSAIC is not a general dataset for analysis of functional traits, as this is available through other extensive repositories (*e.g.,* TRY (Kattge et al., 2020), BIEN (Enquist et al., 2016; Maitner et al., 2018)). Instead, the focus of MOSAIC is on providing taxonomic coverage to species with open-access structured population models available in the COMADRE (Salguero-Gómez et al., 2016), COMPADRE (Salguero-Gómez et al., 2015), and PADRINO databases (Levin et al. forthcoming) (See Fig. 3 and Fig. 4 for spatial scope and taxonomic scope with respect to structured population databases, respectively). MOSAIC provides a much-needed interoperability between existing databases that are germane to demography. By doing so, MOSAIC helps to fill critical data needs of population ecologists and functional trait ecologists (see Fig. 5 for relevant covariance structure). Although demographically motivated, MOSAIC affords opportunities for data to address novel questions more broadly across ecology, evolution, and conservation biology.

**Figure 1.**
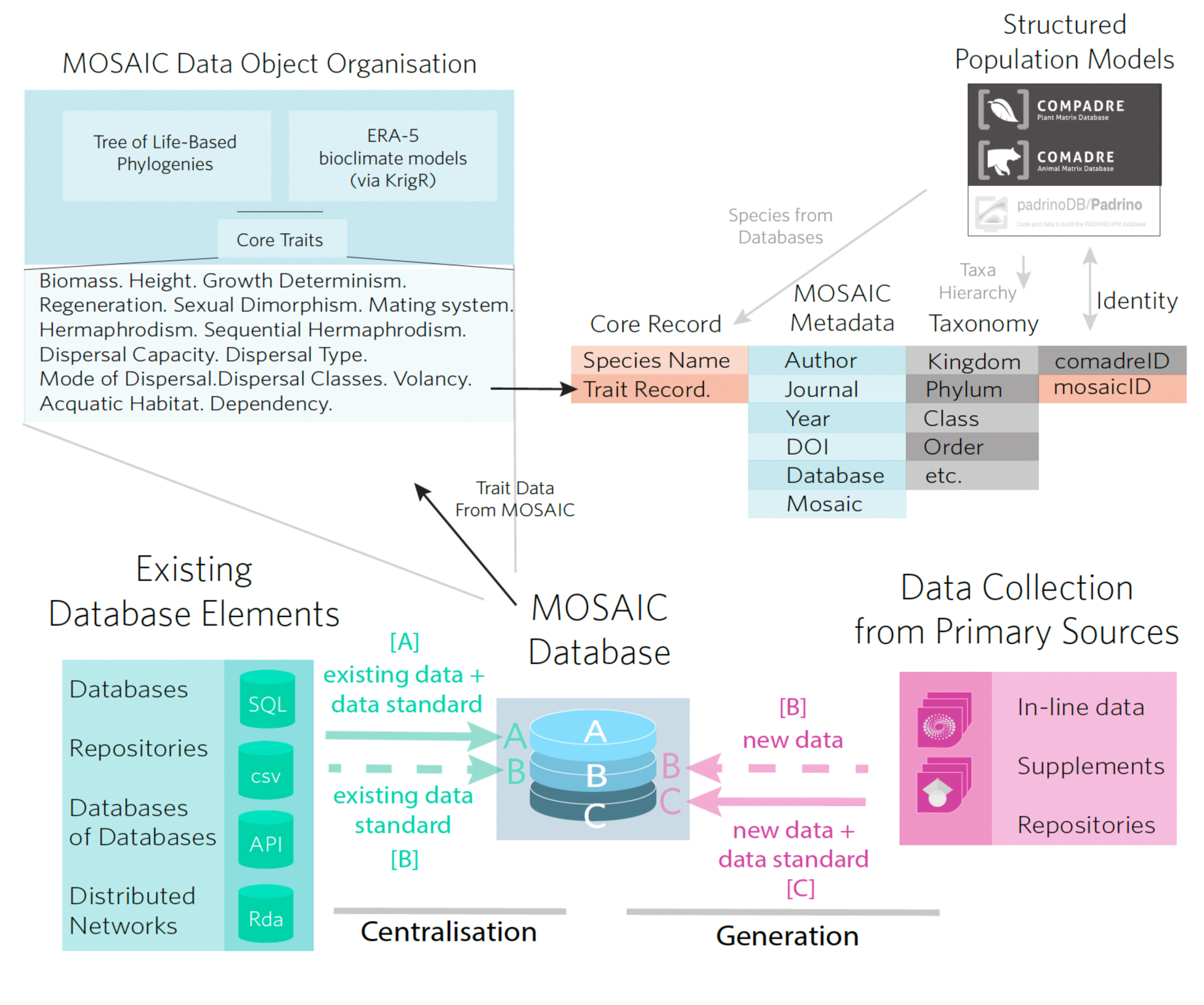
Structure of the MOSAIC database (v1.0.0). The MOSAIC database comprises a combination of existing trait records centralised from data servers, databases, and datasets and new records collected from the primary literature by the MOSAIC team. Existing records are labelled, MOSAIC-A, in the mosaic metadata attribute field and include provenance in the metadata. New data records that adhere to existing data standards and that fit into the remit of an existing database (*i.e.*, a data gap filled by the MOSAIC team) are labelled MOSAIC-B. New records that do not have an existing data standard or for which there is not an existing database are labelled MOSAIC-C.

**Figure 2.**
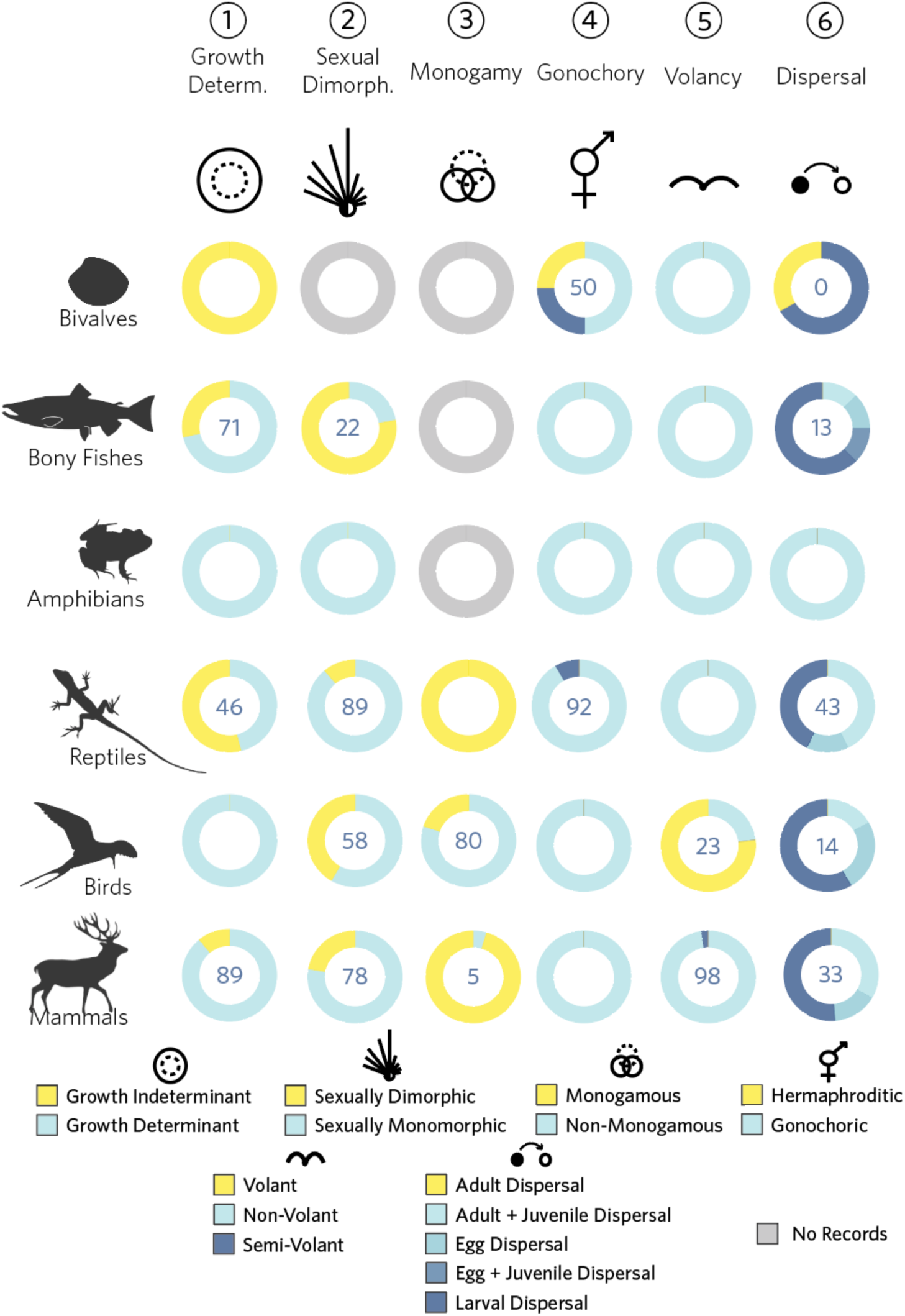
Trait fields in the MOSAIC database (v1.0.0). The MOSAIC database contains a combination of continuous and discrete trait fields. Six of the 14 trait fields are illustrated here for animals, and organised by trait level (all discrete fields) and taxonomic classification. Trait distributions vary strongly by trait and taxonomy. Observed variation in trait values by taxonomic group is a prerequisite condition to their potential value toward explaining reported variation in vital rates and resulting demographic properties across the Tree of Life. Species included in the MOSAIC database are limited to those for which stage-structured population data exist in the COMADRE, COMPADRE, and PADRINO databases.

**Figure 3.**
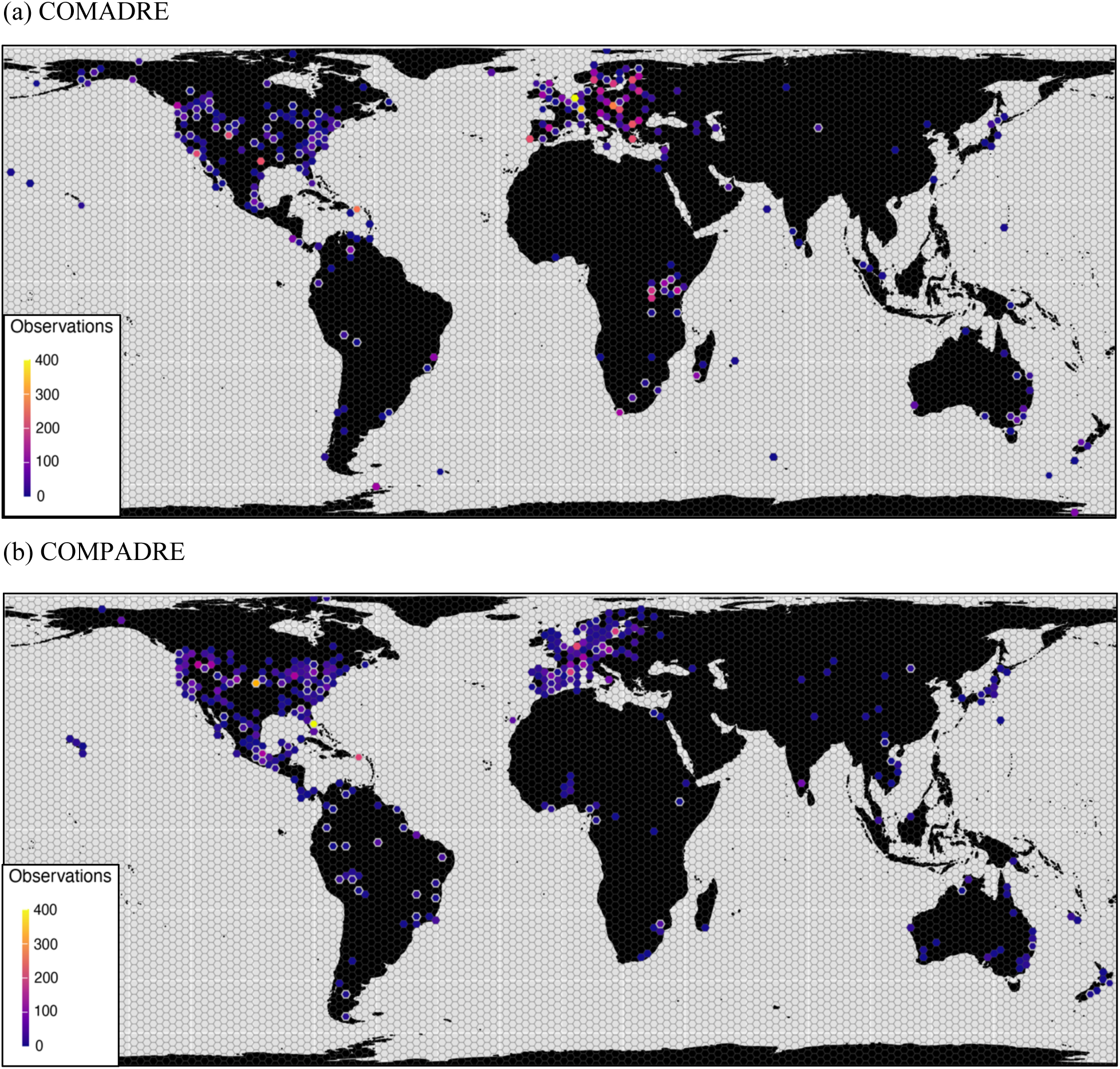
The spatial distribution of MOSAIC (v1.0.0) records relative to the (a) COMADRE and (b) COMPADRE databases. The COMADRE and COMPADRE dataset include animal and plant demographic data, respectively, on all continents except Antarctica, with a substantial bias to North America and Central Europe. MOSAIC introduces trait records that include models in all major geographies of COMADRE and COMPADRE. The map shows the density of matrix population models globally (color graded by density per area in *ca.* 150 km^2^ hexagonal cells). Cells bordered by white represent localities of population models for which there are trait records included in the first release of the MOSAIC trait database. Species prioritised in v1.0.0 of MOSAIC did not include the PADRINO dataset, so no distribution figure is shown for integral projection models.

**Figure 4.**
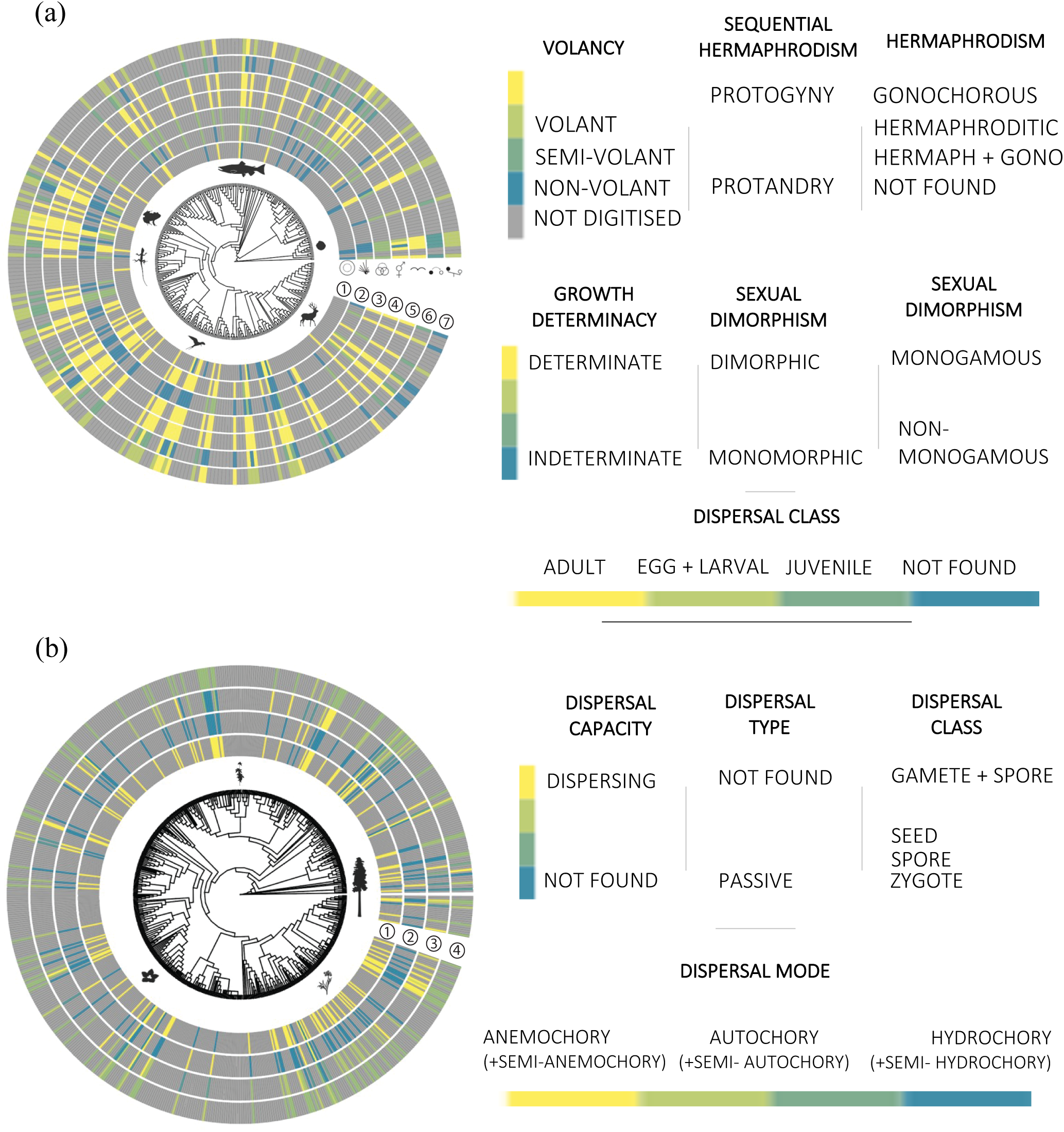
Phylogenetic mapping of traits included in MOSAIC (v1.0.0). (a) Phylogenetic distribution of seven major traits for animals: (1) growth determination; (2) sexual dimorphism; (3) monogamy; (4) gonochory; (5) volancy; (6) dispersal; and (7) biomass. (b) Phylogenetic distribution of four traits in plants: (1) dispersal capability; (2) dispersal type; (3) dispersal mode; and (4) dispersal class. Figures developed in R using the ggtree package.

**Figure 5.**
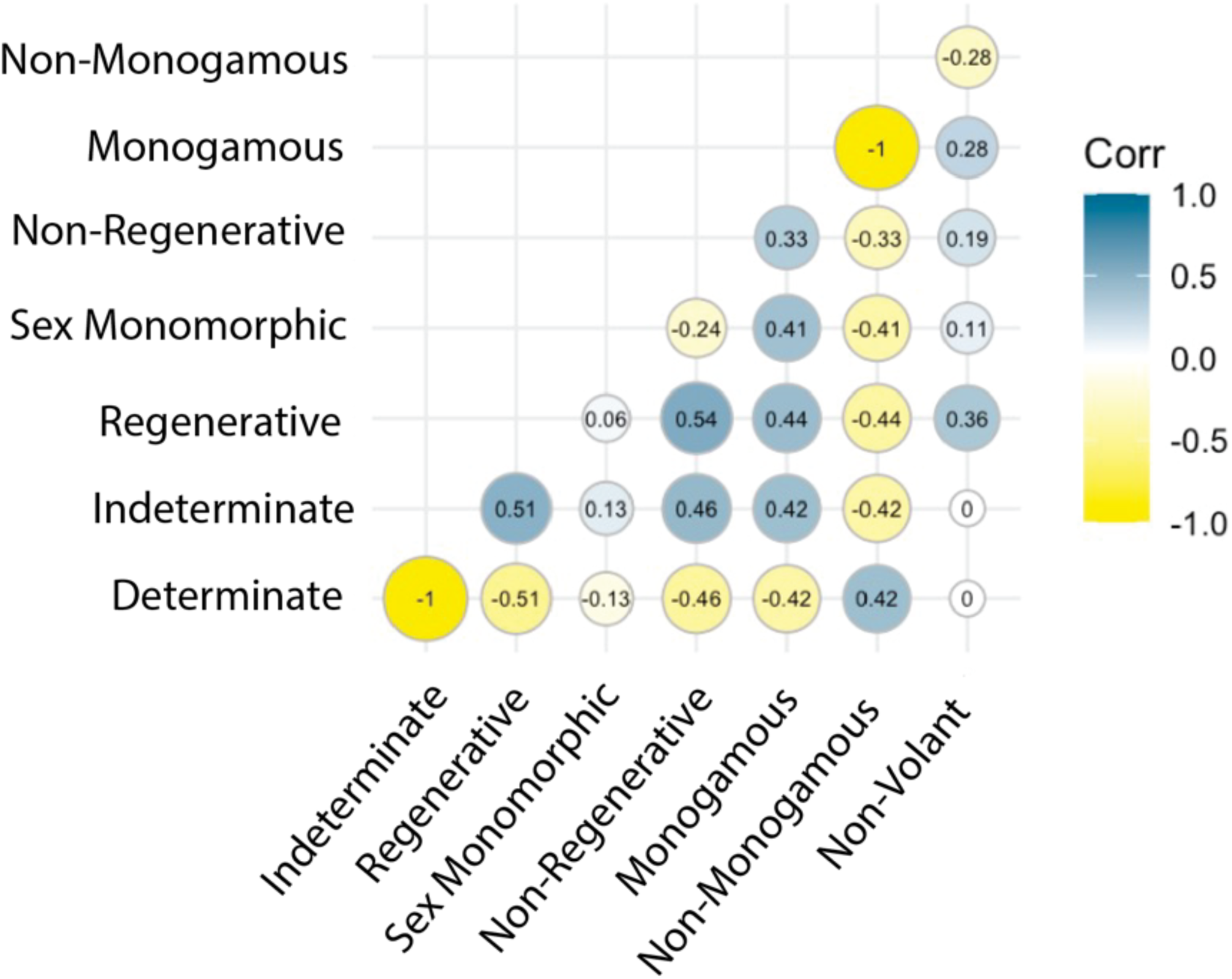
Trait covariance in the MOSAIC database (v1.0.0). Some traits exhibit high correlation in the MOSAIC database, such as between growth determination and between growth regeneration. Trait associations are expected to occur in the MOSAIC dataset and may reflect widespread constraints or statistical anomalies, particularly when dealing with small samples or specific taxonomic subgroups. Trait covariation can be symptomatic of biomechanical constraints (*e.g.*, flight and biomass), major growth characteristics (*e.g.*, modularity and growth determination), or other past or presently compelled associations (*e.g.*, height and vessel density).

**Table 1:**
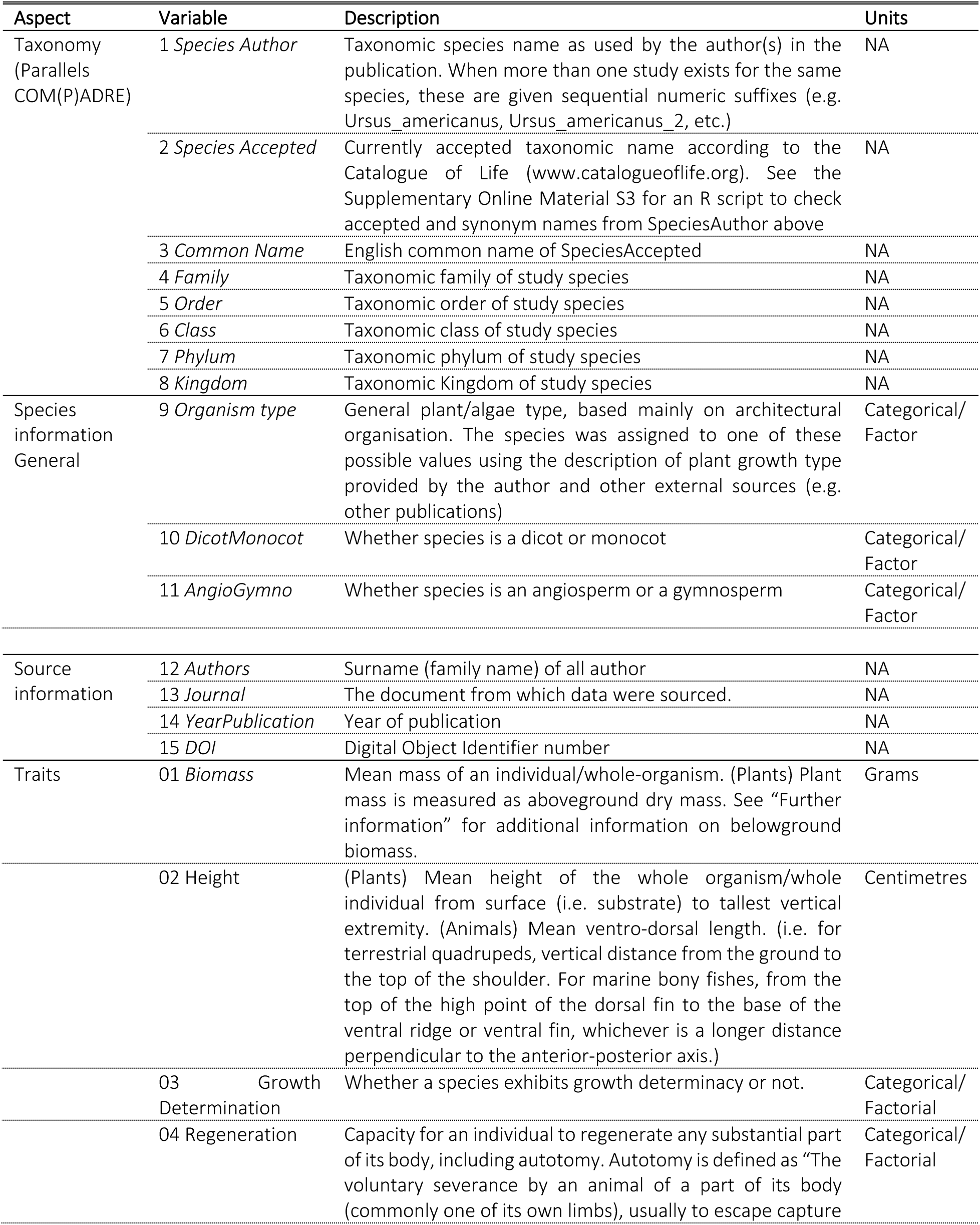
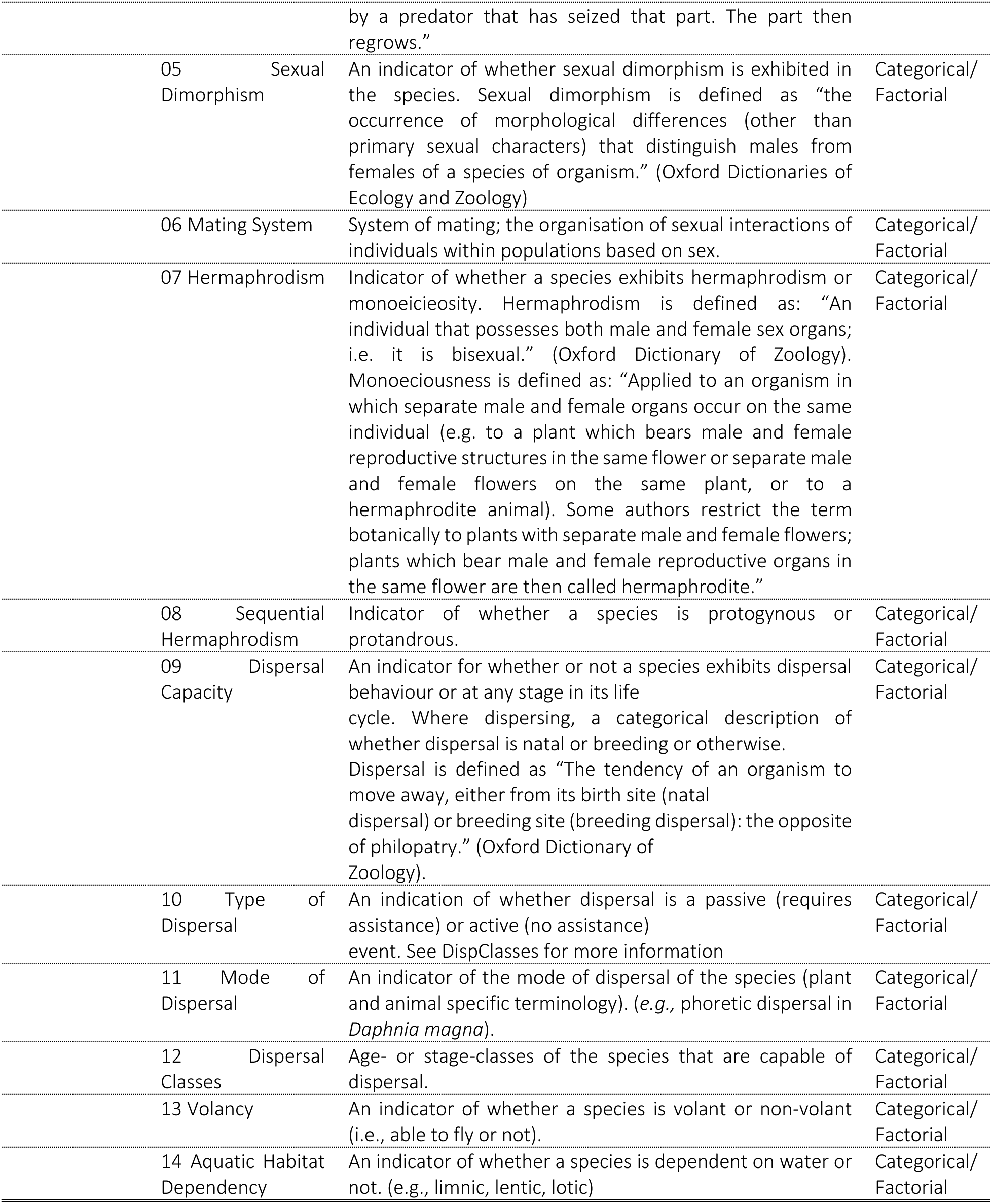
Variable names and meaning contained in the MOSAIC Database, organised into seven general trait domains. A more detailed description can be found in the MOSAIC user guide found at: https://mosaicdatabase.web.ox.ac.uk/.

### Accessing MOSAIC

A guide for downloading the MOSAIC database can be accessed on the MOSAIC portal (https://mosaicdatabase.web.ox.ac.uk). The MOSAIC database is open access and can be downloaded into R using a few lines of code (see: https://mosaicdatabase.web.ox.ac.uk/download-database), which download the MOSAIC data object as an S4 object (Rda file). The database can also be downloaded from GitHub directly as an (.Rda) or comma separated value (.csv) file and loaded into R manually.

The MOSAIC portal hosts information pages and downloadable files about the database. Users can download the *MOSAIC User Guide*, *vignettes*, and *data collection protocols* from the portal webpage (https://mosaicdatabase.web.ox.ac.uk/vignettes and https://mosaicdatabase.web.ox.ac.uk/user-guide). The *User Guide* specifies the classifications, precision, and data types for each trait field. The MOSAIC *User Guide* details the metadata on the structure of data for each trait (*e.g.,* species, genus, mosaic index). An updated list of the databases that are directly or indirectly addressed by the MOSAIC database is maintained on the MOSAIC portal.

### Data Sources

MOSAIC is both a *meta-*database (a database of databases) and a database in its own right, containing new trait records from primary literature (Fig. 1). The MOSAIC database contains records centralised from existing datasets where functional traits relevant to population ecologists can be openly accessed and redistributed (*e.g.,* BIEN (Enquist et al., 2016; Maitner et al., 2018)). The records reflected in the MOSAIC database do not encompass the entirety of the source databases but are instead partial facsimiles of those databases that reflect records relevant to population databases (COMADRE, COMPADRE, PADRINO). MOSAIC has three major components: (1) trait records sourced from existing databases (22%); (2) trait records newly procured through searching the primary literature (71%); and (3) trait record markers that direct users to additional records that are not included in the MOSAIC database (7%). MOSAIC trait markers exist for one of two reasons: the database containing the records of interest does not allow records to be accessed or limits redistribution rights behind individual registration and/or specific use applications; or records in other databases contain multiple records for a species, which do not currently fit within the data structure of version 1.0.0 of MOSAIC (see *Future Targets for MOSAIC*).

In addition to identifying whether a trait record is new, the “MOSAIC” attribute field also identifies whether the attribute field (*i.e.,* trait name) is part of an existing database. For example, the MOSAIC field might indicate that a record for specific leaf area is new for a specific species, and also that the attribute is part of databases such as TRY (Kattge et al., 2020) or BIEN (Enquist et al., 2016; Maitner et al., 2018). By contrast, a new species record for volancy would indicate that there are currently no databases that systematically collect data on volancy attributes and therefore that all volancy records in the MOSAIC database are new.

#### Organisation of Sources in MOSAIC

The MOSAIC attribute field is a factorial variable with three levels: MOSAIC-A, MOSAIC-B, and MOSAIC-C. The first of these levels, MOSAIC-A (Fig. 1), labels only records that reflect existing databases (*i.e.*, provenance of an existing data acquisition service); the second, MOSAIC-B, labels new records collected by the MOSAIC team that are in a trait field within the scope of an existing database initiative (*e.g.,* specific leaf area in BIEN); and the third, MOSAIC-C, labels new records collected by the MOSAIC team on traits for which there is not currently a database initiative centralising records. If a datum has been adopted from another dataset or database, then the relevant source is referenced in the *Database* attribute column. Note that this value will be “NA” for all MOSAIC-C records, logically. Over time, data sharing will move records in MOSAIC-B to MOSAIC-A as the MOSAIC-B traits are assimilated in the database networks that specialise on an existing trait (*i.e.,* data feedback; see Fig. 1).

#### Sources and Provenance of Records

Because of existing limitations on data access, some datasets cannot be transferred into MOSAIC. Where data exist outside of the MOSAIC platform, but have restricted access, the MOSAIC database directs users toward the appropriate database on a trait and taxa-specific level (see `metaMOSAIC`). The *MOSAIC User Guide* (Appendix S1) explains differences between data gaps that are yet to be reviewed and those that are true gaps (*e.g.,* volancy/flight capability of plants). Data were obtained through searching peer-reviewed records of ISI Web of Science, Scopus, and Google Scholar using key words pertinent to the species name and trait field in question (see Appendix S2 for a list of keywords queried). The MOSAIC team also searched the archives of data repositories, including the Dryad digital repository (http://datadryad.org; see Appendix S3 for a complete list of repositories reviewed – to be maintained hereafter on the MOSAIC portal), and reviewed in detail journal archives that have a high occurrence of data publishing, including *Nature Scientific Data*, *Methods in Ecology and Evolution*, and *Journal of Ecology*. MOSAIC also reflects a review of data from data aggregating servers, such as the open traits network (https://opentraits.org/), the ecological data wiki (https://ecologicaldata.org/), environmental data initiative (https://environmentaldatainitiative.org/), and databases that aggregate other databases (*e.g.,* BIEN (Enquist et al., 2016; Maitner et al., 2018) and TRY (Kattge et al., 2020)). The complete set of key words used in this review is detailed in a supplement to the *User Guide* posted on the MOSAIC portal. A current list of databases reviewed in the development of MOSAIC is included in Appendix S3. Suggested data sources and key terms can be submitted through the MOSAIC data portal.

### Database Navigation

The *User Guide* contains detailed information on navigating the MOSAIC R data object. In general, the data object can be queried with the syntax: @values[[mosaic.index]]. In the aforementioned syntax, the `trait.name` and `mosaic.index` should reflect one trait and one or more species index values in the respective locations (*e.g.,* mosaic@volancy@value[[93]]). The complete set of records for a trait can be viewed and assigned by calling the trait from the mosaic object without an index specified (*e.g.,* mosaic@volancy@value). The mosaic index value reflects a unique identifier that can be linked from COMADRE, COMPADRE, and PADRINO through the index field of mosaic (*e.g.,* comadre@data$MatrixID[comadre@data$SpeciesAccepted=="Alces alces"])). Associated metadata for each record can be queried by the following syntax: @metadatafield[[mosaic.index]]. Here, the `metadatafield` should reflect one metadata element (*e.g.,* mosaic@volancy@author[[93]]). The complete metadata for a species can be viewed using the `mos_meta()` function or searched manually by slots, or by querying rows of the database in the comma separated value format (https://mosaicdatabase.web.ox.ac.uk/navigating-mosaic). See the *User Guide* (Appendix S1) for more information. The database is organised to be readily subsetted, including compatibility with Tidyverse package (Wikham et al. 2019) syntax (see *Navigating MOSAIC*; https://mosaicdatabase.web.ox.ac.uk/vignettes, also in Appendix S4). The MOSAIC object can be searched in `R` by taxonomy, trait, locality, duration of study, sex, and other metadata fields applicable to COMADRE, COMPADRE, and PADRINO (see *vignettes*, above, also Appendix SX). Trait values in the database are measured in standardised units of measure for each trait (Table 1). Species names are specified in conformance with the Catalogue of Life (www.catalogueoflife.org), also consistent with the COMPADRE, COMADRE, and PADRINO databases. The *User Guide* provides specific guidelines for querying fields within the data object in R and for negotiating the dataset in finer detail (https://mosaicdatabase.web.ox.ac.uk/user-guide).

### Error Reporting

The MOSAIC portal has an *Error Report* page for reporting potentially erroneous records, or to query additional questions (https://mosaicdatabase.web.ox.ac.uk/suggested-additions-error-reporting; but see also FAQ: https://mosaicdatabase.web.ox.ac.uk/frequently-asked-questions). Potential errors can be reported anonymously or with contact information (*e.g.*, name, email). Decisions on reported errors will be disclosed on the Error Report page (https://mosaicdatabase.web.ox.ac.uk/suggested-additions-error-reporting) and to the reporting party if contact information is included in the request. If it is not possible to submit errors via the portal, users can also to reach out by email to.

### Recommended Records

The MOSAIC portal has a *Recommended Records* page (https://mosaicdatabase.web.ox.ac.uk/suggestion-additions) for reporting suggested records that are not included in the MOSAIC database. Recommendations can be made with or without contact information. Contact information will be used exclusively for clarifying questions and updating the commenter when records are included. MOSAIC will report decisions on the *Recommendations Incorporated* page (https://mosaicdatabase.web.ox.ac.uk/suggestion-additions). Users may also request new data fields to be prioritised in future rollouts. Given the realities of limiting resources, The MOSAIC team will do their best to include the requested records in future versions. Likewise, we also welcome submissions of data collected by individuals via.

### *metaMOSAIC*: licensed data, access limitations, and restricted redistribution of existing records and databases

Not all datasets permit open use, dissemination, and redistribution of their trait data. Where limitations on the data collation and redistribution apply, there may be application procedures, registration, and other actions necessary for an individual to obtain access to specific trait records for analysis (*e.g., TRY*). MOSAIC centralises the metadata for datasets that do not allow data to be redistributed to help navigate to relevant data resources outside the scope of open access. Records in limited access databases can be searched in MOSAIC by taxonomic group and by trait. MOSAIC links researchers to application materials for requesting access to those limited-access databases. The data access of licensed or non-open-access databases is stored in a data object called `metaMOSAIC` that is an extension of the MOSAIC database. The MOSAIC dataset thus provides data where it is accessible and `metaMOSAIC` guides researchers to where data exist with registration. When searching fields in the MOSAIC database, the `metaMOSAIC` adjunct dataset indicates if data are available in these ancillary sources (see *User Guide* in Appendix S1 for more information) and provide links to pertinent sources and data through the provider. `metaMOSAIC` is part of the main database object accessible through the MOSAIC portal.

### Database Updates

The MOSAIC database will be updated as new data are added to the COMADRE, COMPADRE, and PADRINO databases. Updates of the MOSAIC database will account for newly discovered data sources and new literature that adds to or changes the species-level traits in the database, as well as correct errors from earlier versions. New MOSAIC versions will be released periodically with a notice published on the website, in the data object metadata, the mosaic GitHub page (https://github.com/mosaicdatabase), and through updates in associated packages in conformance with standard semantic versioning (a three-part version code reflecting major, minor, and patch updates, in respective positions). Updates will be published to the mosaic portal (https://mosaicdatabase.web.ox.ac.uk), social media (Twitter: @mosaicdatabase), and GitHub (https://github.com/mosaicdatabase). The MOSAIC portal includes a location for submission of recommendations for additions or suggested changes to the database.

### Future Targets of MOSAIC

#### Versioning

MOSAIC database updates and version history will be posted to the MOSAIC portal in the future.

#### Taxonomic Scope

Future versions of MOSAIC are scheduled for expansion to include 50 core traits across some 1,500 species (the current entirety of COMPADRE, COMADRE, and PADRINO; *MOSAIC target traits*; see *User Guide* [Appendix S1]). The MOSAIC core traits list will guide the future roll-out of attributes gathered and stewarded by the MOSAIC team (available on the MOSAIC portal website; see below), but additional trait suggestions can be submitted through the online portal.

#### Interspecific Variation in Hierarchical Data Structures

Future versions of the MOSAIC database will adopt a hierarchical file structure that will accommodate multiple records per species. Once records for COMADRE, COMPADRE, and PADRINO are fully populated across the MOSAIC traits with mean, pooled, or other representative quantities (*e.g.,* mean leaf size for all plants or adult bodymass of animals), secondary records will be added. Existing trait databases may contain multiple records per species (see, for example, structure of COMADRE (Salguero-Gómez et al., 2016), COMPADRE (Salguero-Gómez et al., 2015), BIEN (Enquist et al., 2016; Maitner et al., 2018), TRY (Kattge et al., 2020)), although some databases host a single record per species’ trait, such as age and growth rate for animals in AnAge (De Magalhᾶes & Costa, 2009). To facilitate research into intra-specific trait variation (Albert, 2015; Messier, McGill, & Lechowicz, 2010; Siefert et al., 2015), MOSAIC provides provenance of records, whether records were subject to selection or merger (means or pooling), and fields that identify whether multiple records are known to exist. Where records for a given species were isolated from existing databases, mean values are often retrieved, and the database sourcing additional data is noted in the database under the attributed field (“Additional Trait Data Available” field).

#### citMOSAIC: Citizen Science

In addition to `metaMosaic`, which guides users to licenced data not reported in the MOSAIC database, MOSAIC plans to roll out a database of identical structure to MOSAIC that gathers information from citizen science datasets. Like MOSAIC, citMOSAIC will have three components (citMOSAIC-A, citMOSAIC-B, citMOSAIC-C), reflecting the same relationship of databases and fields to MOSAIC (see https://mosaicdatabase.web.ox.ac.uk/citizen-mosaic for more information). citMOSAIC will be kept independent of the main MOSAIC database to avoid conflation of peer reviewed literature from citizen science data. Where appropriate to use datasets together, the metadata, query functionality, and design of citMOSAIC will mirror that of MOSAIC to promote interoperation of databases. citMOSAIC will be downloadable from the MOSAIC portal website.

### Cautionary Notes

Records in the MOSAIC database are gathered and standardised under the protocols detailed in the *User Guide* (Appendix S1; also available through the MOSAIC portal). Users should be attentive to the precision, levels, and context of data in MOSAIC when used for analysis. For those records in the dataset that come from multiple individuals, we present them as statistical components (*e.g.,* minimum, maximum, or mean trait values). Functional traits in MOSAIC may be estimated from populations studied in COMADRE, COMPADRE, PADRINO, or other databases (Table S3). The potential temporal and spatial mismatch between databases that are linked in an analysis merits close attention (Csergő et al., 2017). The studies in the MOSAIC database also include research conducted by different investigators using independent tools, technologies, sample designs, and study methods. The influence of research methods and instruments on the error values in the dataset may require additional consideration for potential bias, noise, and imprecision. Where more than one life history trait value exists for a given species, MOSAIC users will need to determine whether averaging or selective filtering to one study is most appropriate in view of the specifics of the given research question. In certain cases, trait values for a species might only be available for a single st/age and therefore may not provide a complete picture of the trait variation amongst st/ages. Users are encouraged to be cautious when contextualising the scope of representation of the values in the database and their analyses.

### Representation, Variance, and Bias in MOSAIC v1.0.0

MOSAIC can help to mechanistically examine the causes and correlates of demographic variability across the thousands of animal and plant species housed in COMPADRE, COMADRE, and PADRINO. In its version 1.0.0, MOSAIC contains a high degree of variance in trait values known to shape demographic outcomes across major taxonomic groups. For example, determinant growth is present for 0% of Amphibians and Birds and 100% of Bivalves, while volancy is present for 0% in Reptiles, Amphibians, and Bony Fish and 77% in Birds (Fig. 2). Animal adult biomass and plant height follow lognormal trait distributions (see Appendix S5). 11% of mammals are indeterminate growers *vs.* 54% of reptiles and 0% birds, and 95% of mammals are monogamous compared to 100% of reptiles and 20% of birds in MOSAIC. Recent studies have examined how different vital rates are explained by functional traits (e.g., Carmona, Bello, Azcarate, Mason, & Peco 2019; Pistón et al. 2018; Adler et al. 2014; Poorter et al. 2008). However, understanding how trait variation across taxa translates to demographically influential properties remains underdeveloped.

MOSAIC’s initial release (v1.0.0) includes records for all major regions of the globe for which we have structured population models (Fig.3). Nevertheless, species trait values are not necessarily gathered from the same localities as population models (see *future directions* for more information on systematising spatial mismatch). This is an important consideration for users of MOSAIC (and more generally of trait-based approaches) wishing to bring together functional traits and demographic rates, as traits and vital rates are known to vary considerably within the same species across spatial scales (Violle et al. 2012; Albert et al. 2010; Bolnick et al. 2011). Moreover, while there is at least one trait for each of these locations, the data density remains variable. Thus, records are not necessarily representative of the global spread or the full spatial scope of MOSAIC. For example, the highest complete coverage for MOSAIC traits is concentrated toward localities with the with the longest-term demographic models (see COMPADRE locations associated with MOSAIC records).

Phylogenetically, the initial release of MOSAIC is somewhat limited. Version 1.0.0 covers 300 of the 1,400 species currently available in COMPADRE, COMADRE and PADRINO. However, MOSAIC trait data are well distributed across clades (Fig. 4). While there is not a highly skewed phylogenetic concentration with respect to the existing structured population models or clustering of records into small groups across the Tree of Life, phylogenetic density of MOSAIC records remains low. Therefore, information from the MOSAIC database may be limited for a given genus or order and, as such, should be approached with caution. In future versions of MOSAIC, the phylogenetic bias is expected to diminish with more samples and stronger phylogenetic representation.

Covariance across traits is also an important source of confound in existing analyses linking traits and vital rates. Positive and negative correlations across traits that have independent influences on vital rates can create apparent associations of demographic properties with traits, spuriously functionalising non-functional traits (McElreath 2021). Disentangling the relevance of key axes of trait variation for their demographic influences demands a clear quantification of the direction and strength networks of trait associations, trade-offs, and demographic consequences. Population biologists seek to understand not only how individual traits relate to different aspects of demographic performance (*e.g.,* population growth rate, risk of quasi-extinction, *etc.*), but also understanding how trait syndromes shape those demographic outcomes. MOSAIC presents a highly varied covariance structure in trait values for the examined 300 species. For example, without *a priori* expectation, indeterminate growth and regeneration traits show strong correlation (r=0.51; Fig. 5), which could influence each other’s effects on vital rates. The same could be argued for the correlation between volancy and reproductive strategy (r=0.28 with monogamy; Fig. 5).

The MOSAIC database can be used as a platform to showcase the lack of overlap between trait and vital rate data for the same species. This picture calls for a more systematic way address global biases in ecological data quantification/collection. Even where we have complete information about species in the COMADRE, COMPADRE, and PADRINO databases, we are subject to the constraints and biases of those datasets, such as spatial bias toward high-GDP countries and the phylogenetic bias toward temperate regional perennial plants (Kendall et al. 2019; Römer et al. 2021). The compounding of error and density across datasets highlights the need to prioritise stronger representation of functional traits linked with demography. The standardised framework of MOSAIC is an ideal platform to work from to achieve this goal.

In view of potential biases introduced by low sampling density and the patchiness of cover in traits, users of the database are advised to consult the literature about the representativeness and congruency between MOSAIC data and related trait diversity within clades. Users need to be mindful of the scope of the questions that they are setting out to answer and to be aware of the influence of sample sizes and coherency or heterogeneity of traits across different taxonomic levels.

## Discussion

### From Databases to Data Networks

Broad aperture digitisation efforts (*e.g.,* BIEN (Enquist et al., 2016; Maitner et al., 2018) and TRY (Kattge et al., 2020)) have helped resolve many answers to demographic questions. Examples include whether there are trait spectra and key trade-off patterns amongst functional traits and whether these are correlated with particular environments and life history strategies (Laughlin, 2018; Reich et al., 2003; Wilkes et al., 2020), Trait-based ecology and Trait Driver Theory (Enquist et al., 2015) are indebted to such opportunity-driven research programmes. More generally, however, the trait-based ecology paradigm has failed to support clear answers to many research questions of central interest to demographers (Bellier, Kéry, & Schaub, 2018; Laughlin et al., 2020). This limited reach of the functional trait programme coincides with a dearth of species-specific overlap across the range of functional traits that are collected by the functional trait databases.

The proliferation of databases and open data initiatives over the last two decades (Cheruvelil & Soranno, 2018) evidences an interest in improving both data access and data usability (Gallagher et al., 2020; Michener, 2006; Whitlock, 2011). While existing databases standardise trait fields, collate records, and link associated metadata, existing databases often store data for simple, quantitative traits. Relatively few ecological trait databases store diverse data types (such as rate arrays, population time series, physiological rates at different structural levels, and habitat shape files) that may be associated with multidimensional, ecological study systems (but see CESTES (Jeliazkov et al., 2020), GFBio (Diepenbroek et al., 2014), DarwinCore (Wieczorek et al., 2012)).

The digitisation and standardisation of *existing* data and their integration with complementary, new data presents a growing set of challenges and opportunities in ecology (Salguero-Gómez, Jackson, Gascoigne 2021). Efforts to gap-fill records can leverage the value of existing datasets while expediting the schedule for specific research outcomes. As trait datasets grow, the importance of targeted, gap-filling initiatives to address bias and to capitalise on existing data will also increase (Conde et al., 2019). The value of existing records is further enhanced through improvements in the interoperability of datasets. Much of this work is done manually, at a high cost, and with little support from funding agencies (Salguero-Gómez, Jackson, Gascoigne 2021), and yet it has been effective at facilitating research and creating new value for old data. In recent years, initiatives have sought to improve the interoperability of datasets by guiding prospective data structure or retroactively harmonising existing datasets. These include programmes that develop universal standards to improve global interoperability (such as DarwinCore (Wieczorek et al., 2012) and Frictionless [https://frictionlessdata.io/]data standards) or that contain guidelines for data metastructure (such as the FAIR principles (findability, accessibility, interoperability, and reusability, *sensu* Wilkinson et al., 2016) and the OpenTraits framework (Gallagher et al., 2020)). These initiatives address emerging and scaling challenges of ecoinformatics, such as the protocols by which we share data, search data, and preserve provenance within hierarchical data storage structures. These protocols will be essential in centralising datasets as diverse as government monitoring datasets (*e.g.,* those stored in U.S. Data clearing houses [https://www.data.gov; https://www.dataone.org/]; National Biodiversity Atlas [https://nbnatlas.org/]); centralised monitoring and experimental networks (*e.g.,* LTER and NEON), raw or reanalysed remote sensing datasets (*e.g.,* Landsat data, NASA EarthData datasets, ERA-5 data), and private datasets (https://www.natureserve.org/) that will demand versatile and navigationally efficient data structures.

Population ecology has benefitted from widespread open-access databases but requires further dataset integration to answer its central questions. Understanding whether and how some morphological or physiological traits predict demographic outcomes and why others fail to do so is of central interest to questions in physiological, population, and community ecology (Adler et al., 2014; Chalmandrier et al., 2021; Laughlin et al., 2020; Yang, Cao, & Swenson, 2018). Population ecologists routinely use data that are distributed across a wide range of databases. Comparative and macroecological researchers use phylogenies (*e.g.*, Daskalova, Myers-Smith, & Godlee, 2020; Levin, Crandall, Pokoski, Stein, & Knight, 2020; Roper, Capdevila, & Salguero-Gómez, 2021), adult bodymass (*e.g.*, Capdevila et al., 2020; Healy et al., 2014; Terry, O’Sullivan, & Rossberg, 2022; Williams, McRae, Freeman, Capdevila, & Clements, 2021), and high-resolution, global climate information (*e.g.*, Daskalova, Bowler, Myers-Smith, & Dornelas, 2021; Paniw, Maag, Cozzi, Clutton-Brock, & Ozgul, 2019; Stenseth & Mysterud, 2002) to answer relevant biological, evolutionary and ecological questions and to contextualise their findings. Population ecologists frequently examine a subset of physiological, morphological, and behavioural attributes associated with demographic outcomes ("functional traits", *sensu* Violle et al., 2007). The trait-based research programme seeks to, among other aims (*e.g.*, Díaz et al., 2016), identify the intrinsic and extrinsic regulators of vital rates and the causes of variation and constraints on possible trait values (Buckley & Puy, 2022; Salguero-Gómez, Violle, Gimenez, & Childs, 2018). Not all traits predict demographic outcomes and functional traits may exercise influence on only a few demographic pathways (Pistón et al., 2019; Salguero-Gómez & Laughlin, 2021). The answers to these questions rely on the existence of vital rate and trait data, the overlap of which has been limited in the absence of targeted attention. For instance, of the hundreds of thousands of records available across thousands of plant species in TRY (Kattge et al., 2020) and over 345 plant species in COMPADRE (Salguero-Gómez et al. 2015, COMPADRE Ver 1.0.0), Adler et al. (2014) report functional trait-vital rate relationships for only 222 plant species due to their limited data overlap.

Ecological data are complex and their structures will need consistent rules to link datasets together. It will be important for future datasets to adopt hierarchical database designs that render large, thematically, and structurally diverse data to be readily locatable and usable without expert knowledge. Here, we show one such example in the scope of comparative research, using thematic groups and a strategy of achieving adequate record breadth before revisiting depth of records for specific species. The need for open access data, integrated workflows, and interoperable data systems is increasing with the scaling of data collection through use of robotics and technologies. The gaps in existing data systems, interoperability, and data acquisition can be filled strategically for specific applications, offering targeted and efficient dataset development. With data interoperability guiding the structure of new datasets, the modular development of area-specific datasets will enable more generalised use over time and help meet the aims of existing database initiatives.

## ACKNOWLEDGEMENTS

We thank Mark Roper, Sara Middleton, Etienne Cousin, and Thomas Marrien for their contributions to discussion at the beginning of the project, Jessica Hass and Lauren Hinchcliffe for collecting data for the project, John Park for his contributions to the conception of citMOSAIC, and Sam Levin for valuable comments on the Rmosaic package. This work was supported by a NERC IRF (NE/M018458/1) to RS-G. RCRC acknowledges postdoctoral support from the Regional Valencian Government and the European Social Fund (APOSTD/2020/090).

## S1: MOSAIC User Guide

USER GUIDE TO THE MOSAIC LIFE HISTORY DATABASE

A Companion to COM(P)ADRE and PADRINO/A demographic databases

*Working document – 06 February 2022*

## General Instructions

### Database Organization

The data associated with MOSAIC are provided in a single R data (extension.Rdata) and as a comma separated value (.csv) file format. The code for downloading the data can be located from the mosaic portal (https://mosaicdatabase.web.ox.ac.uk/download-database) In addition, these files are accompanied by R scripts and a nexus phylogeny available in the Supplementary Information of the manuscript introducing MOSAIC, and in our GitHub repository (https://github.com/mosaicdatabase/mosaicdatabase)

MOSAIC_v_1_0_0.RData: Contains basic information regarding the source of publication, as well as ecological, biogeographic, and taxonomic details of the demographic study for each study species, the demographic information (i.e., the matrix population model) and metadata.

### Database Design

In developing the MOSAIC database, we balance level of detail with accessibility. A highly detailed, comprehensive profile of life history traits for species in COM(P)ADRE and PADRINO/A, if possible to collect, would be difficult to navigate. So rather than collating a wealth of information in many different formats, we designed this dataset to highlight a smaller collection of traits in a single format which is of most interest, and expressed, by COM(P)ADRE and PADRINO/A users. In MOSAIC’s future updates, we plan to add additional detail and additional fields, but we initially took a limited approach and plan to keep the design minimal. If there are life history traits, alternative formats for existing variables, or other features you would like to see added to MOSAIC, please suggest them to us at

Understanding the diverse needs of users, we include in this guidance document additional direction on obtaining information on the variables we report in more detail. In this guide, we also highlight the scope of use, and caution against the most foreseeable abuses of data. We ask that all users approach this dataset with a caution and pay close attention to what variables and their statistical expression reflect.

### The Meanings of NA and NDY in MOSAIC

NA in the MOSAIC data generally means that the data are not applicable. An example of where the data are not applicable is volancy within plants, as this trait does not occur in plants. NDY in the MOSAIC data means that the data have not yet been digitized. NF in the MOSAIC data means that the data are not available to date as no affirmative records were found upon review.

### Disclaimer

The MOSAIC digitization team does its best to ensure data accuracy, and every piece of information goes through multiple error-checks prior to its release in www. mosaicdatabase.web.ox.ac.uk. However, we claim no responsibility for any damage that may arise from using MOSAIC. A list of error checks and potential issues in the use and interpretation of the database are described in the main manuscript. The end user is ultimately responsible for his/her interpretations of the data.

**What is new in this version?**

Version 1.0.0

- The first version of the database. No updates.

#### DIAGRAM OF THE MOSAIC DATABASE ARCHITECTURE

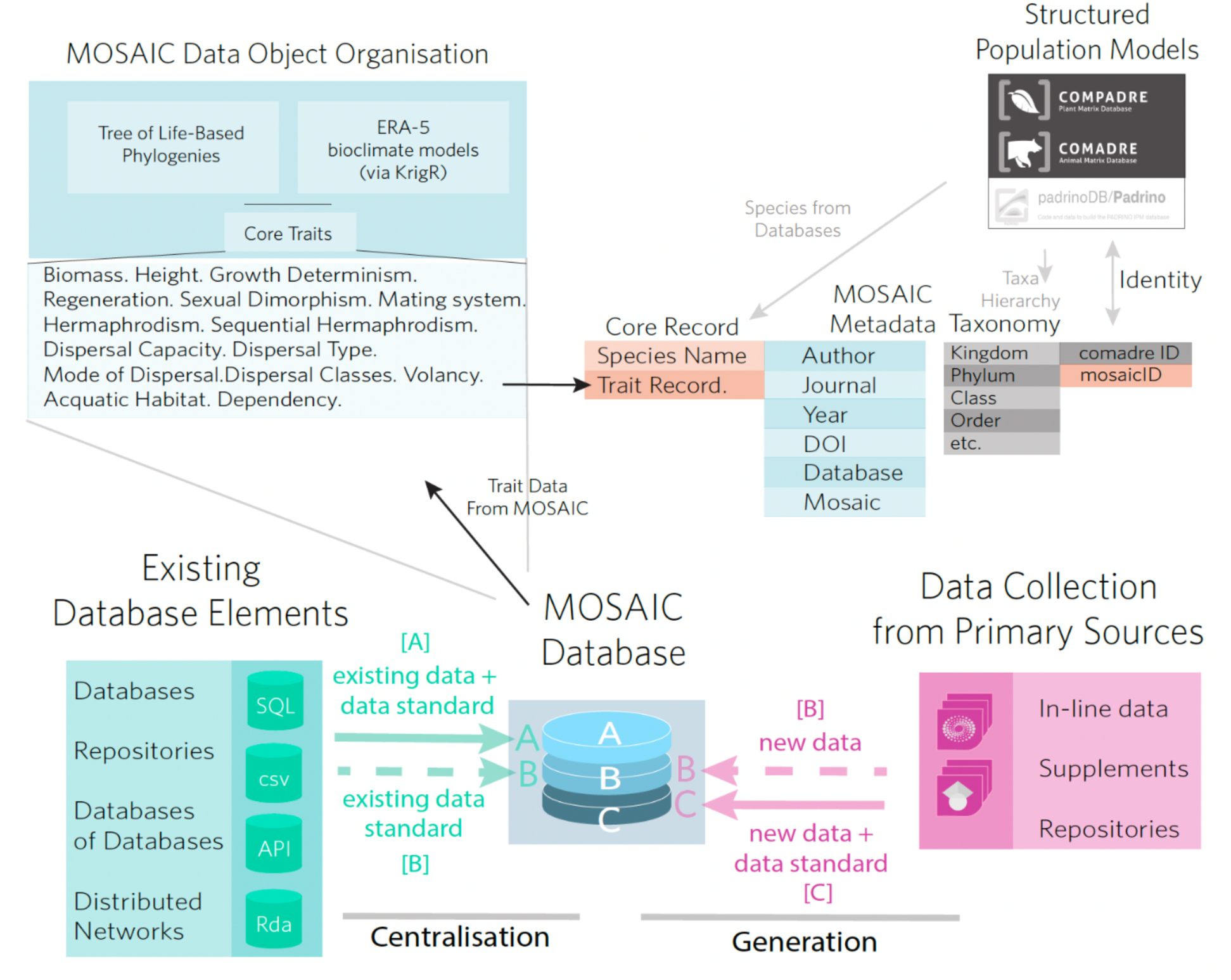

### Variables in MOSAIC

The MOSAIC database is constructed of objects containing life history information across variables and organized into themes to aid in navigation. The metadata object is the object containing information about every study for which data is stored in the MOSAIC database. Every variable containing a value in the MOSAIC database will have corresponding metadata.

Associated with every data record is a value and its corresponding metadata. The metadata details data providence and relationships to existing databases. The fields within metadata are detailed below:

### Format Guide

**[Index]** *Variable Name*

**Definition**: [Definition of the variable]

**Possible values, [**cat. = categorical/discrete; cont. = continuous], [r variable class: character, numeric, integer, complex, or logical.]

- <XX> [variable with two digits] XXX –XXX [discrete variable name] – [discrete variable definition] <…>XXX<…> [discrete variable with additional content on either side]
- Units: [unit of measure (integer, percent, ratio, mm, km^2^, g, etc.)]
- Precision: [for continuous variables: scientific notation of the precision of the measure - e.g., 1e^1^ for km = precise to the tens of km (3270 km), no decimal; 1e^-1^ for temperature = precise to the tenth of a degree (10.4 °C)]
- Error boundaries: [for continuous variables: boundaries beyond which values are errors]

**Usage Notes**: [Notes on boundaries of use – note that this is non-exhaustive and highlights major potential errors]

**Additional Information:** [Guidance on databases containing more detailed information]

**Source Data: [**Datasets from which information was gathered; note that this will not include individual papers unless the papers are associated with a larger dataset/database. This category reflects databases.]

**Last updated**: [Date that information was retrieved from other databases or searches]

#### A1 *Species Accepted*

**Definition:** Currently accepted latin name.

Possible values, cat., factor

- <genus_species> - e.g., Taraxacum_officinale

**Usage Notes**: NA

**Additional Information**: NA

**Source Data:** This information is obtained from The Encyclopaedia of Life

**Last updated**: 24 January 2022

#### A2 *Kingdom*

**Definition:** Kingdom to which species belongs

**Possible values**, cat., factor

- <kingdom> - e.g., Plantae, Fungi, Rhodophyta, Chromista (yes, MOSAIC includes fungi and algae as well as plants)

**Usage Notes:** NA

**Additional Information:** NA

**Source Data:** The Encyclopaedia of Life

**Last updated**: 24 January 2022

#### B1 *Authors*

**Definition:** Surname (family name) of all authors

**Possible values**, NA, character

- <name(s)> - Separated with “;” e.g., “Smith; Jones”

**Usage Notes**: NA

**Additional Information**: NA

**Source Data**: NA

**Last updated**: 24 January 2022

#### B2 *Journal Name*

**Definition:** The document from which data were sourced.

**Possible values,** cat., factor

- <abbreviated journal name> - Where the data come from a scientific journal article, the abbreviated journal name is given. We use the standard abbreviation of the journal compliant with the ISO-4 standard.
- Book - Records are from a book, or book chapter
- PhD thesis - Records are from a doctoral thesis
- MSc thesis - Records are from a masters thesis
- Report - Records are from a report
- Conference talk - Records are from a conference talk
- Conference poster - Records are from a conference poster

**Usage Notes**: NA

**Additional Information**: NA

**Source Data**: NA

**Last updated**: 24 January 2022

#### B3 *Year Publication*

**Definition:** Year of publication

**Possible values**, cont., numerical

- <YYYY> - e.g., 2012

**Usage Notes**: NA

**Additional Information:** NA

**Source Data:** NA

**Last updated**: 24 January 2022

#### B4 *DOI/ISBN Number*

**Definition:** Digital Object Identifier number

**Possible values**, NA, character

- <XXXXXXXXXXXXXX> - e.g., doi.org/10.1073/pnas.1506215112

**Usage Notes**: NA

**Additional Information**: NA

**Source Data**: NA

**Last updated**: 24 January 2022

### PRIMARY Variables

#### A1 *Biomass*

**Definition:** Maximum reported mass of adult individual/whole-organism. For plants, only aboveground dry mass is measured. See “Additional information” for additional information on belowground biomass.

**Possible values**, cont., numerical

- Units: Grams (g)
- Precision: 0.000
- Error Boundaries: 0-150,000,000g
- NF – Body mass reviewed and inconclusive (no affirmative records found upon review)
- NDY – Body mass not digitised yet (yet to be evaluated)
- NA – not applicable

**Usage notes:** When both male and female data were reported, only the maximum value was considered independently of the gender of the individuals.

**Additional Information:** Belowground biomass is not reported because information availability is appreciably more limited than for aboveground biomass. The BIEN database includes information on belowground biomass that can be referenced and utilized where of interest.

**Source Data:** Amniote, TRY database.

**Last updated**: 24 January 2022

#### A2 *Height*

**Call:** -Morph$Height

**Definition:** (Plants) Maximum height of the whole organism/whole individual from surface (i.e. substrate) to tallest vertical extremity.

**Possible values**, cont., numerical

- Units: centimeters (cm)
- Precision: 0.000
- Error Boundaries: 0-1000cm

**Usage notes:**

- Depending on the species group, height can have profoundly different meanings. Height corresponds with embolic risk in some woody plants and corresponds with size-based fitness in others.

**Additional Information:** NA

**Source Data:** TRY database.

**Last updated:** 24 January 2022

#### A3 *Growth Determination*

**Call:** -Morph$Growth$Determination

**Definition:** Growth indeterminacy is defined by continuous growth of individuals throughout their lifetimes (measured by mass, length, bone ossification, or other indicators). This field reflects a binary classification of whether an individual is growth (in)determinate.

**Possible values**, cat, factor

- Growth indeterminate – growth continues throughout an individual’s lifespan
- Growth determinate – growth ceases or attenuates to negligibility before the end of an individual’s lifespan
- NF - Growth determination reviewed and inconclusive (no affirmative records found upon review)
- NDY - Growth determination not digitised yet (yet to be evaluated)
- NA – Growth determination not applicable to the subject area

**Usage notes:** Additional classification systems for characterizing growth indeterminacy exist in the literature. Most well recognised is a six-type scheme describing growth and determination by Sebens (1987), which offers detailed characterization of growth patterns. Broad-scale information about growth and age for most species could not be located in the literature, and therefore a simplified schema is used. More resolved classifications might be incorporated into future versions of MOSAIC.

**Additional Information:** NA

**Source Data:** 24 January 2022

**Last updated**: NA

#### A4 *Regeneration*

**Call: -**Morphology$Growth$Regeneration

**Definition:** Capacity for an individual to regenerate any substantial part of its body, including autotomy. Autotomy is defined as “The voluntary severance by an animal of a part of its body (commonly one of its own limbs), usually to escape capture by a predator that has seized that part. The part then regrows.”

**Possible values**, cat., factor

- Regenerative – individuals are capable of regenerating
- Non-regenerative – individuals do not exhibit the capacity to regenerate tissues
- NF - regenerative abilities reviewed and inconclusive (no affirmative records found upon review)
- NDY - regeneration information not digitised yet (yet to be evaluated)
- NA – regenerative abilities not applicable to the subject area

**Usage notes:** There are a number of more resolved schemes for detailing whether regenerating different parts of the body. For particular questions pertaining to the nature of injury, the level of recovery, the role of depredation, and the consequences for reproduction, more detail may be appropriate. This dataset covers the most general applications and initially screens for regenerative capability (from poor recovery of appendages to complete regeneration of limbs).

**Additional Information:** NA

**Source Data:** NA

**Last updated**: 24 January 2022

#### A5 *Sexual Dimorphism*

**Call:** -Morph$Dimorph

**Definition:** An indicator of whether sexual dimorphism is exhibited in the species. Sexual dimorphism is defined as “the occurrence of morphological differences (other than primary sexual characters) that distinguish males from females of a species of organism.” (Oxford Dictionaries of Ecology and Zoology)

**Possible values**, cat., factor

- Sexually Dimorphic – species is sexually dimorphic
- Sexually Monomorphic – species is sexually monomorphic (i.e. non-dimorphic)
- NF - Sexual dimorphism reviewed and inconclusive (no affirmative records found upon review)
- NDY - Sexual dimorphism not digitised yet (yet to be evaluated)
- NA – Sexual dimorphism not applicable to the subject area

**Usage Notes:** NA

**Additional Information:**

**Source Data:** NA

**Last updated**: 24 January 2022

#### B1 *Mating system*

**Call: -**Reproduction$MatingSystem

**Definition:** System of mating; the organization of sexual interactions of individuals within populations based on sex.

**Possible values**, cat., factor

- Monogamy – exclusive mating between one male and one female
- Non-monogamy - Non-monogamy was assigned based on genetic or behavioural evidence
- NF - Mating system reviewed and inconclusive (no affirmative records found upon review)
- NDY - Mating system not digitised yet (yet to be evaluated)
- NA – Mating system not applicable to the subject area

**Usage notes:**

- Metric does not identify size of groups for plural mating systems
- Metric does not identify enforcement mechanisms for different mating systems

**Additional Information:** NA

**Source Data:** NA

**Last updated**: 24 January 2022

#### B2 *Sexual allocation*

**Call: -**Reproduction$Allocation

**Definition:** Indicator of whether a species exhibits hermaphroditism or monoeicieosity. Hermaphrodism is defined as: “An individual that possesses both male and female sex organs; i.e. it is bisexual.” (Oxford Dictionary of Zoology). Monoeciousness is defined as: “Applied to an organism in which separate male and female organs occur on the same individual (e.g. to a plant which bears male and female reproductive structures in the same flower or separate male and female flowers on the same plant, or to a hermaphrodite animal). Some authors restrict the term botanically to plants with separate male and female flowers; plants which bear male and female reproductive organs in the same flower are then called hermaphrodite.”

**Possible values**, cat., factor

- Hermaphroditic – species is hermaphroditic
- Monoecious – species is monoecious
- Dioecious or Gonochorous – species is dioecious (Gonochorous was adopted for animals)
- NF - Hermaphroditism reviewed and inconclusive (no affirmative records found upon review)
- NDY - Hermaphroditism not digitised yet (yet to be evaluated)
- NA – Hermaphroditism is not applicable to the subject area

**Usage notes:**

●

**Additional Information:** NA

**Source Data:** NA

**Last updated**: 24 January 2022

#### B3 *Sequesntial hermaphroditism*

**Call:** Reproduction$SeqHermaph

**Definition:** Indicator of whether there is a sex switch during the organism’s lifespan.

**Possible values**, cat., factor

- Protogynous – species is protogynous: organisms that are female and at some point in their lifespan change sex to male.
- Protandrous – species is protandrous: organisms that are male and at some point in their lifespan change sex to male.
- NF - Protogyny/Protandry reviewed and inconclusive (no affirmative records found upon review)
- NDY - Protogyny/Protandry not digitised yet (yet to be evaluated)
- NA – Protogyny/Protandry is not applicable to the subject area.

**Usage notes:** NA

**Additional Information:** Note that is not rare flowers present protogyny/protandry but this was not considered in Mosaic so far

**Source Data:** NA

**Last updated**: 24 January 2022

#### C1 *Dispersal Capability*

**Call:** -Movement$Dispersal

**Definition:** An indicator for whether or not a species exhibits dispersal behaviour or at any stage in its life cycle. Where dispersing, a categorical description of whether dispersal is natal or breeding or otherwise. Dispersal is defined as “The tendency of an organism to move away, either from its birth site (natal dispersal) or breeding site (breeding dispersal): the opposite of philopatry.” (Oxford Dictionary of Zoology).

**Possible values**: cat., factor

- Dispersing – Exhibits at least one age-/stage-class which disperses; natal or breeding components unknown.
- Natal Dispersal – Permanent dispersal of at least one age-/stage-class
- Breeding Dispersal – Dispersal of adults between breeding attempts in at least one age-/stage-class
- Multi-Dispersal – Both natal and breeding dispersal reported in the species; see DispClasses for more information
- Non-Dispersing – Species observed to have no dispersal traits/behaviour
- NF - dispersal capability reviewed and inconclusive (no affirmative records found upon review)
- NDY – dispersal capability unknown/not yet evaluated
- NA – not applicable

**Usage Notes:** NA

**Additional Information:** NA

**Source Data:** NA

**Last updated**: 24 January 2022

#### C2 *Type of Dispersal*

**Call:** -Movement$TypeDisp

**Definition:** An indication of whether dispersal is a passive (requires assistance) or active (no assistance) event. See DispClasses for more information.

**Possible values**: cat., factor

- Active – organism utilises its own morphology for the dispersal event
- Passive – organism is unable to disperse through their own means and require an external factor
- Active and Passive – organism is able to disperse with assistance but can also use external factors. Active and passive dispersal can occur within the same life stage or can occur in different life stages.
- NF – type of dispersal reviewed and inconclusive (no affirmative records found upon review)
- NDY – type of dispersal unknown/not yet evaluated
- NA – not applicable

**Usage Notes:** NA

**Additional Information:** NA

**Source Data:** NA

**Last updated**: 24 January 2022

#### C3 *Mode of Dispersal*

**Call:** -Movement$ModeDisp

**Definition:** An indicator of the mode of dispersal of the species (plant and animal specific terminology).

**Possible values**: cat., factor

- Motile – the dispersal of animal species without assistance
- Phoretic – the dispersal of animal species by attaching to another animal
- Water currents – the dispersal of animal species by water
- Motile and water currents – animal species that disperse without assistance and by water, both forms of dispersal can occur within the same life stage or can occur in different life stages
- Anemochory – the dispersal of plant seeds by wind
- Anthropochory – the dispersal of plant seeds by humans
- Autochory – the dispersal of plant seeds without assistance from an external vector (e.g., by gravity or ballistic dispersal)
- Hydrochory – the dispersal of plant seeds by water
- Zoochory – the dispersal of plant seeds by animals
- Anemochory and Anthropochory – plant seeds can be dispersed by wind and humans
- Anemochory and Autochory – plant seeds can be dispersed by wind and without the help of an external vector
- Anemochory and Hydrochory – plant seeds can be dispersed by wind and water
- Anemochory and Zoochory – plant seeds can be dispersed by wind and animals
- Autochory and Hydrochory – plant seeds can be dispersed without the help of an external vector and by water
- Autochory and Zoochory – plant seeds can be dispersed without the help of an external vector and by animals
- Hydrochory and Zoochory – plant seeds can be dispersed by water and animals
- Autochory, Anthropochory and Zoochory – plant seeds can be dispersed without the help of an external vector, by humans and by animals
- NF – mode of dispersal reviewed and inconclusive (no affirmative records found upon review)
- NDY – mode of dispersal unknown/not yet evaluated
- NA – not applicable

**Usage Notes:** Plant seed dispersal modes can be subdivided into further categories, however we collated lower order categories into the higher order categories identified here

**Additional Information:** NA

**Source Data:** NA

**Last updated**: 24 January 2022

#### C4 *Dispersal Class*

**Call:** -Movement$DispClass

**Definition:** Age- or stage-classes of the species that are capable of dispersal.

**Possible values**: cat., factor

- Adult – dispersal stage is an individual that has reached maturity, we include sub-adults into this category
- Egg – dispersal stage is a vessel within which an embryo develops and is expelled by an adult allowing for dispersal
- Fertile material – dispersal stage is a part of an individual, or in some cases a complete individual, that contains fertile material (e.g., the alga *Fucus vesiculosus*; detached floating material/individual that contains gametes)
- Gamete – dispersal stage is the reproductive cell not within a vessel
- Juvenile – dispersal stage is an individual that has not reached maturity
- Larval – dispersal in a specific juvenile stage restricted to non-mammal species, species can have multiple larval stages
- Seed – dispersal stage is fertilized, specific to plant species and, in our definition, references seeds and/or fruits that are dispersed
- Sperm – dispersal stage is the male gamete
- Spore – dispersal stage is a single cell that only contains half of the chromosome of the adult, can produce eggs or sperm
- Sporophyte – dispersal stage is a nonsexual phase of a species producing two diploid spores
- Zoospore – dispersal stage is a motile asexual spore
- Zygote – dispersal stage is a fused male and female gamete
- Adult and Juvenile – dispersal stage can be both the adult and juvenile stage
- Egg and Larval – dispersal stage can be both the egg and larval stage
- Gamete and Spore – dispersal stage can be both the gamete and spore
- Zoospore, Sperm and Sporophyte – dispersal stage can be a zoospore, sperm or sporophyte
- NF – dispersal class reviewed and inconclusive (no affirmative records found upon review)
- NDY – dispersal class unknown/not yet evaluated
- NA – not applicable

**Usage Notes:** Dispersal can occur in more than one age- or stage-class, where this occurs it is noted as such within the database

**Additional Information:** NA

**Source Data:** NA

**Last updated**: 24 January 2022

#### C5 *Volancy*

**Call:** -Movement$Volancy

**Definition:** An indicator of whether a species is volant or non-volant (i.e., able to fly or not).

**Possible values**, cat., factor

- Volant – the species is volant (Most Birds (Class Aves), all Bats (Order Chiroptera), and some invertebrate species)
- Non-volant – the species is non-volant
- Semi-volant – the species has gliding abilities (e.g., Gliding lizards (*Draco* spp.); flying squirrels such as the Northern flying squirrel (*Glaucomys sabrinus*); flying fish (Exocoetidae); gliding frogs such as Wallace’s flying frog (*Rhacophorus nigropalmatus*); and gliding ants such as *Cephalotes atratus*).
- NF – volancy reviewed and inconclusive (no affirmative records found upon review)
- NDY – volancy unknown/not yet evaluated
- NA – not applicable

**Usage Notes**: NA

**Additional Information:** NA

**Source Data:** NA

**Last updated**: 24 January 2022

## S2: MOSAIC Trait Primary Search Key Words

### MOSAIC: Keywords used in primary literature search

Taxonomic names (binomial nomenclature) were used in connection with the field-specific terms in the search of primary literature in identifying the records of interest. The below include names that were searched in review of the primary literature.

Biomass—

NA (database only)

Height—

NA (database only)

Growth determination—

Sources: *Web of Science; Google Scholar; Scopus*
Terms: Growth determinat*, growth determination, growth determinate, growth indetermination, growth

Regeneration—

Sources: *Web of Science; Google Scholar; Scopus*
Terms: Regenerat*, regeneration, regenerate, rejuvenation

Sexual dimorphism—

Sources: *Web of Science; Google Scholar; Scopus*
Terms: Dimorphic

Mating system—

Sources: *Web of Science; Google Scholar; Scopus*
Terms: Monogamy, monogamous, polygyny, polygynous, polyandry, polyandrous, mating system

Hermaphrodism—

Mating system—

Sources: *Web of Science; Google Scholar; Scopus*
Terms: Hermaphrod*, hermaphroditic, hermaphrodism, hermaphrodite, gonochoric, gynochorous, monoecious, dioecious, sexual differentiation,

Sequential hermaphrodism—

Sources: *Web of Science; Google Scholar; Scopus*
Terms: Sequential hermaphrodism, protandry, protogyny, protogynous hermaphroditism, protandrous hermaphroditism

Dispersal capacity—

Sources: *Web of Science; Google Scholar; Scopus*
Terms: Dispers*, dispersal, dispersing, dispersal capacity, dispersal capability

Type of dispersal—

Sources: Web of Science; Google Scholar; Scopus
Terms: Dispers*, active, passive

Mode of dispersal—

Sources: *Web of Science; Google Scholar; Scopus*
Terms: Dispers*, motile, phoretic, currents, anemochory, anthropochory, autochory, hydrochory, zoochory, anthropochory

Dispersal classes—

Sources: *Web of Science; Google Scholar; Scopus*
Terms: Dispers*, adult, jouvenile, egg, fertile, seed, sperm, spore, sporophyte, zoospore, zygote, gamete

Volancy—

Sources: *Web of Science; Google Scholar; Scopus*
Terms: Volant, non-volant, flight, flying, flightless, non-flying, winged, glide, gliding, ground

Aquatic habitat dependency—

Sources: *Web of Science; Google Scholar; Scopus*
Terms: Anadromous, catandromous, estuarine, brackish, lotic, lentic, lemnitic, littoral, palagic, marine, freshwater, saltwater, sea, ocean.

## S3. Databases Searched

### MOSAIC: Databases Reviewed to Date for the MOSAIC Database

Type. Database Name. DOIs.

1. **Plants**. Botanical Information and Ecology Network (BIEN) (doi: 10.7287/peerj.preprints.2615v2)
2. **Plants**. TRY: Global Plant Trait Database (TRY) (doi: 10.1111/j.1365-2486.2011.02451.x)
3. **Phylogeny.** Open Tree of Life: A synthesis of phylogeny and taxonomy into a comprehensive tree of life (doi: 10.1073/pnas.1423041112)
4. **Vertebrates**. Amniote: An amniote life history database to perform comparative analysis with birds, mammals, and reptiles (doi: 10.1890/15-0846R.1)
5. **Amphibians**. AmphiBIO: a global database for amphibian ecological traits. (doi:L 10.1038/sdata.2017.123)
6. **Mammals**. PanTHERIA: a species-level database of life history, ecology, and geography of extant and recently extinct mammals (doi: 10.1890/08-1494.1)
7. **Animals**. AnAge Database of Animal Ageing and Longevity. (doi: 10.1111/j.1474-9726.2008.00442.x)
8. **General (Sex)**. Tree of Sex: A database of sexual systems (10.1038/sdata.2014.15)
9. **General (Variable)**. Open Trait Network Databases (https://opentraits.org/)

## S4: Vignettes

### Navigating MOSAIC

#### Vignette #1 - Navigating MOSAIC

Welcome to the **MOSAIC database**, a database of functional traits for comparative demography. The database, user, guide, and additional information can be found and http://mosaicdatabase.web.ox.ac.uk.

MOSAIC is a database that aggregates existing databases and adds new records for functional traits that currently do not have a database established. In this vignette, we will show you how to download the dataset, search records, and relate MOSAIC records with COMADRE, COMPADRE, and PADRINO databases.

`Optional clearance of working space.`

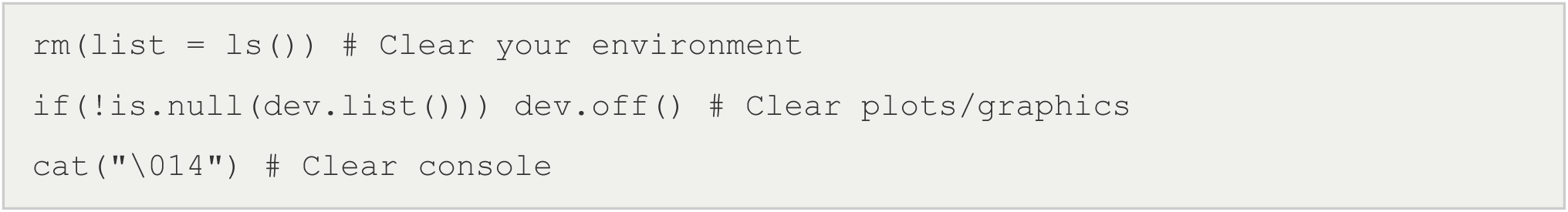

`Downloading MOSAIC`

MOSAIC can be downloaded as an S4 object by running the below code in R. S4 data objects in R are an object oriented system in the R language that allow control of constituent data fields Similar to S3 objects (which use the “$” operator). S4 are comprised of objects that can be searched with the “@” operator or slots, discussed in more detail below.

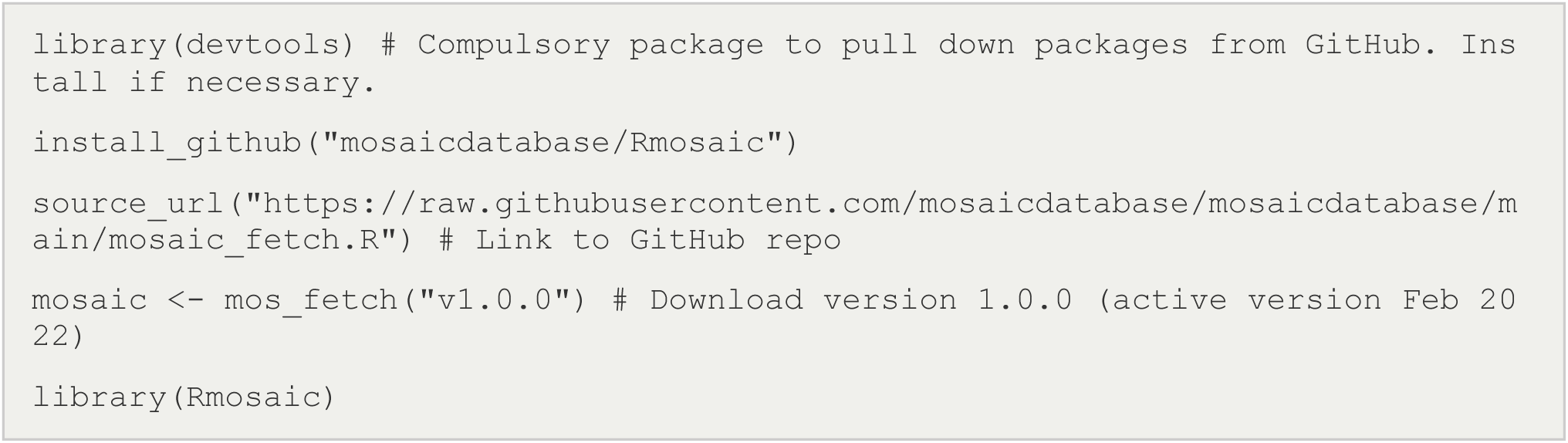

`Basics of manually navigating MOSAIC`

Mosaic traits can be searched using the “@” operator. Attribute names searched this way are analogous to the columns of a dataframe in a relational database structure.

Once downloaded, you should be able to type statements mosaicdatabase@[insertfield] (where [insertfield] is a particular trait). If you are working in Rstudio, after the “@” a drop-down of the slots (traits) should autopopulate.

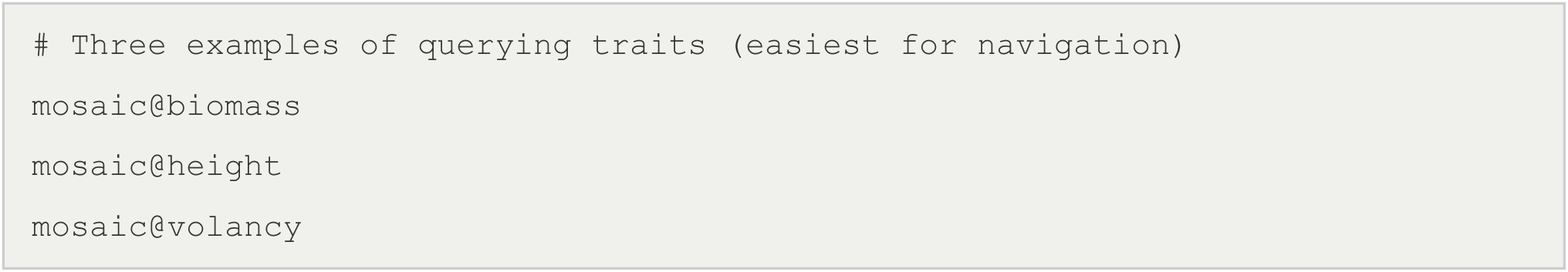

`The species corresponding with each index can be queried by prompting:`

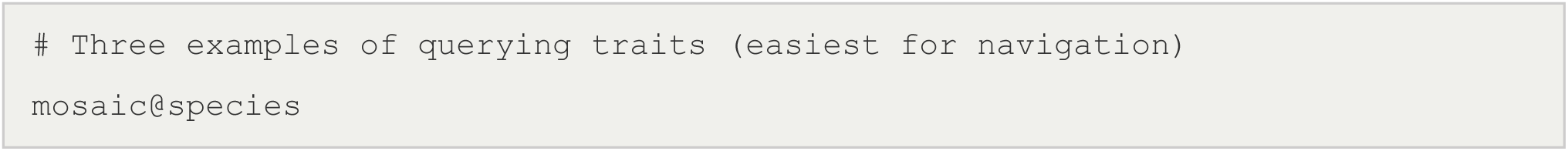

Data in mosaic can also be access using slots. Slots are the recommended mode of searching the database - though it has the disadvantage of not enabling the autopopulation of the attributes contained in the database (traits must be spelled out manually).

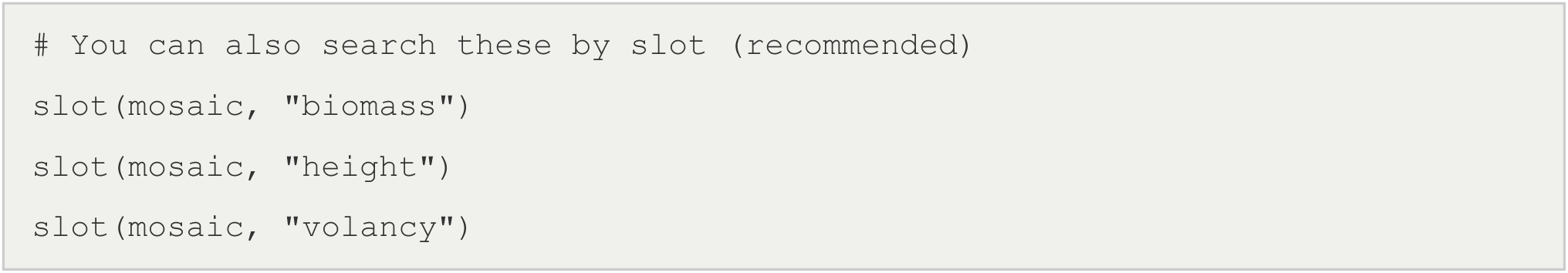

Within each trait object, there are eight fields in mosaic. The first of field is called “values” and contains the data. Values are unitless values (either numeric or factorial) that are reported in units described in the **Mosaic User Guide** http://mosaicdatabase.web.ox.ac.uk/user-guide. The metadata is organised into additional attributes, reflecting the individual elements of the metadata for a given record, including the authors, journal, year of publication, databases from which data are sourced (if applicable)

<u>mosaic@metaTaxa</u> maps the complete taxanomic classification structure of a species - from Kingdom to species for taxonomic clustering.

`The other six attributes - “author”, “year”, “journal,”doi“,”database“, and”mosaic" - are metadata corresponding with each value record. For instance:`

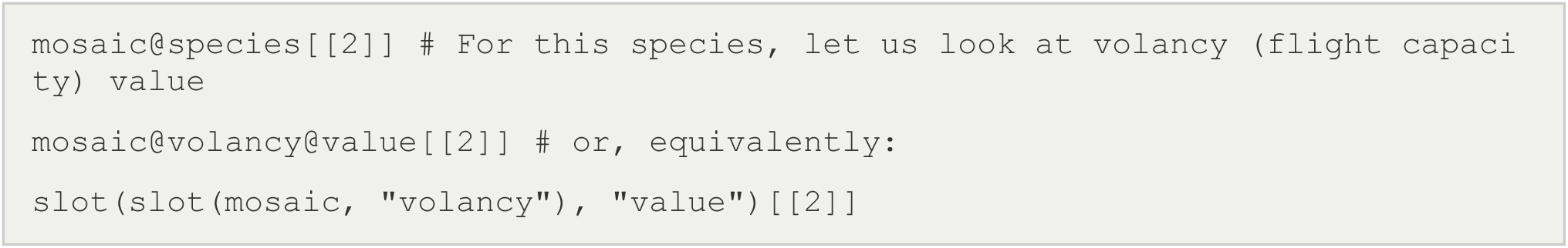

`Corresponds with the following metadata`

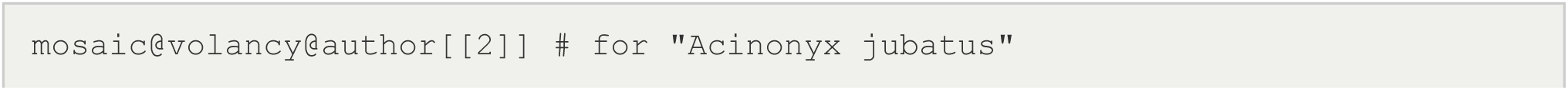
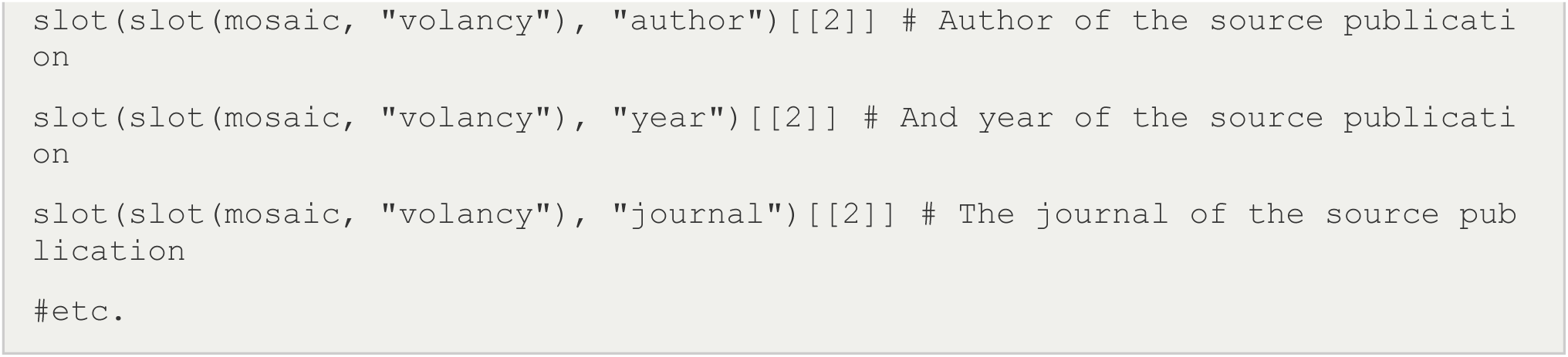

`Using MOSAIC functions to quickly access files`

`A series of convenience functions can be sourced from the MOSAIC GitHub page to facilitate navigating and working with the mosaic database that can be accessed by running the follwing script.`

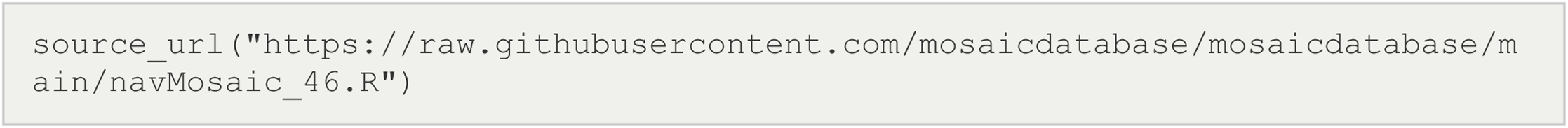

`Below we highlight some of the basic queries for which the mosaic functions can assist.`

`Is a species included in Mosaic?`

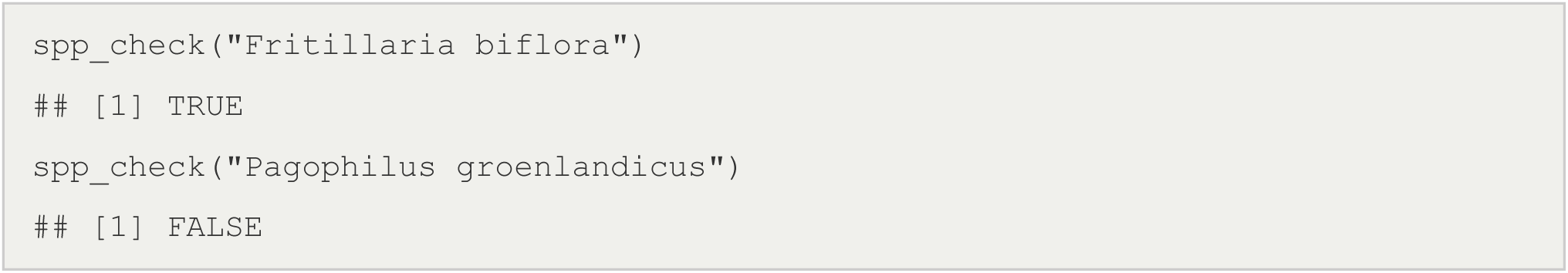

`Can I see all records for a given trait?`

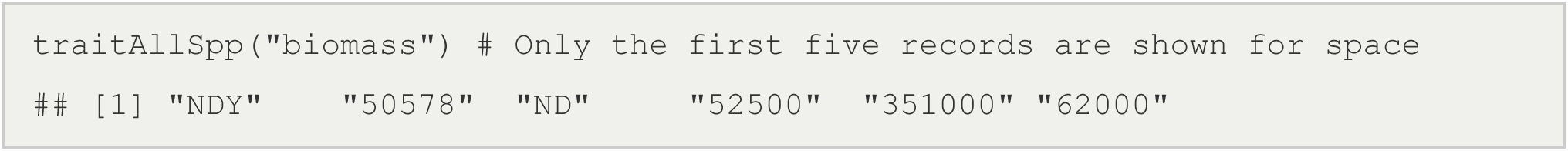

`Can I see an overview of all records for a given species?`

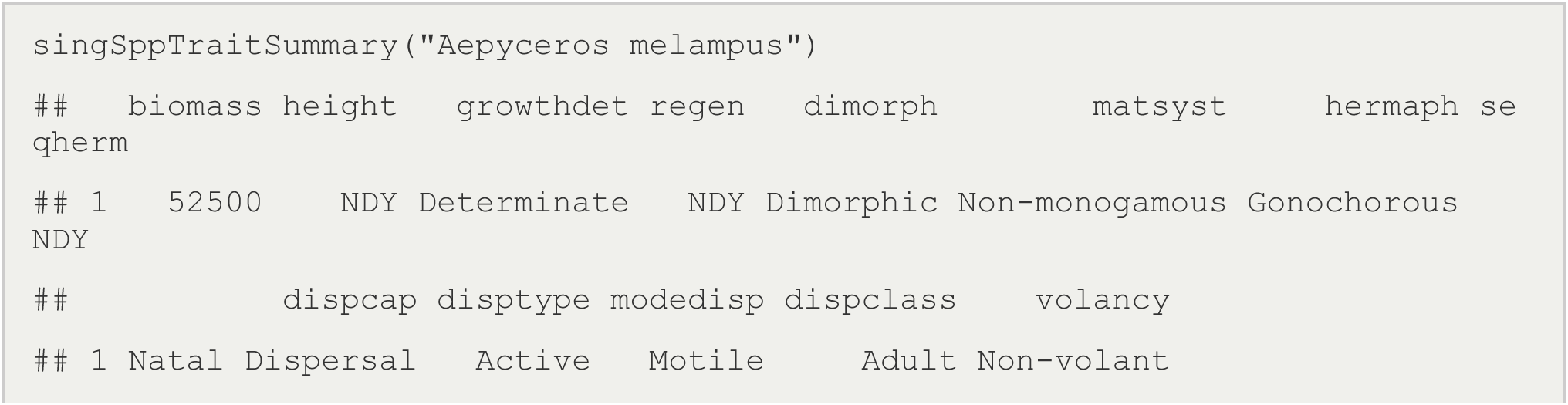
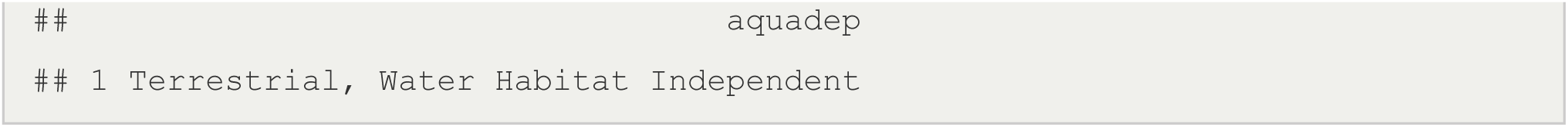

`Can I see all records for more than one species?`

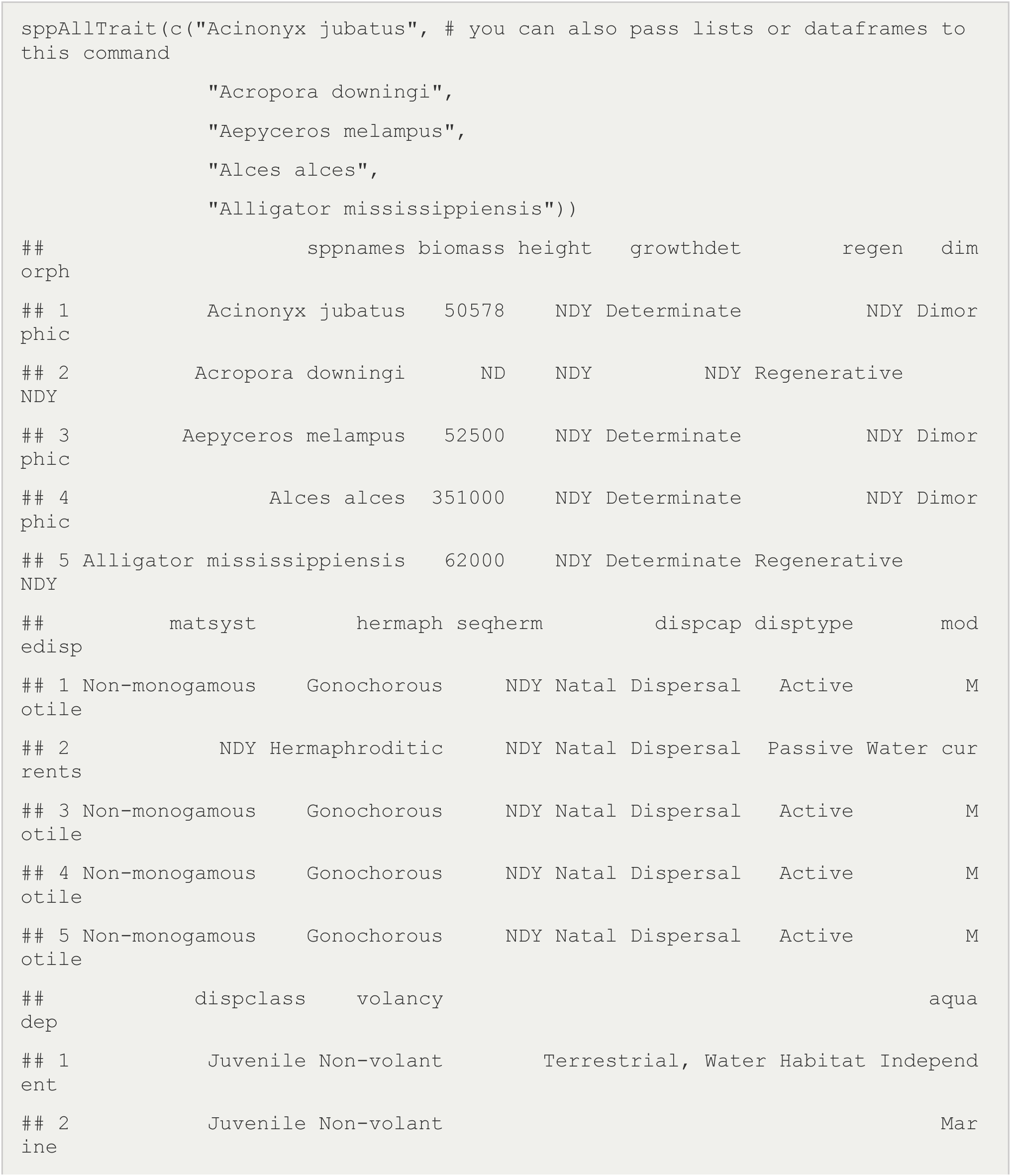
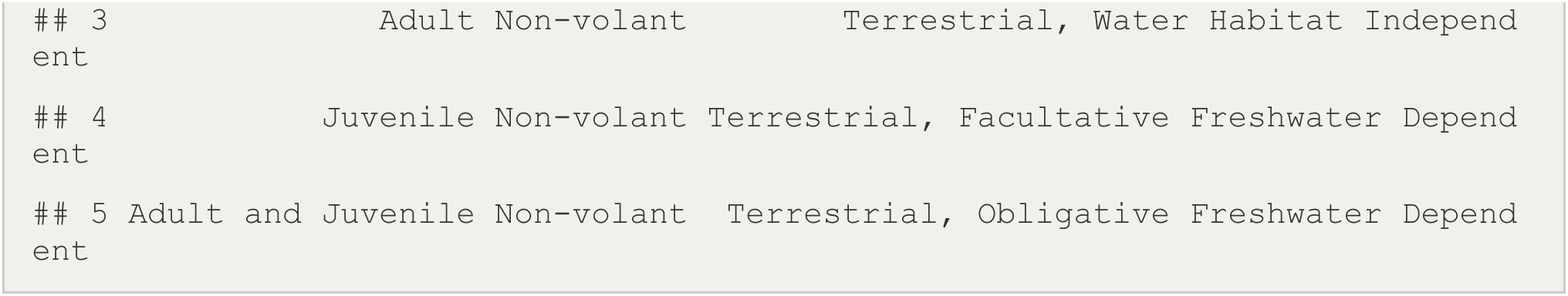

`Can I get a breakdown of counts/frequency of trait values?`

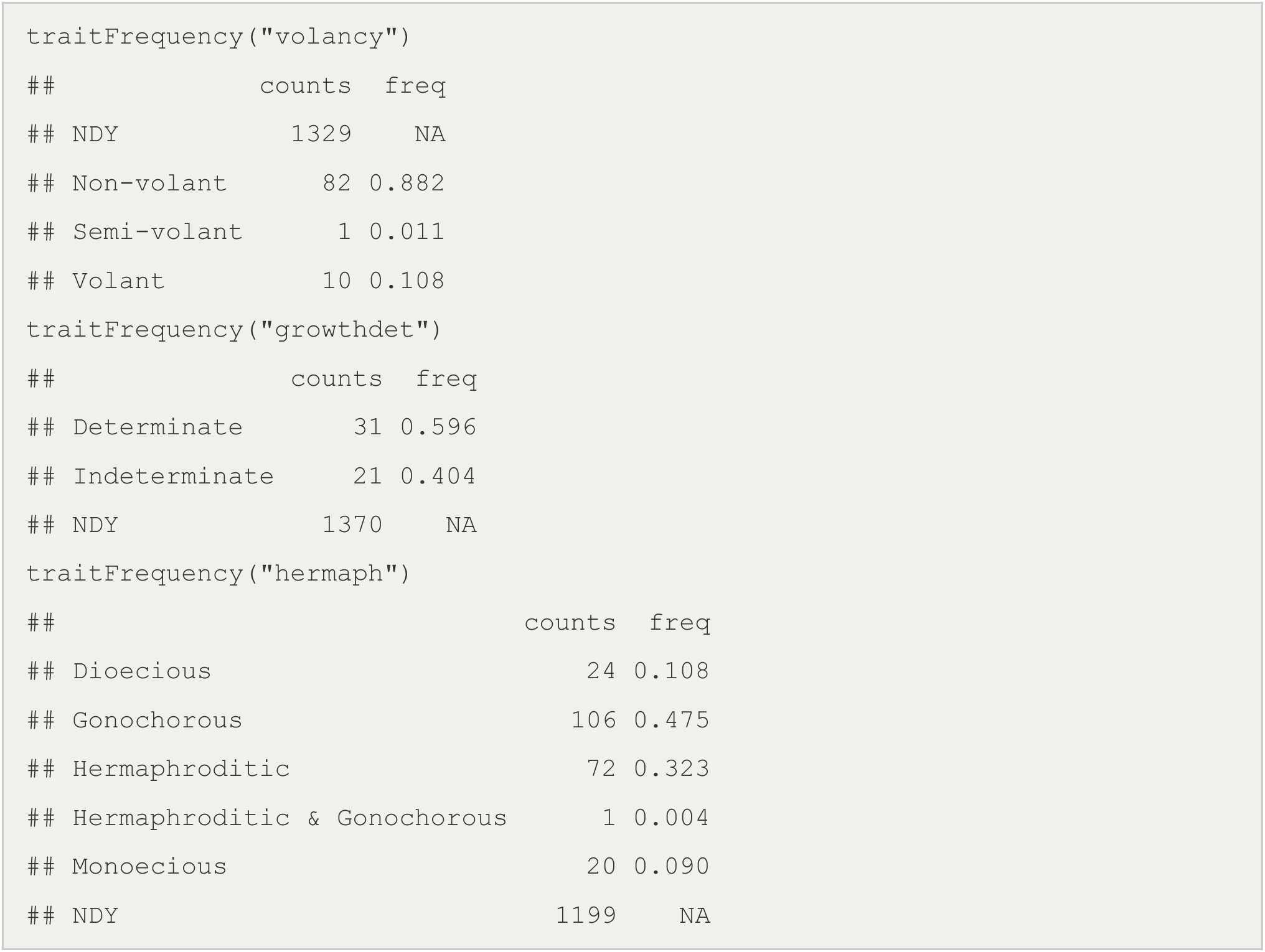

`Can I get all metadata for one or more traits?`

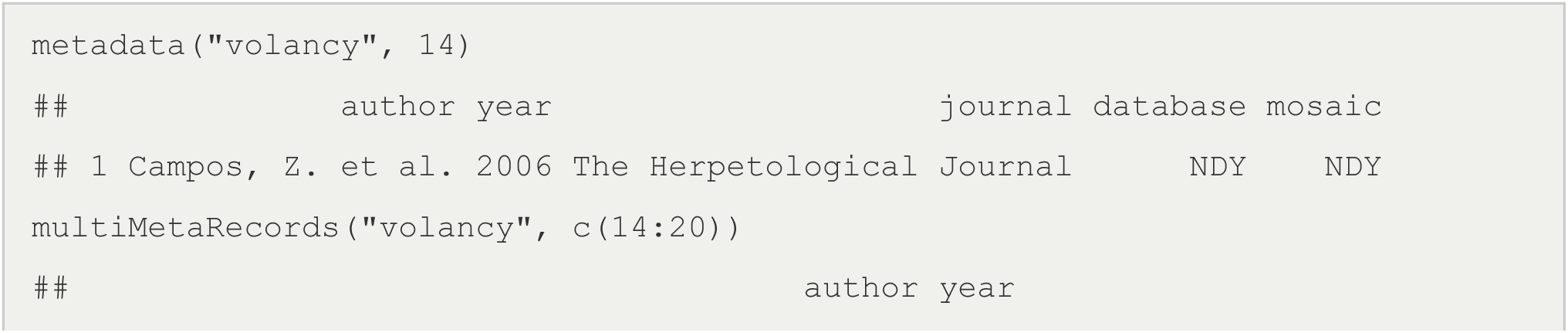
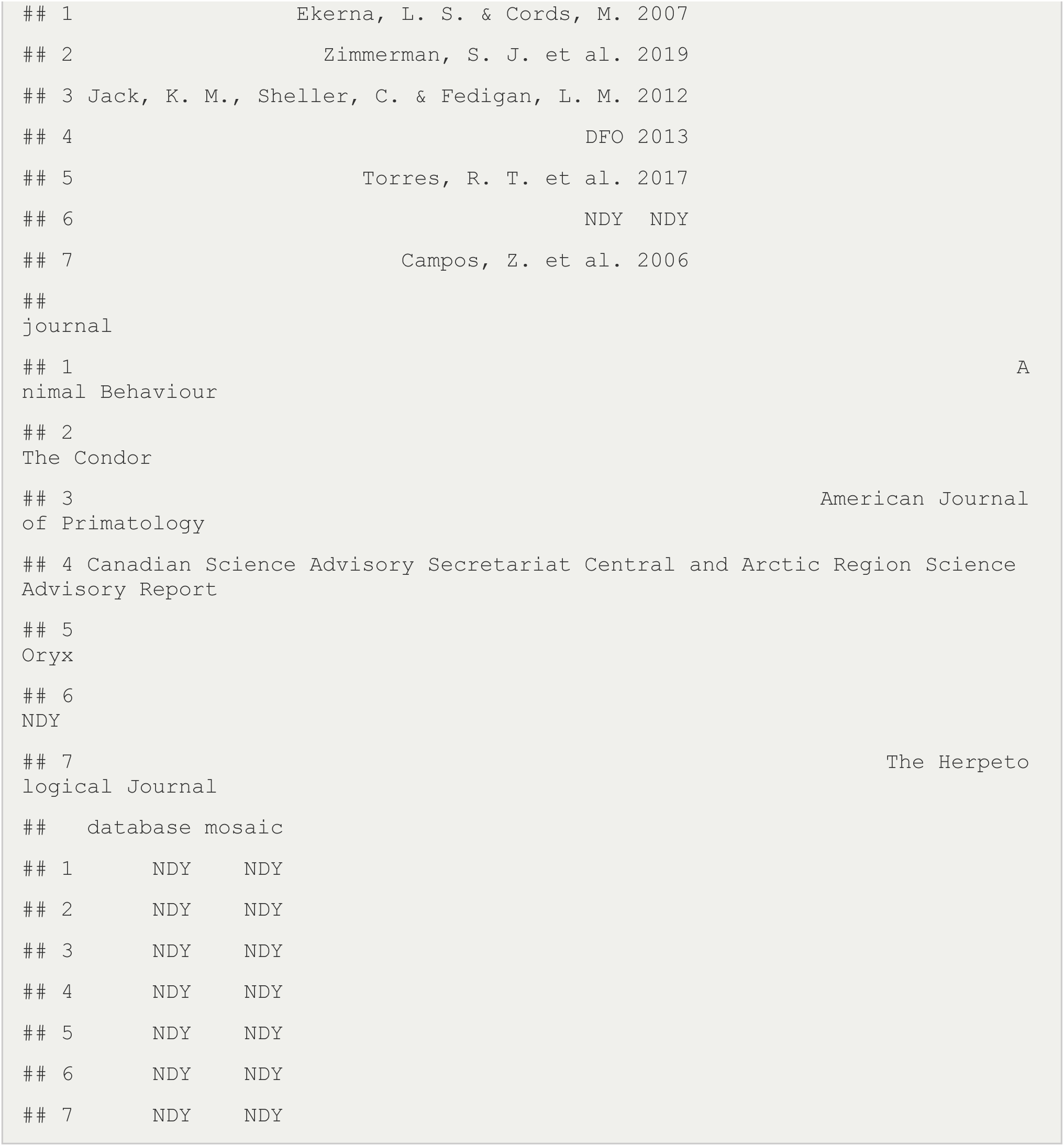

### Tidyverse + MOSAIC

http://mosaicdatabase.web.ox.ac.uk

`Updated 14 March 2022`

#### Vignette #2 – Tidyverse + MOSAIC

MOSAIC was built for use and integration with other datasets for comparative demography. Demographers are often interested in how traits scale across different taxonomic levels. The Tidyverse https://www.tidyverse.org/ collection of R packages (Dplyr in particular) are commonly used by ecologists to simplify workflows that subset and manipulate data. Here, we walk throught the basics of how Tidy tools can be use with the MOSAIC database to quickly isolate specific groups of organism or classes of records.

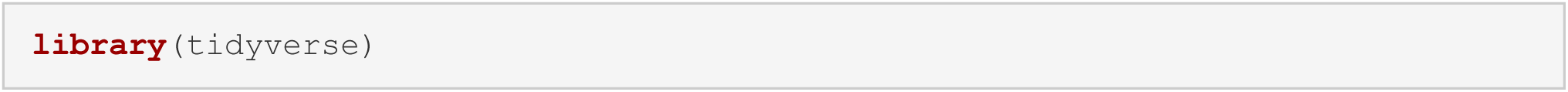

#### Accessing MOSAIC

Download MOSAIC from the mosaic portal. For more information on the basics of downloading MOSAIC and navigating the data structure, see: <u>Vignette #1</u>:

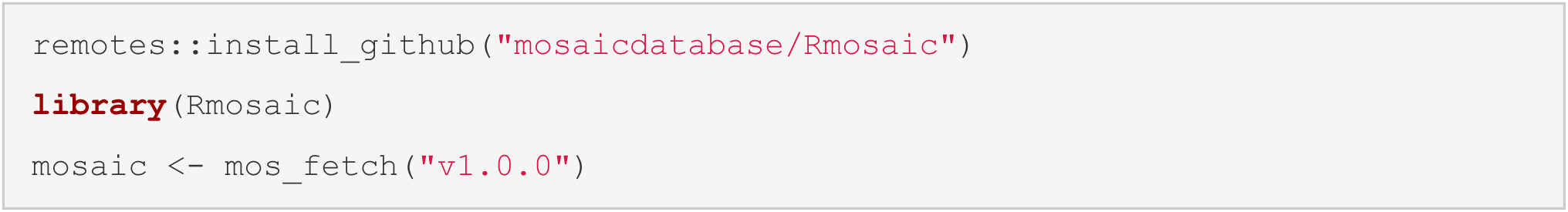

#### Using Tidyverse with MOSAIC

Tidyverse allows for the easily filtering of outliers and NAs, and the isolation of records to particular groups of taxa in MOSAIC. For example, one can isolate the differences in biomass between birds and mammals and see how the decomposition of variance in biomass between the groups informs the representation of biomass values in MOSAIC

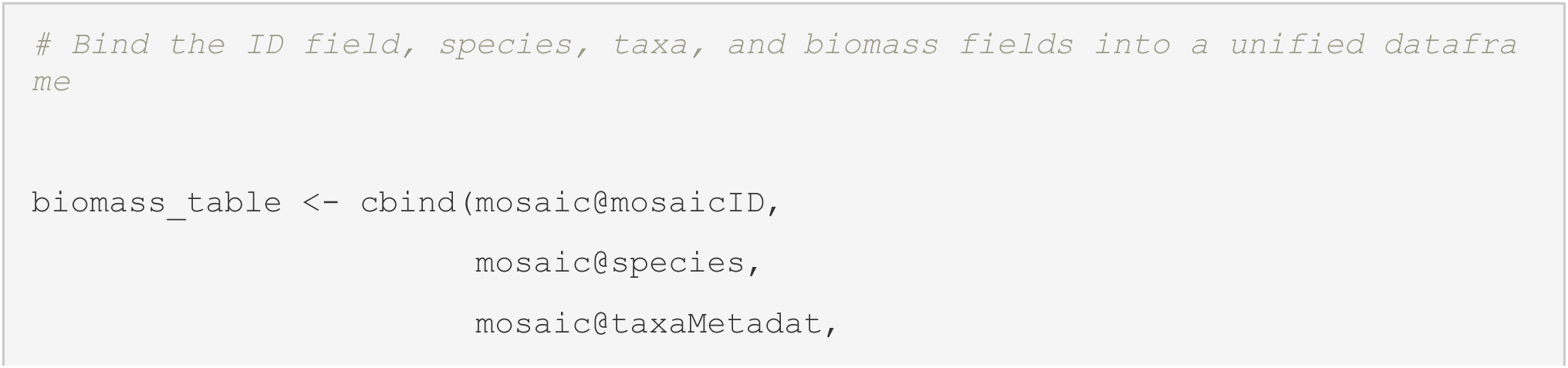
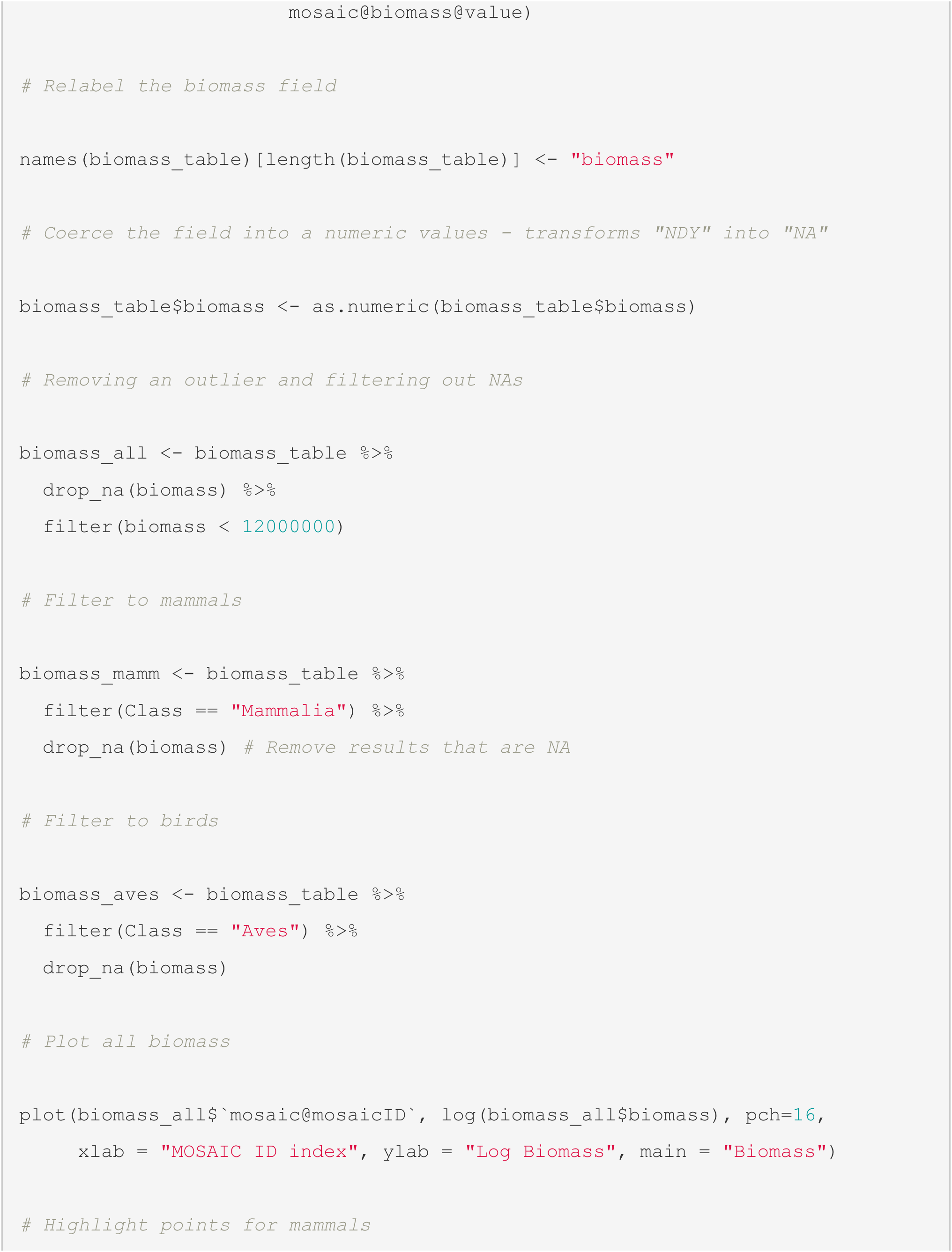
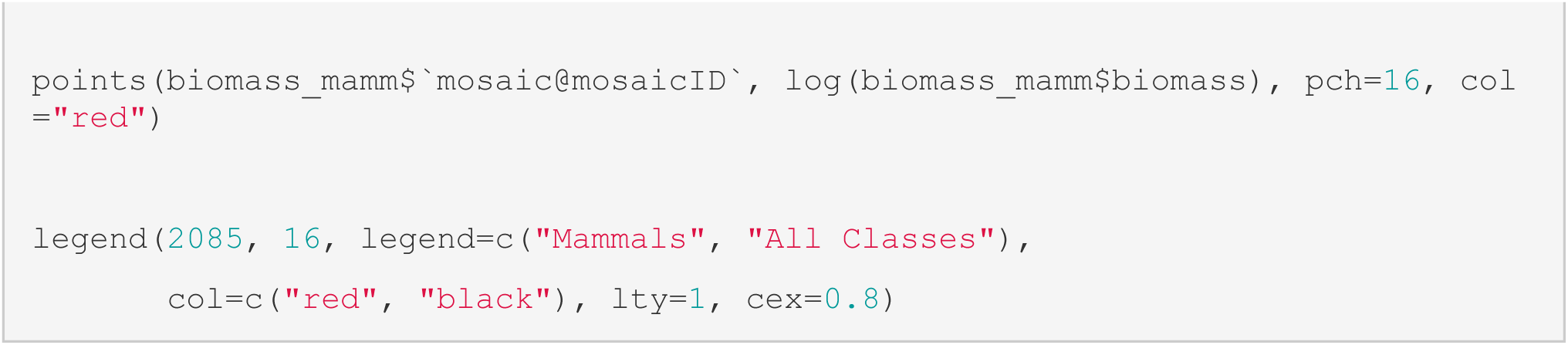

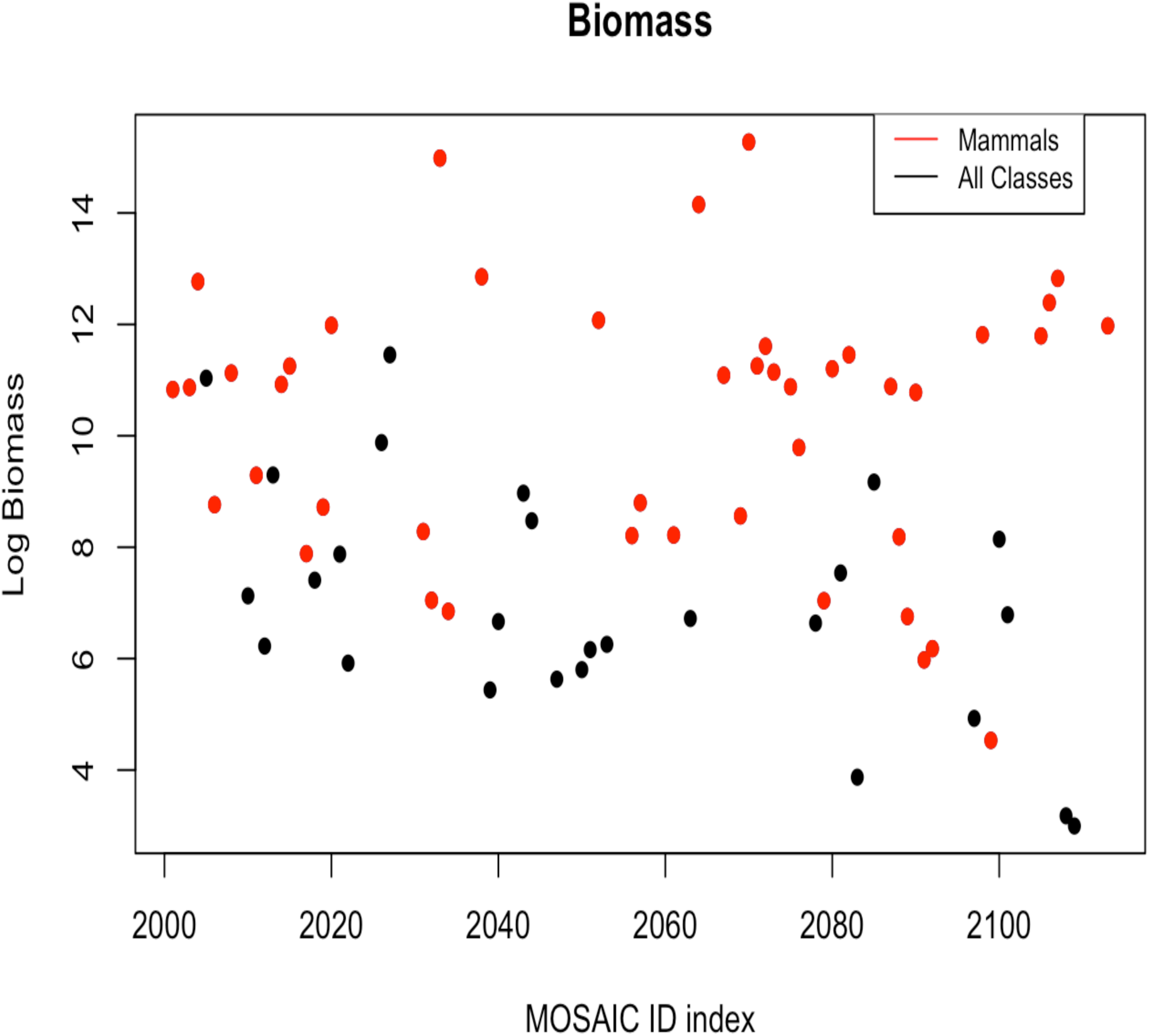

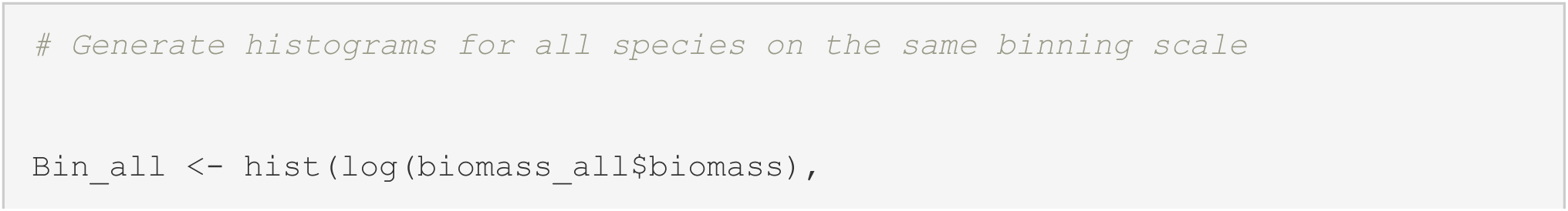
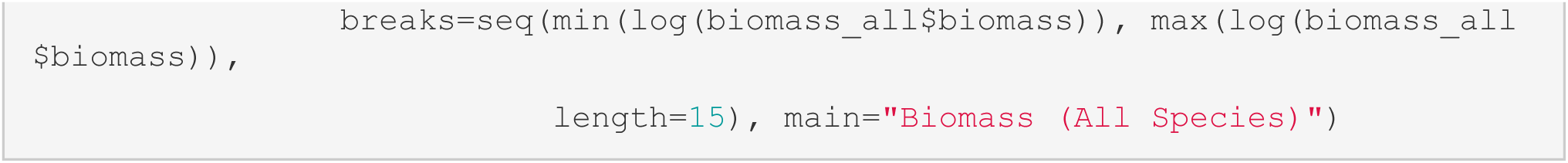

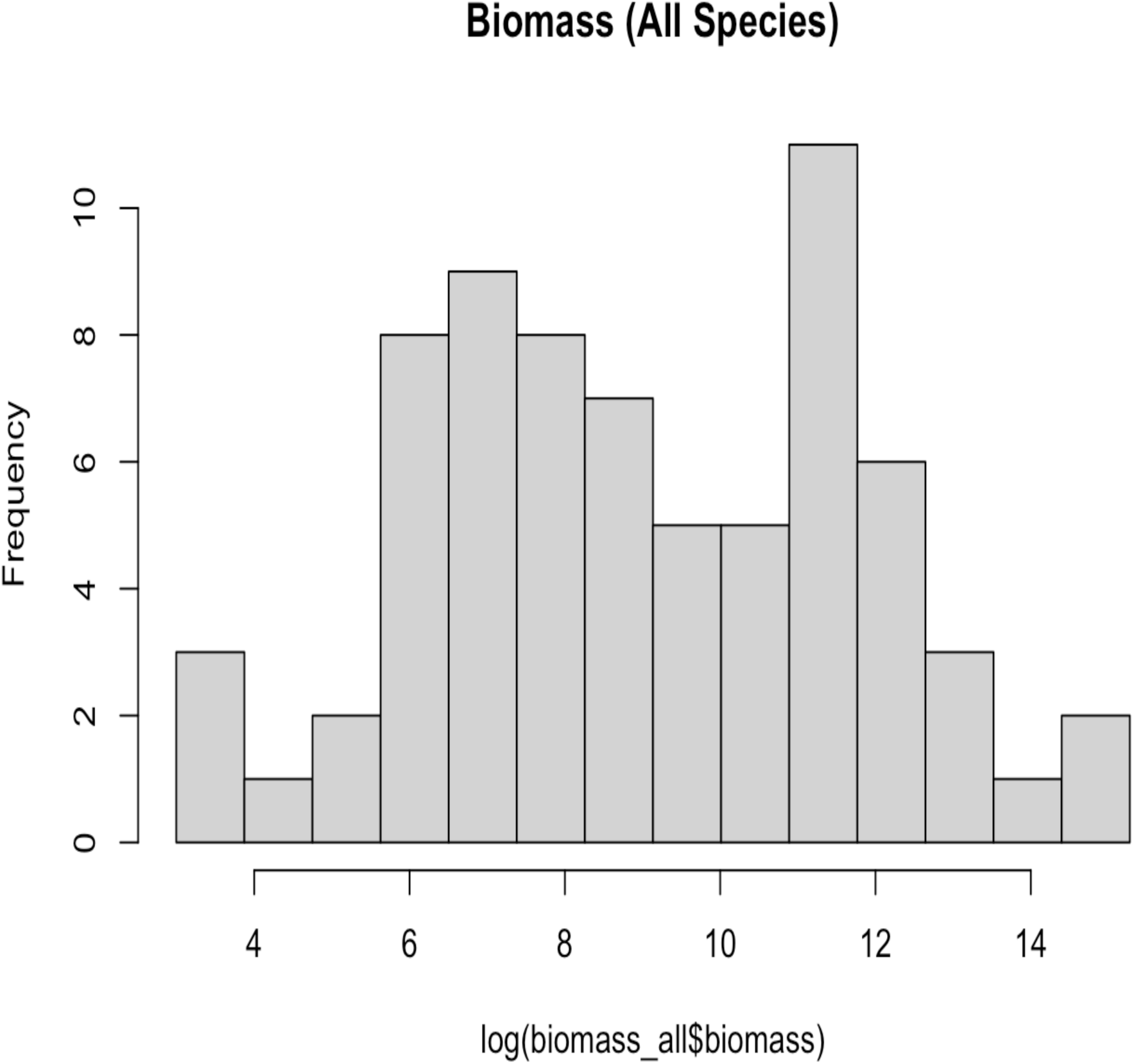

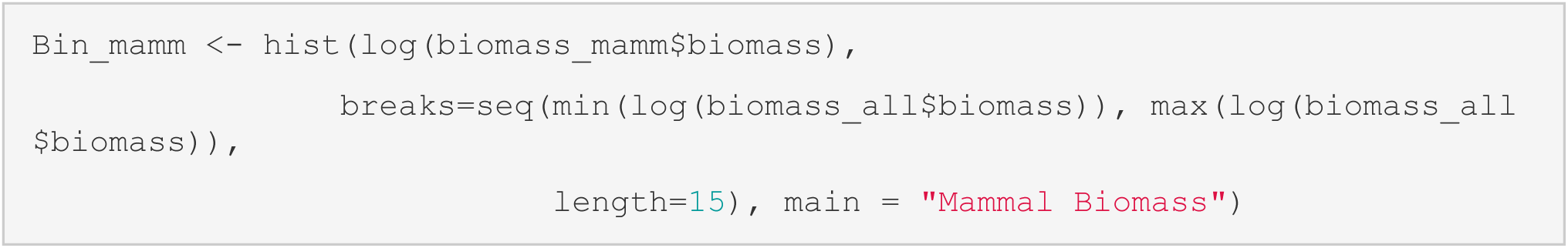

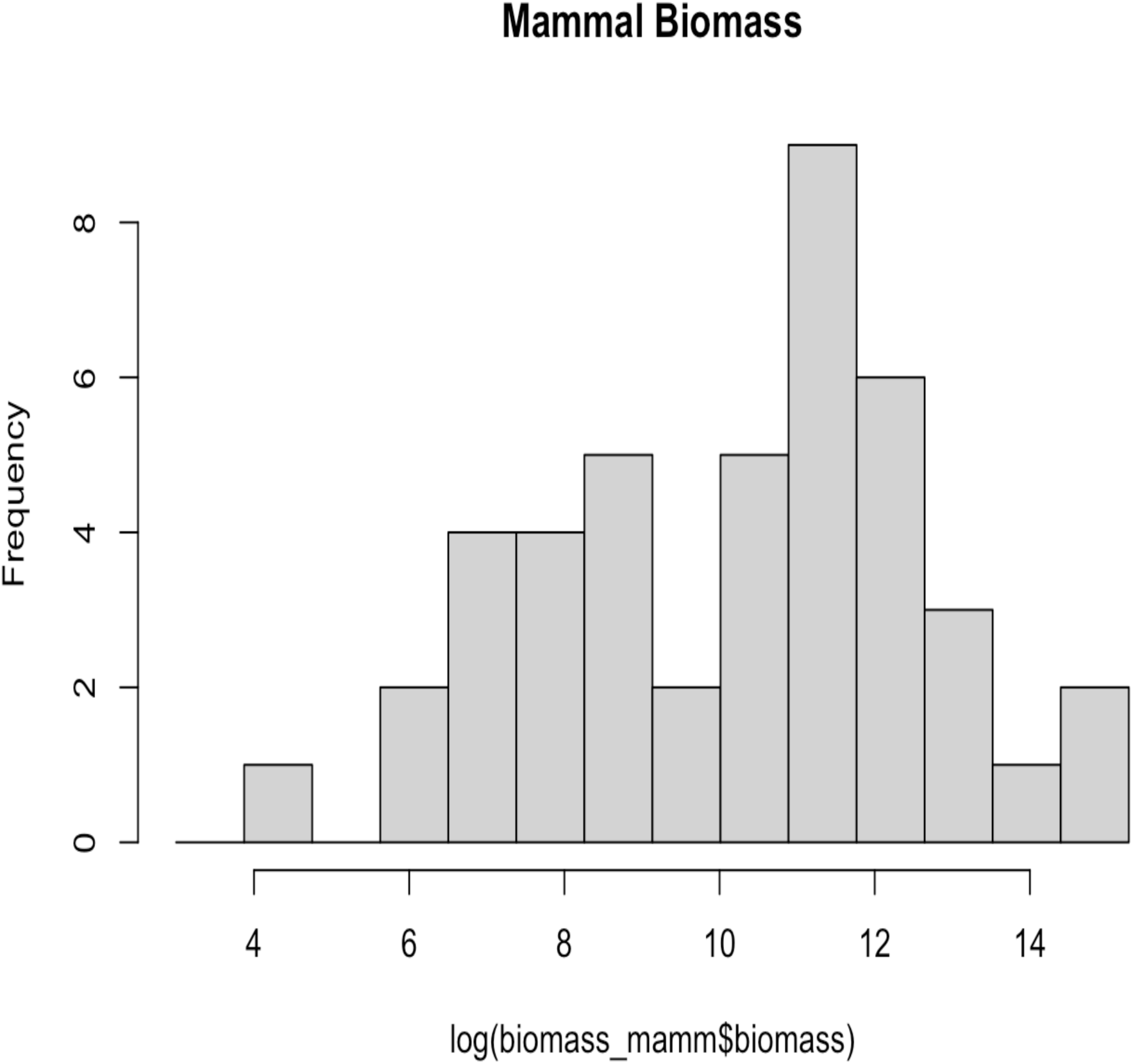

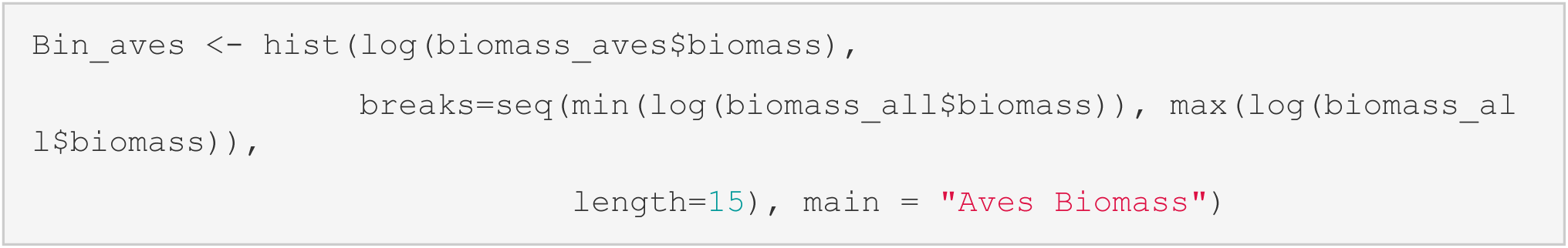

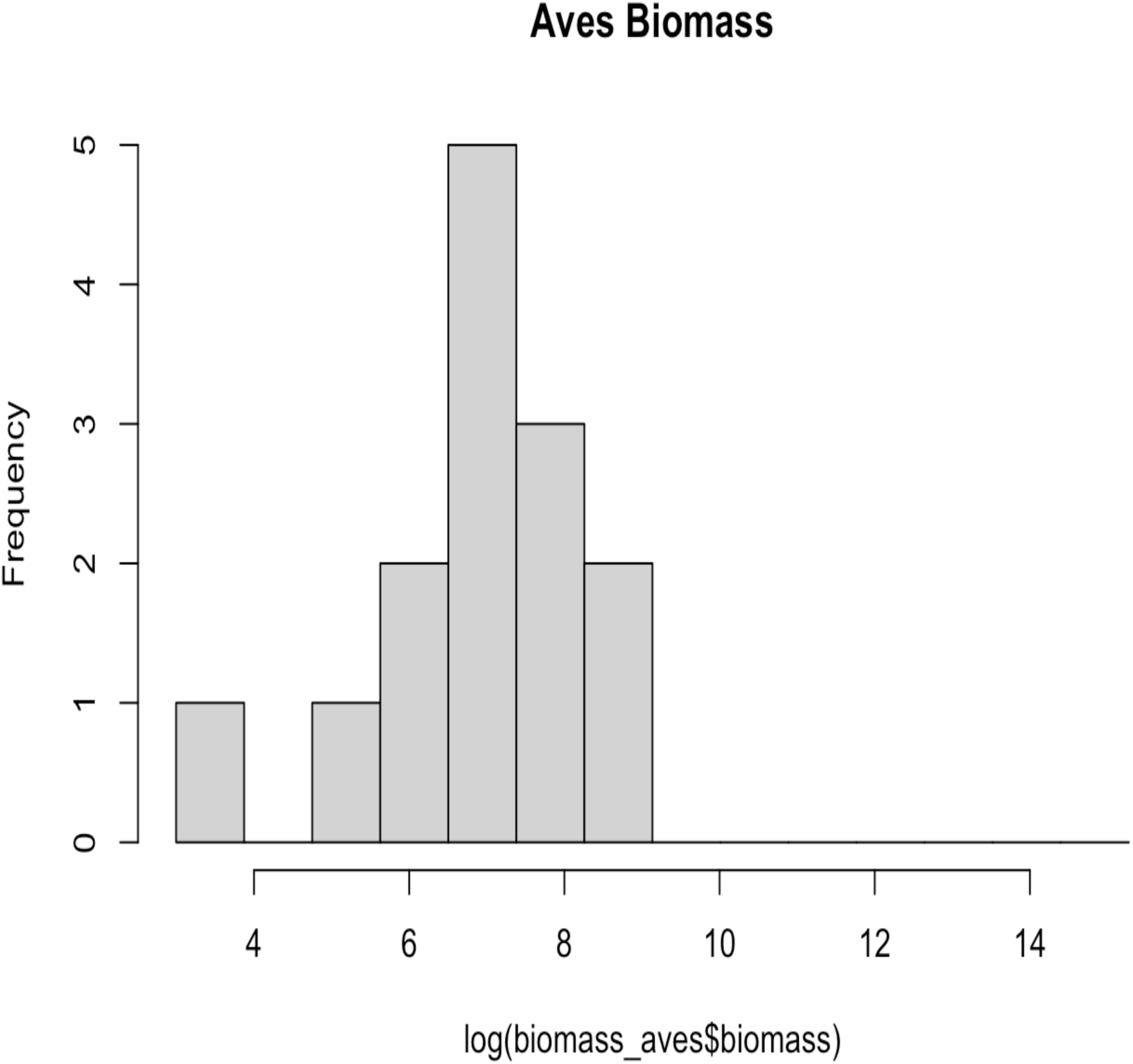

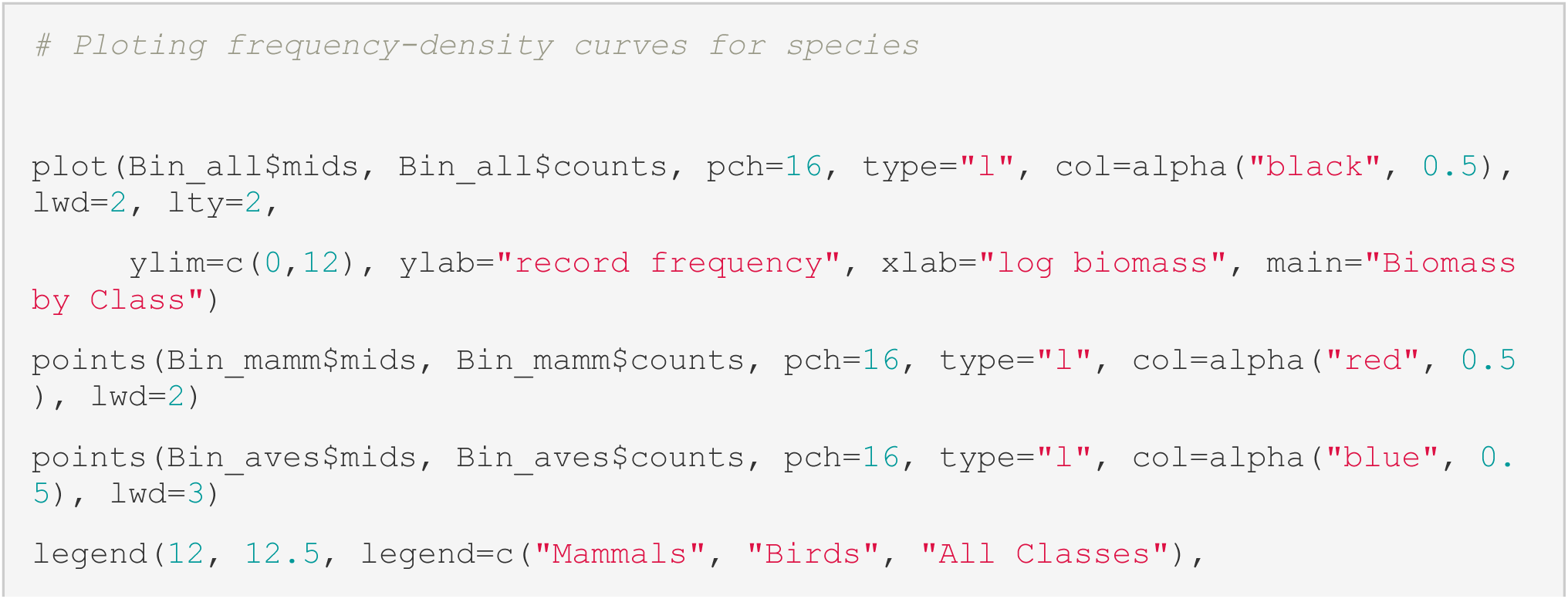
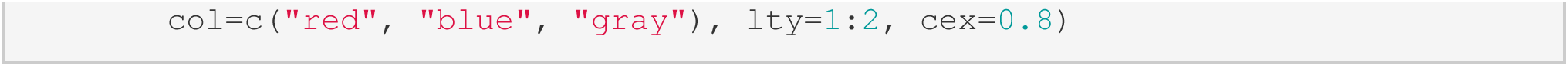

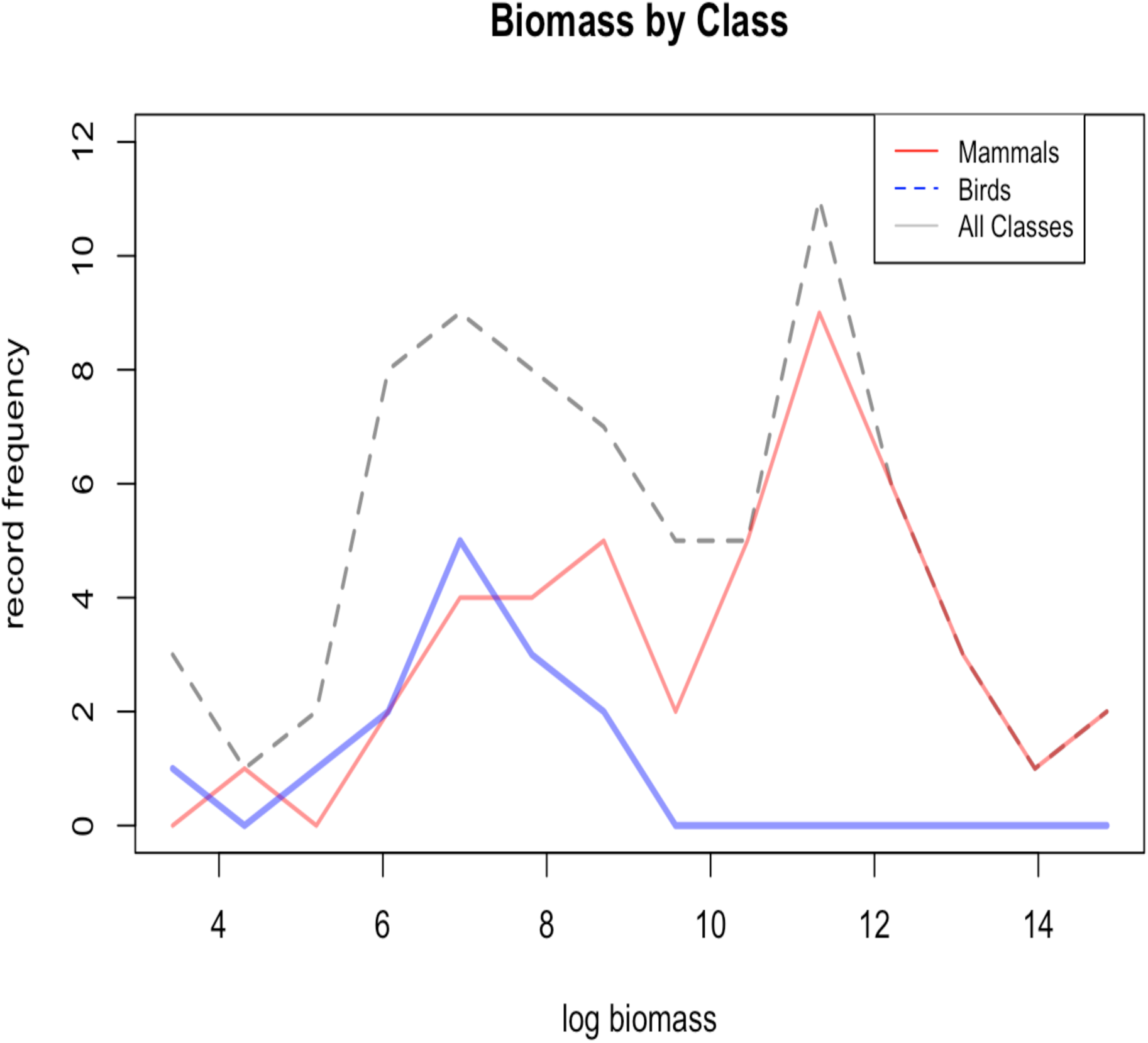

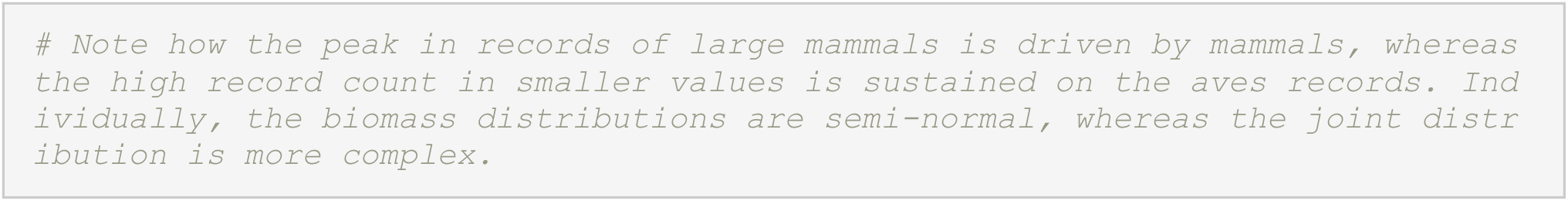

`Alternatively, the same data could be plotted by group with boxplots, illustrating the same sotry.`

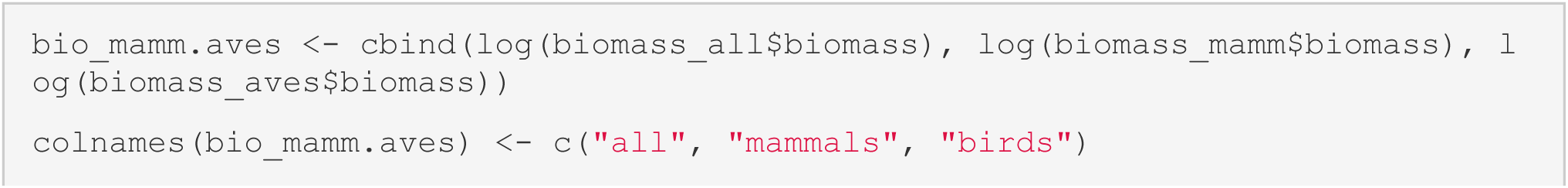
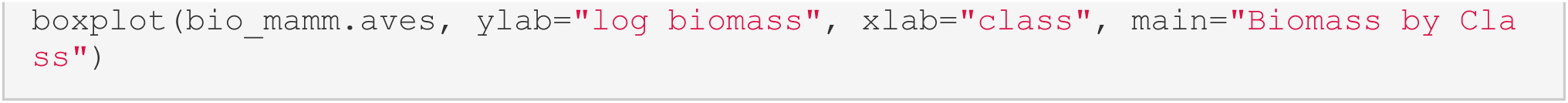

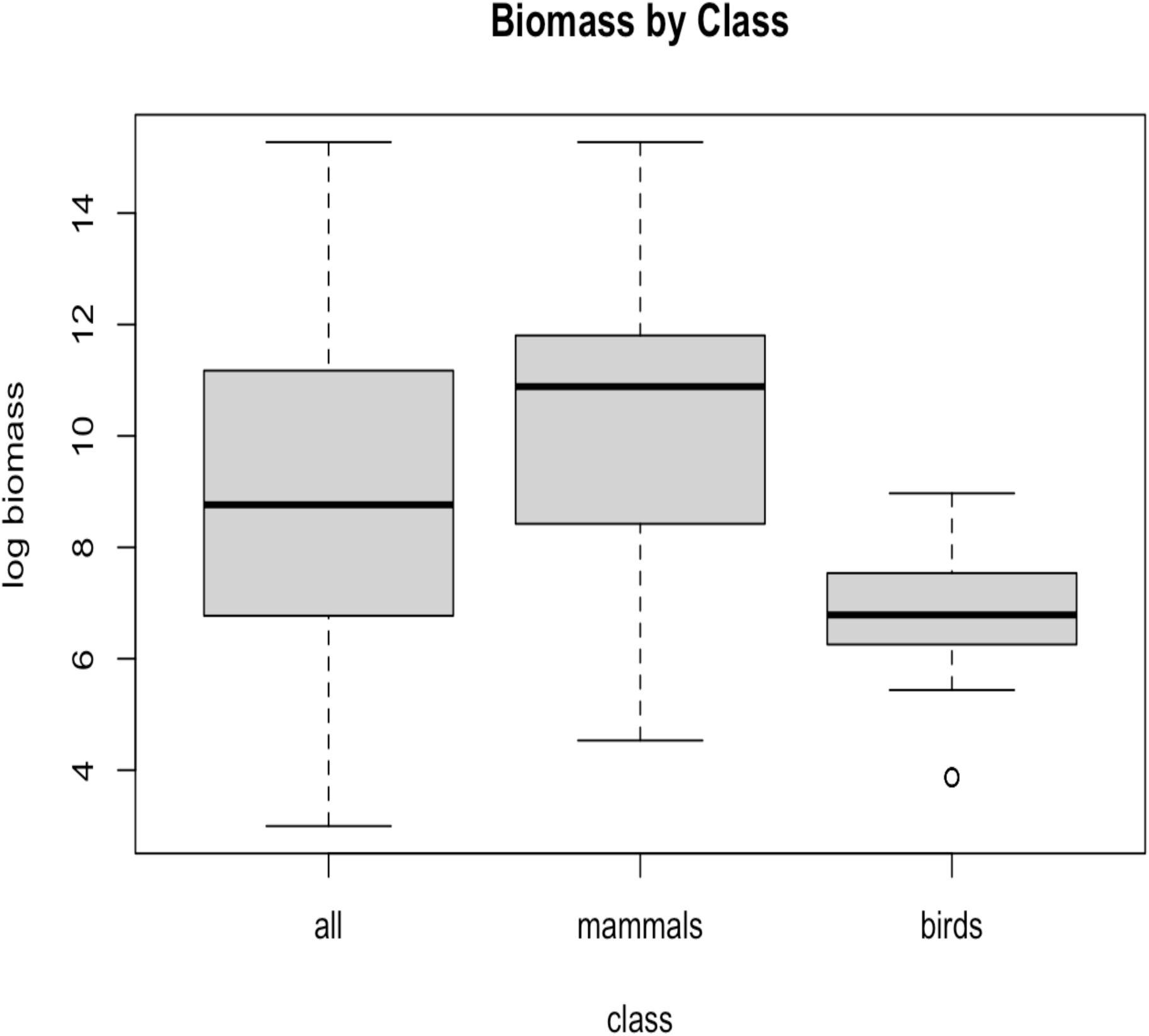

`Factor variables can similarly be parsed by class.`

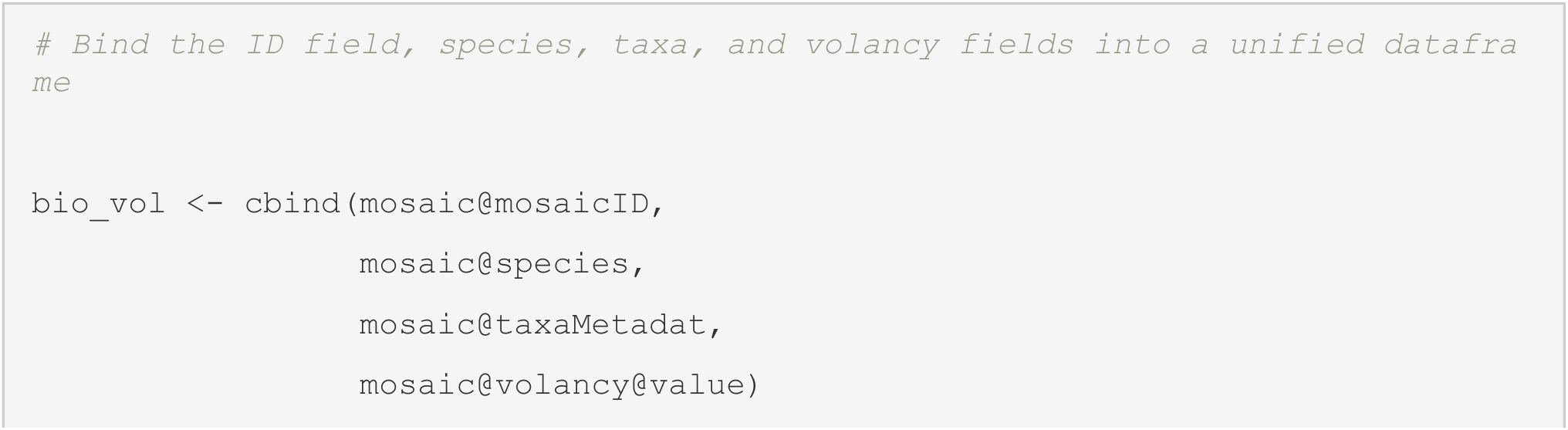
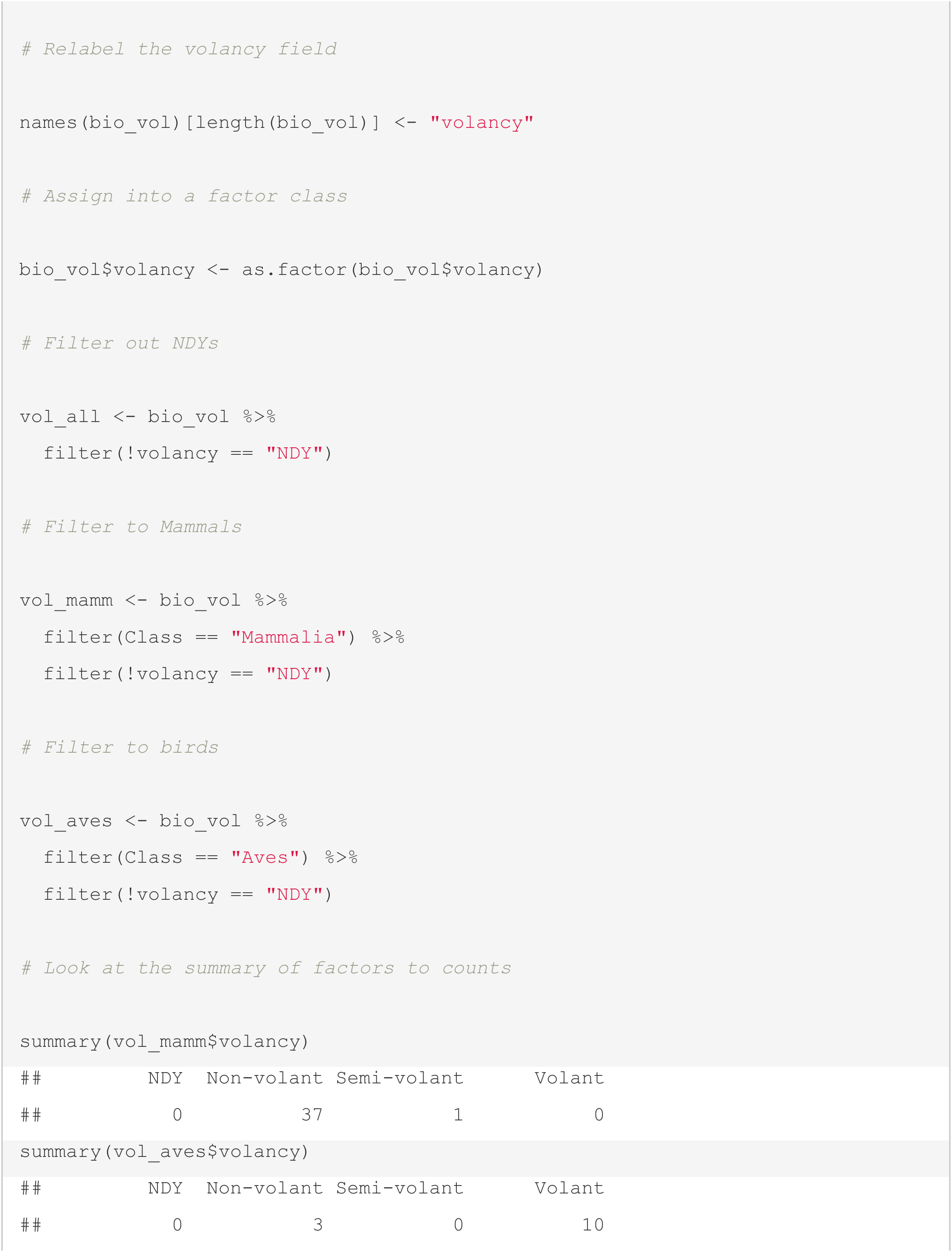
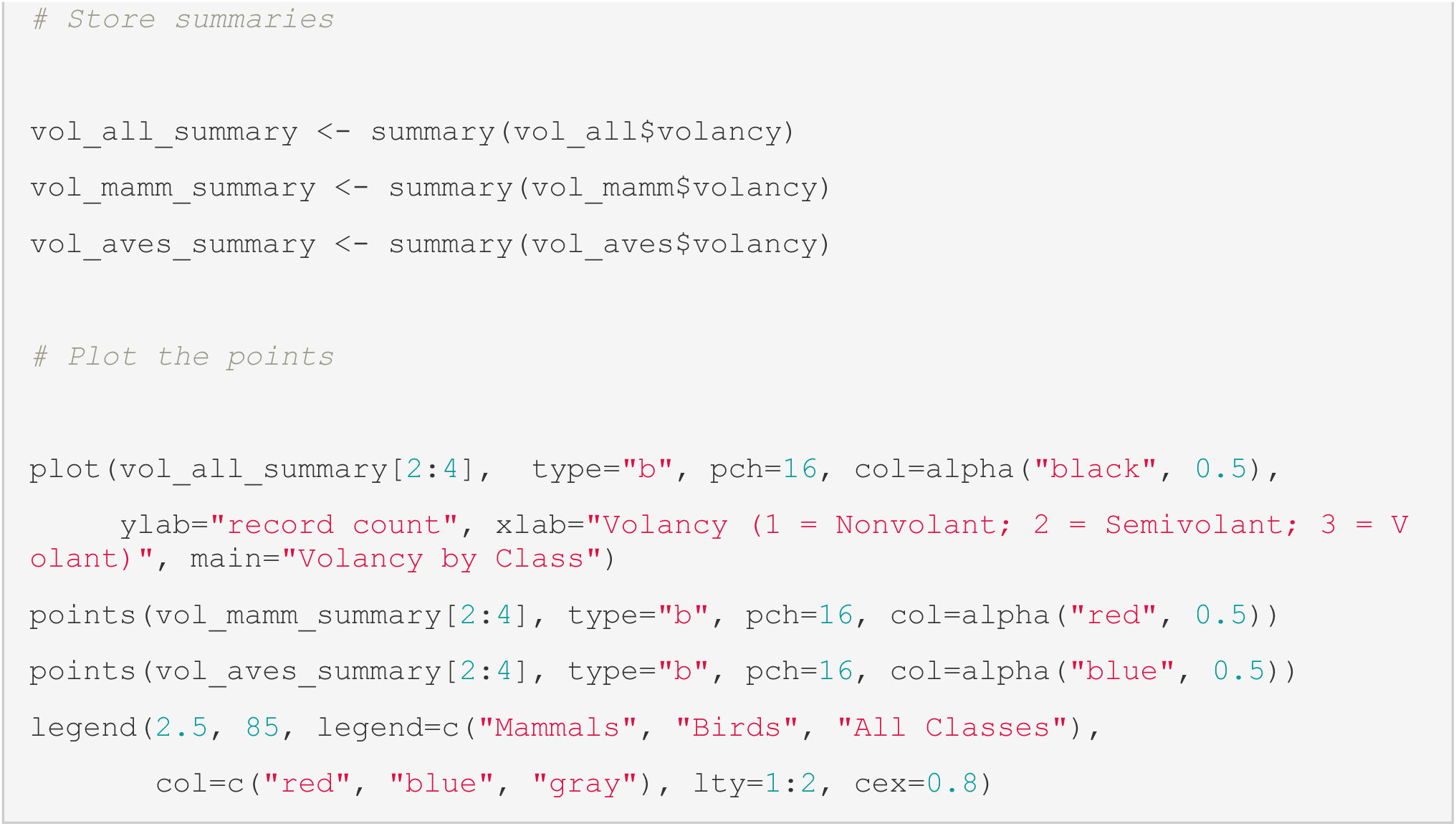

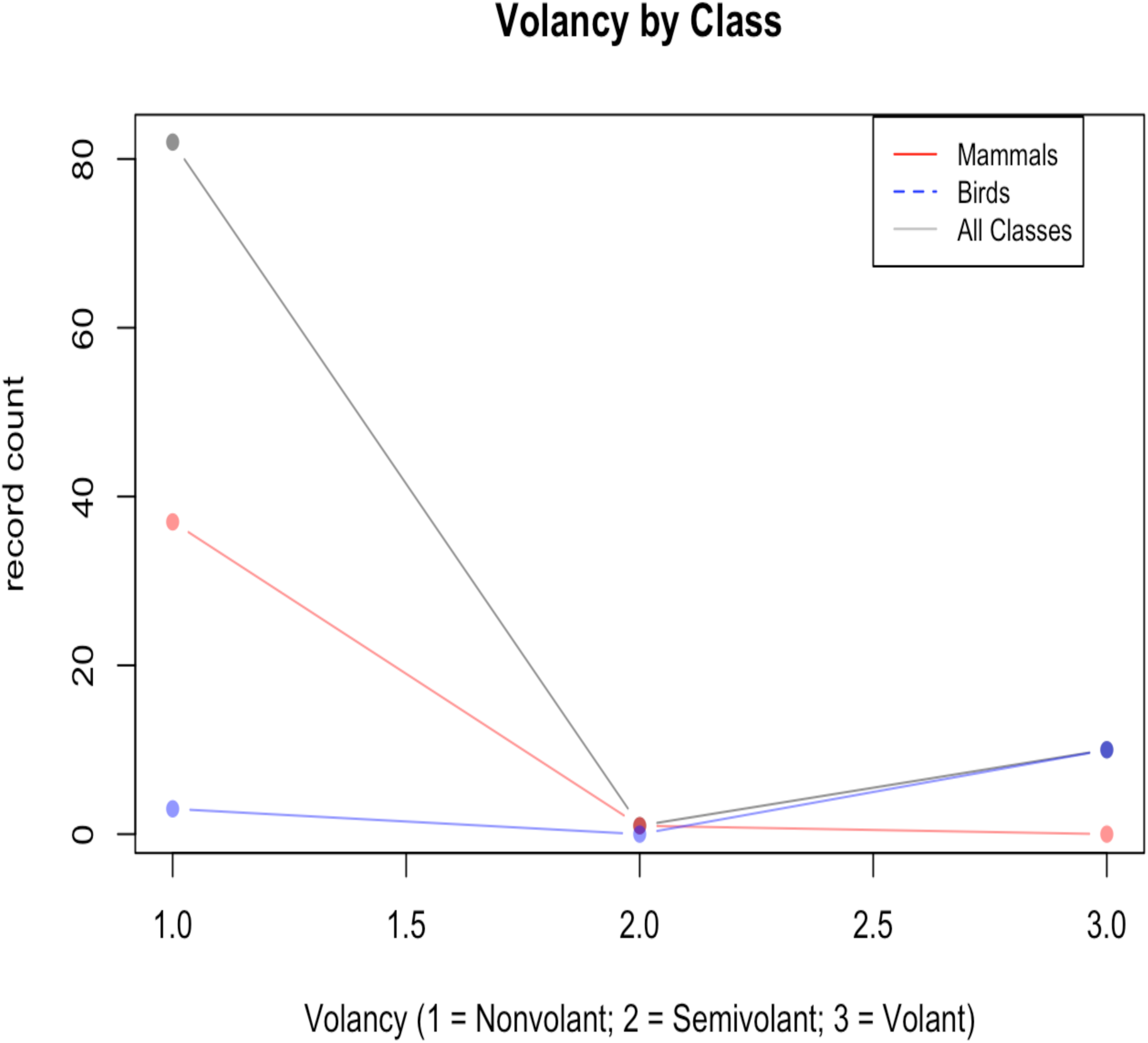

`If one is more generally interested in the class breakdown of factors across a trait field without a priori interest in a specific variable, one can take advantage of the tools.`

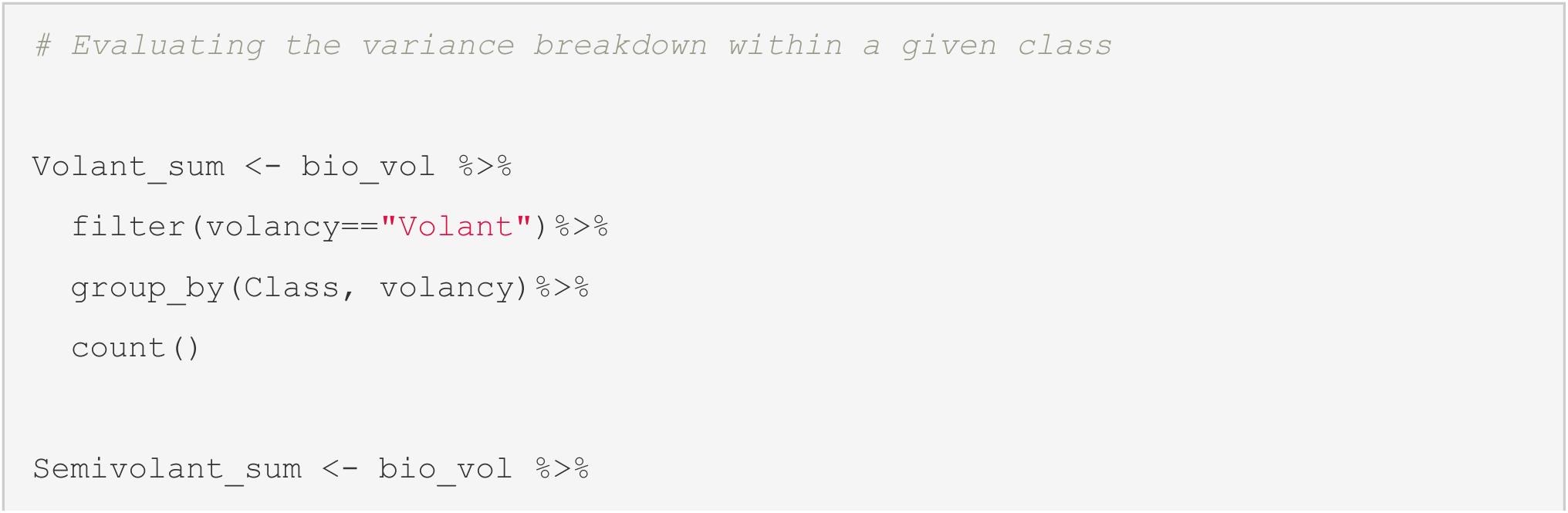
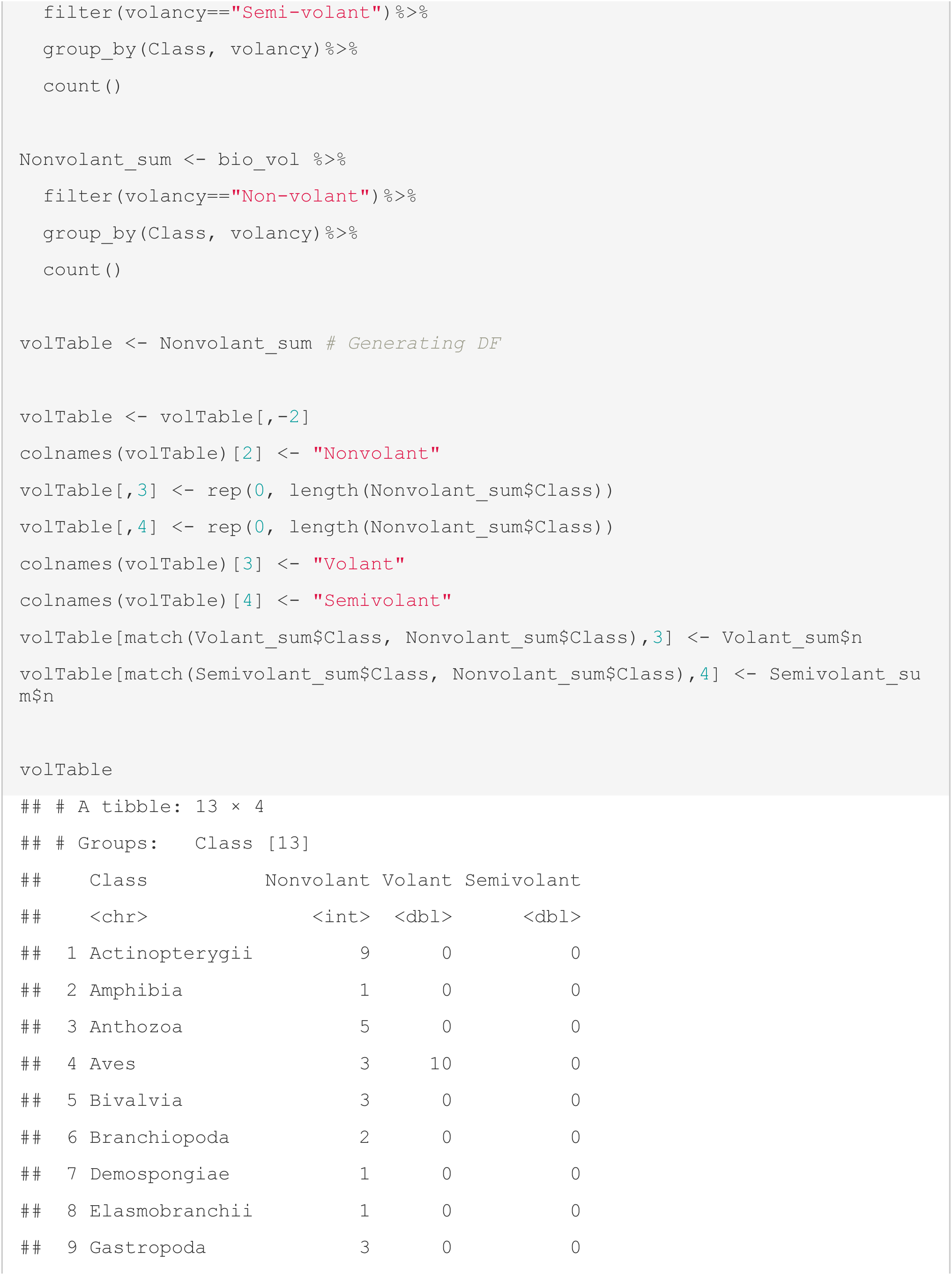
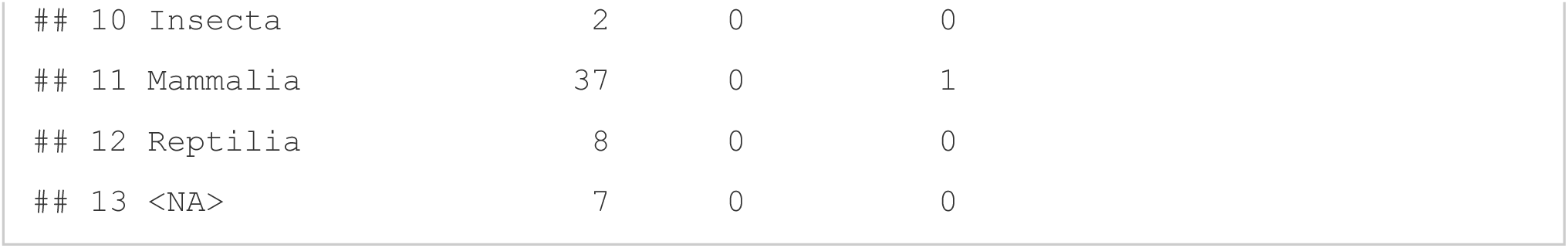

### Basic Regression using MOSAIC

http://mosaicdatabase.web.ox.ac.uk

`Updated 22 February 2022`

#### Vignette #3 - Basic Regression Using MOSAIC & COMPADRE

MOSAIC was built with the intention of its use in comparative biodemography - to strengthen the map of which functional traits predict vital rates and which do not. Traits in the MOSAIC database (factorial and numeric variables) are formatted to be easily accessed and used for regression-based analyses, among other types of analysis. MOSAIC was also designed for quick integration with the COMADRE, COMPADRE, and PADRINO structural population model databases, as outlined in the model exercise below.

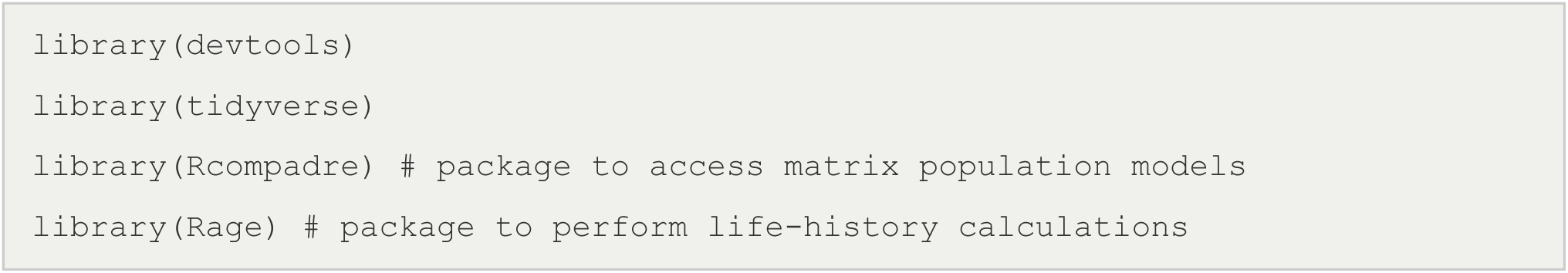

`Accessing MOSAIC`

Download MOSAIC from the mosaic portal. For more information on the basics of downloading MOSAIC and navigating the data structure, see: <u>Vignette #1</u>:

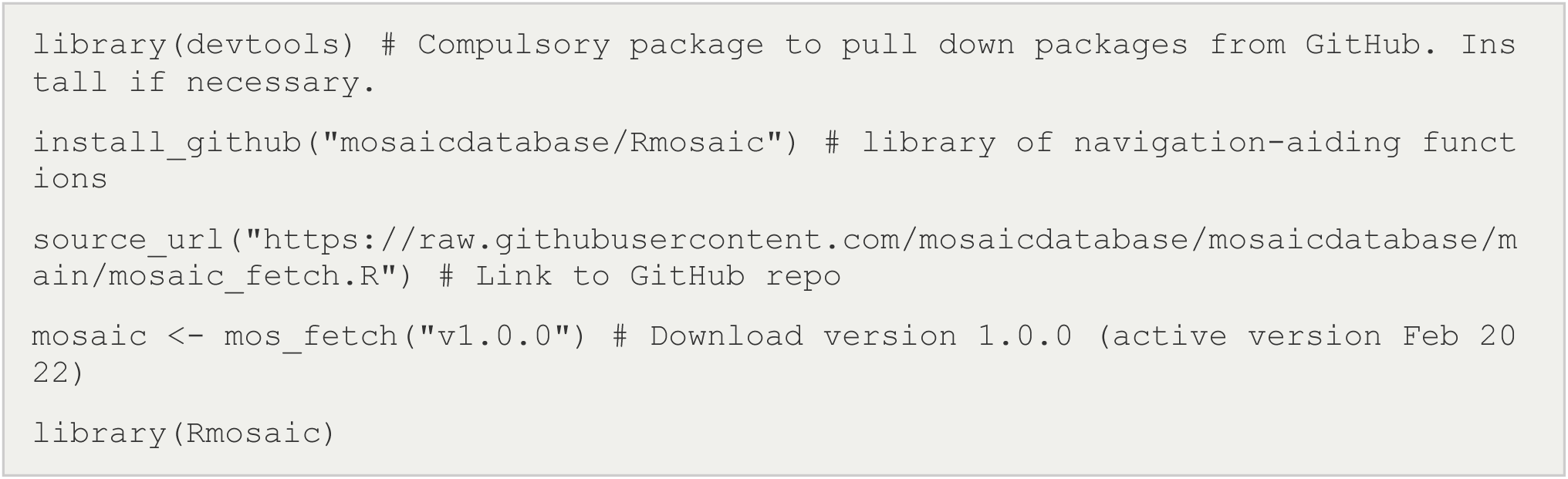

`Regressing Biomass on Generation Time`

In this exercise, the simple bivariate relationship of biomass on generation time is explored through basic regression. Bare in mind, that the below offer the building blocks for more complex multiple regression exercises and other forms of analysis – such as geospatial interpolation and ordination-based practices (see Vignette #4 for a brief vignette of PVA using mosaic).

`#1: Extract biomass data for mammals`

`Extract the biomass values for mammals in the MOSAIC dataset.`

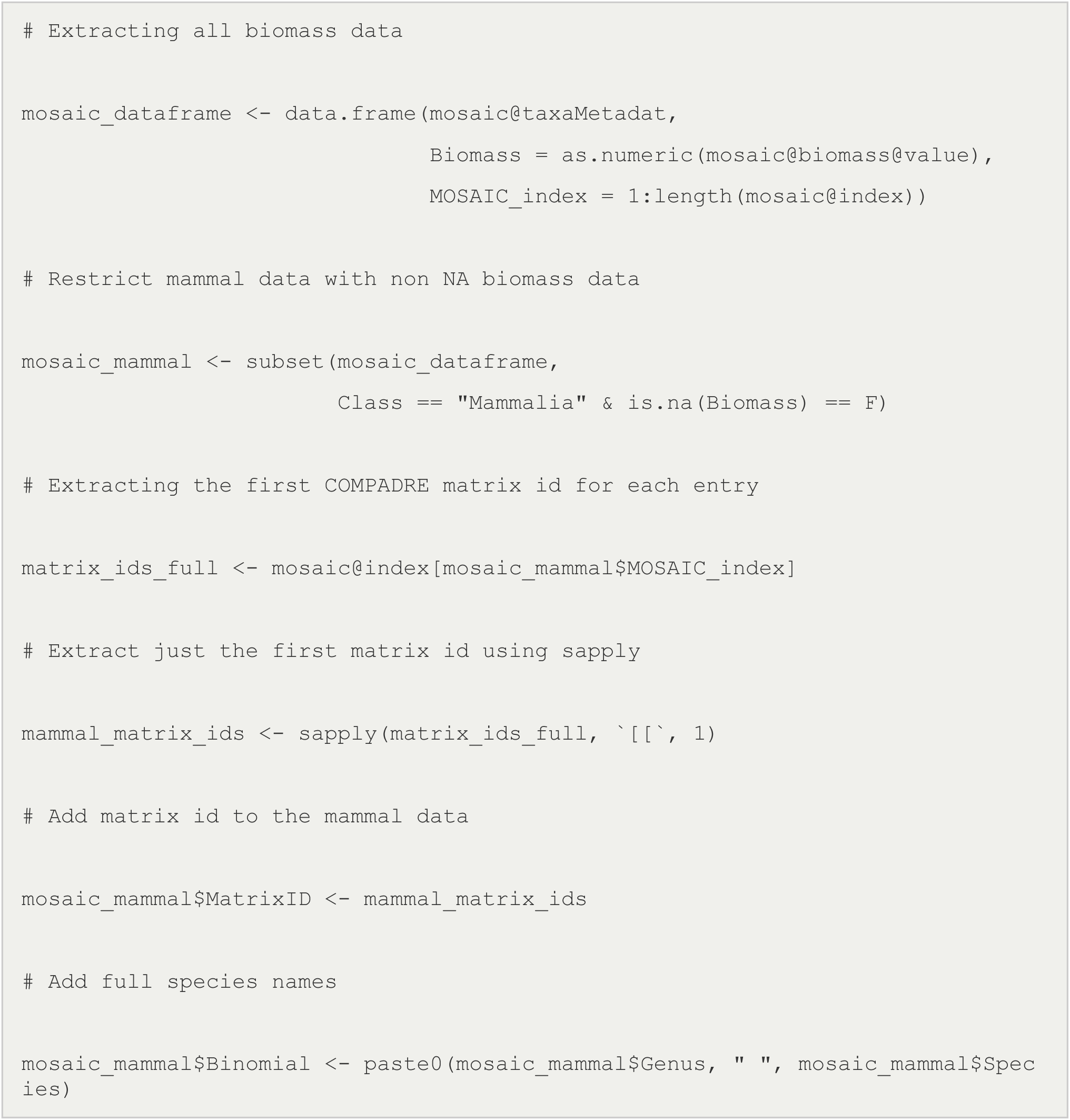

`#2: Download COMADRE matrix population models`

A cdb_fetch() function is used to clone the Comadre database <www.compadre-db.org≥ - which is an s3 copy of a SQL-based relational database - into an S4 data object locally manipulable in R.

Using ids associated with each matrix population model, we will subset the Comadre database of matrix population models for the animals. First we use the Rcompadre package to ‘fetch’ the Comadre database.

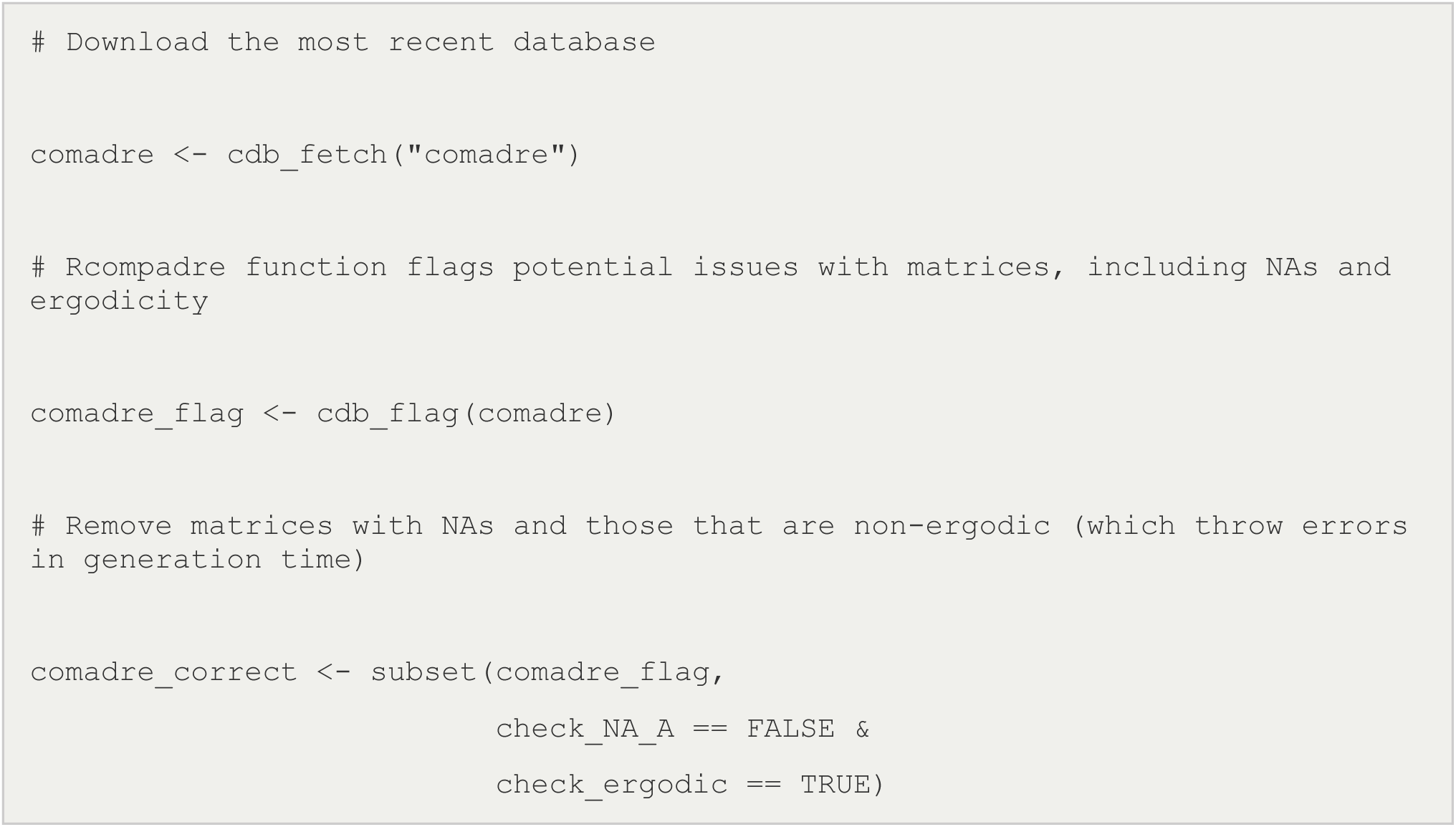

`The subsets of mosaic and Compadre can be overlapped in one line of code.`

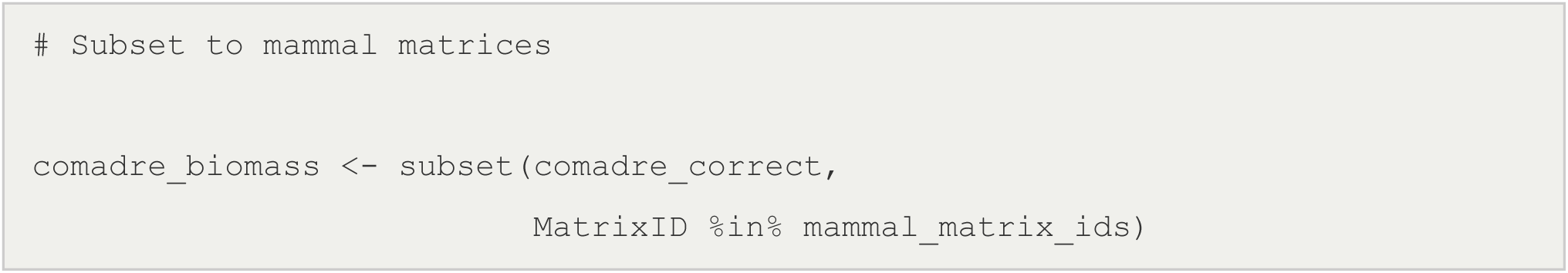

`3. Calculating generation time`

Rcompadre enables sub-setting of a matrix population model (life table) into constituent matrices representing the survival and fecundity/reproductive components (U and F matrices, respectively). Functions in the Rage package often require the specification of both the U and the F matrix, not the combined A matrix. The Rage package contains a suite of common demographic calculations - including GENERATION TIME - which are used to simplify this exercise. Note that other common demographic derivations, including transient indices can be extracted using the identical procedure.

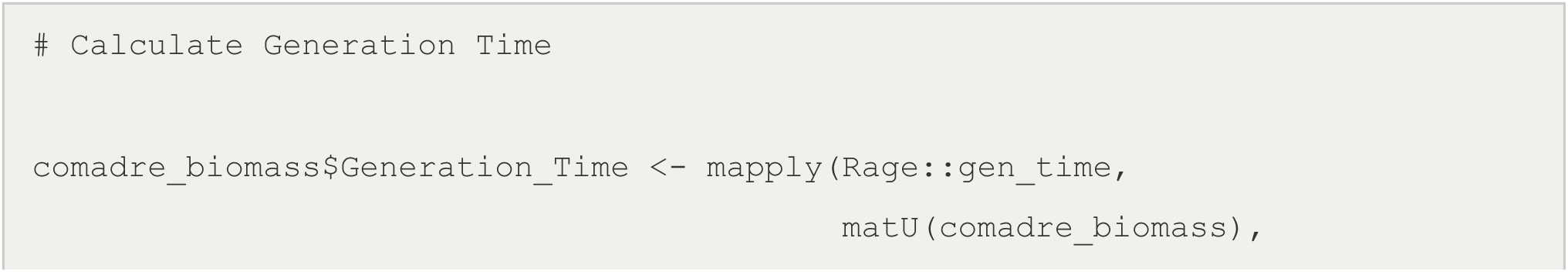
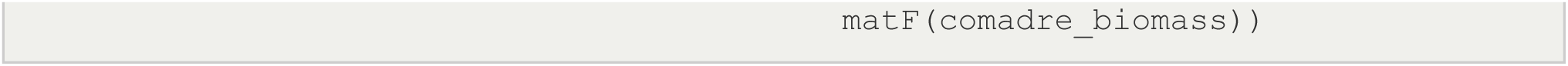

`4. Explore the relationship of generation time data and biomass data`

`Matrix IDs can be used to bridge generation time from comaadre with mosaic records.`

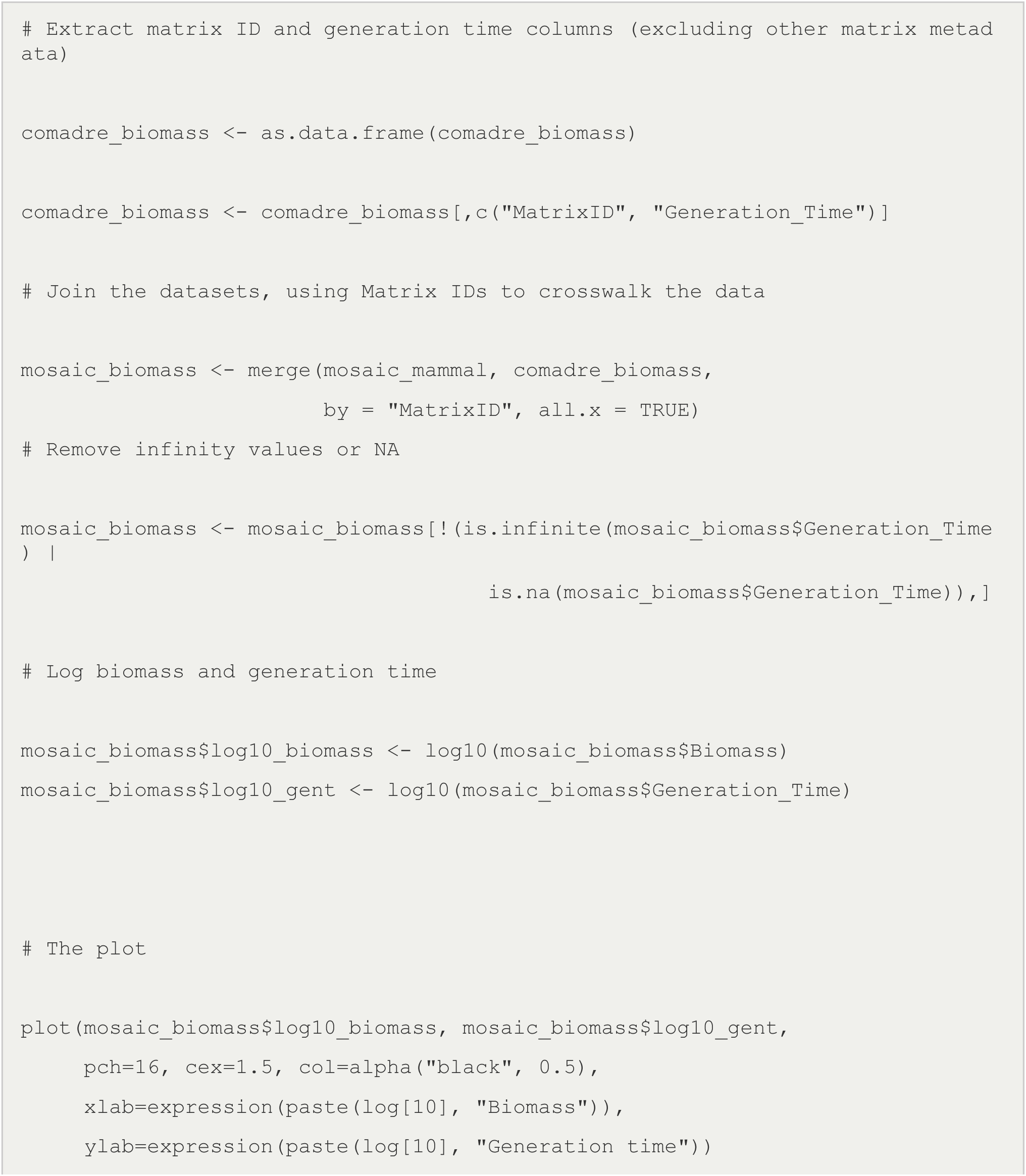
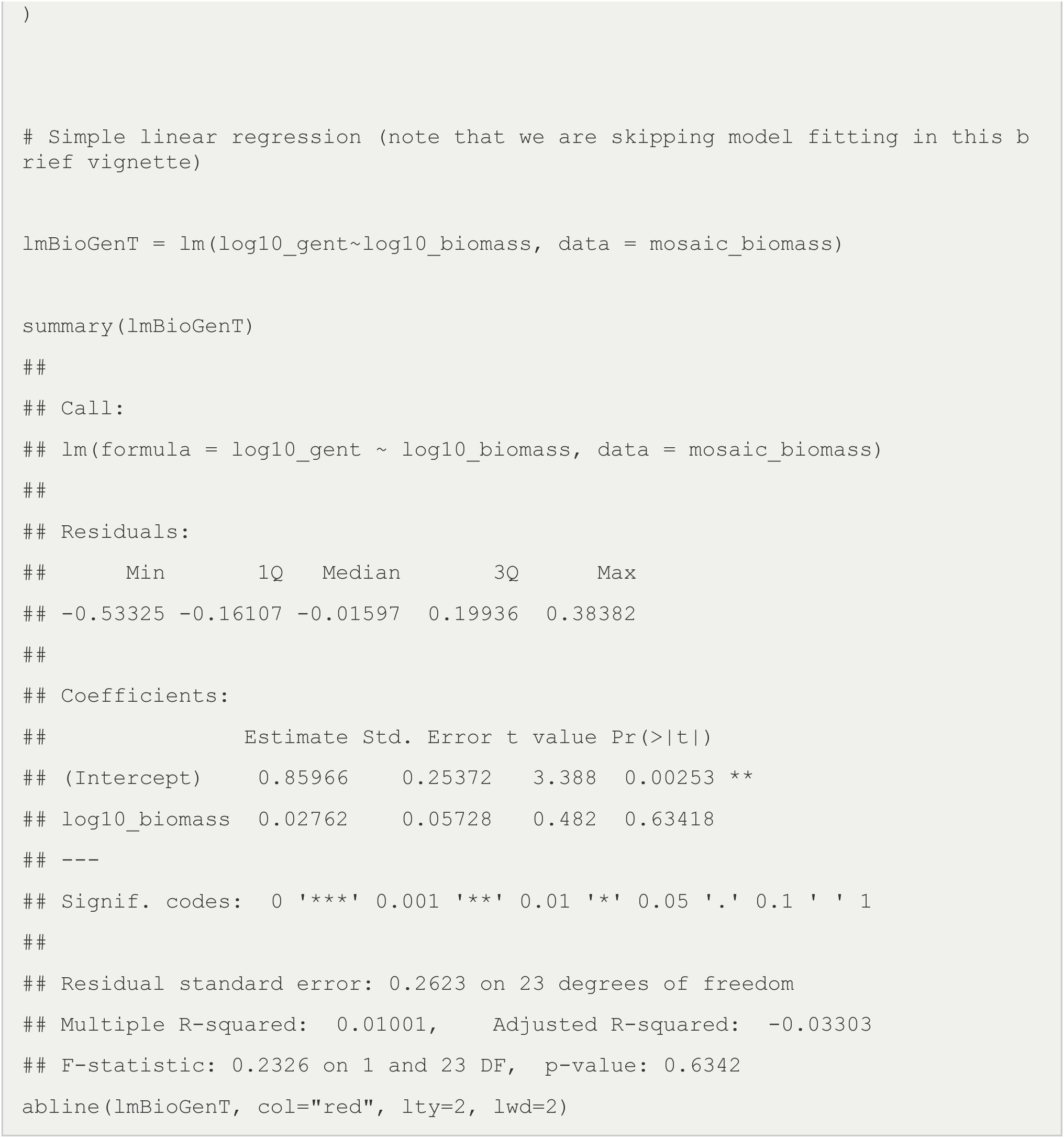

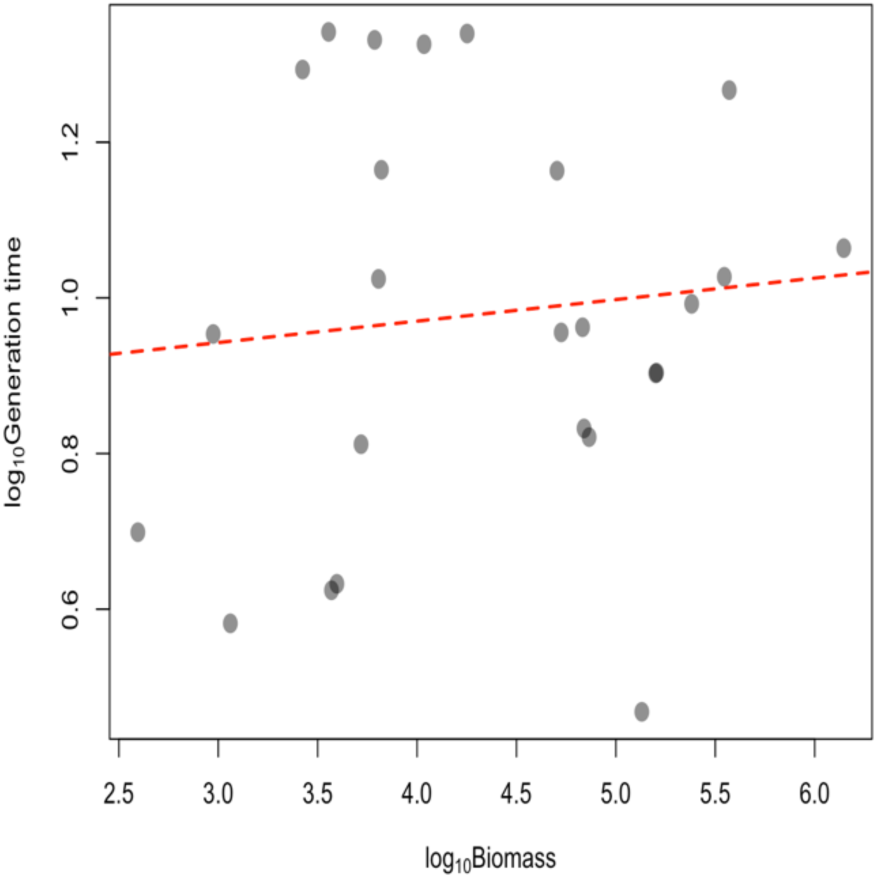

### Dimension Reduction + MOSAIC

http://mosaicdatabase.web.ox.ac.uk

`Updated 14 March 2022`

#### Vignette #4 – Dimension Reduction + MOSAIC

MOSAIC was built for integrated analyses across trait fields. Whether the data are continuous (e.g., biomass) or categorical (e.g., dispersal class), MOSAIC allows researchers to explore these dimensions to identify the primary axes of variation across functional traits. Here, we will use a Multiple Factor Analysis (MFA) to explore the primary axes of variation between: biomass, dispersal capacity, growth determination, habitat type and mating system.

Note: These traits are picked solely to illustrate the analysis method that can be scaled for other traits in MOSAIC, demographic rates in COMADRE, COMPADRE and PADRINO and climate data from ERA5-Land.

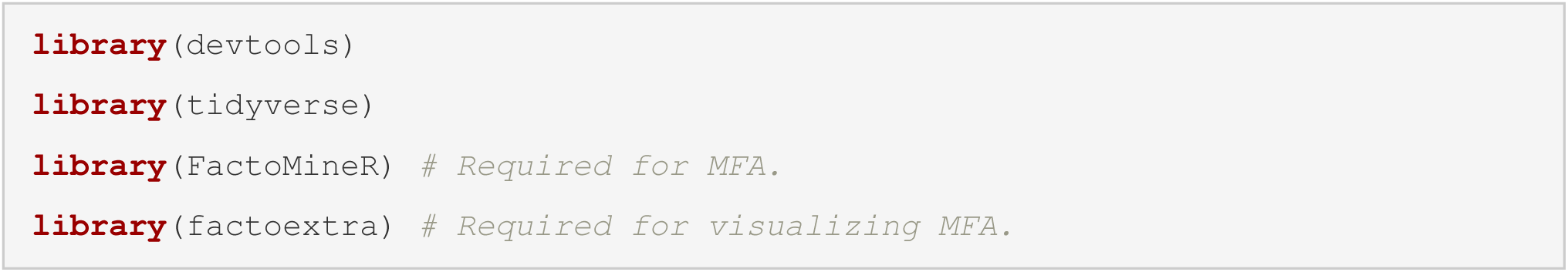

`Accessing MOSAIC`

Download MOSAIC from the mosaic portal. For more information on the basics of downloading MOSAIC and navigating the data structure, see: <u>Vignette #1</u>:

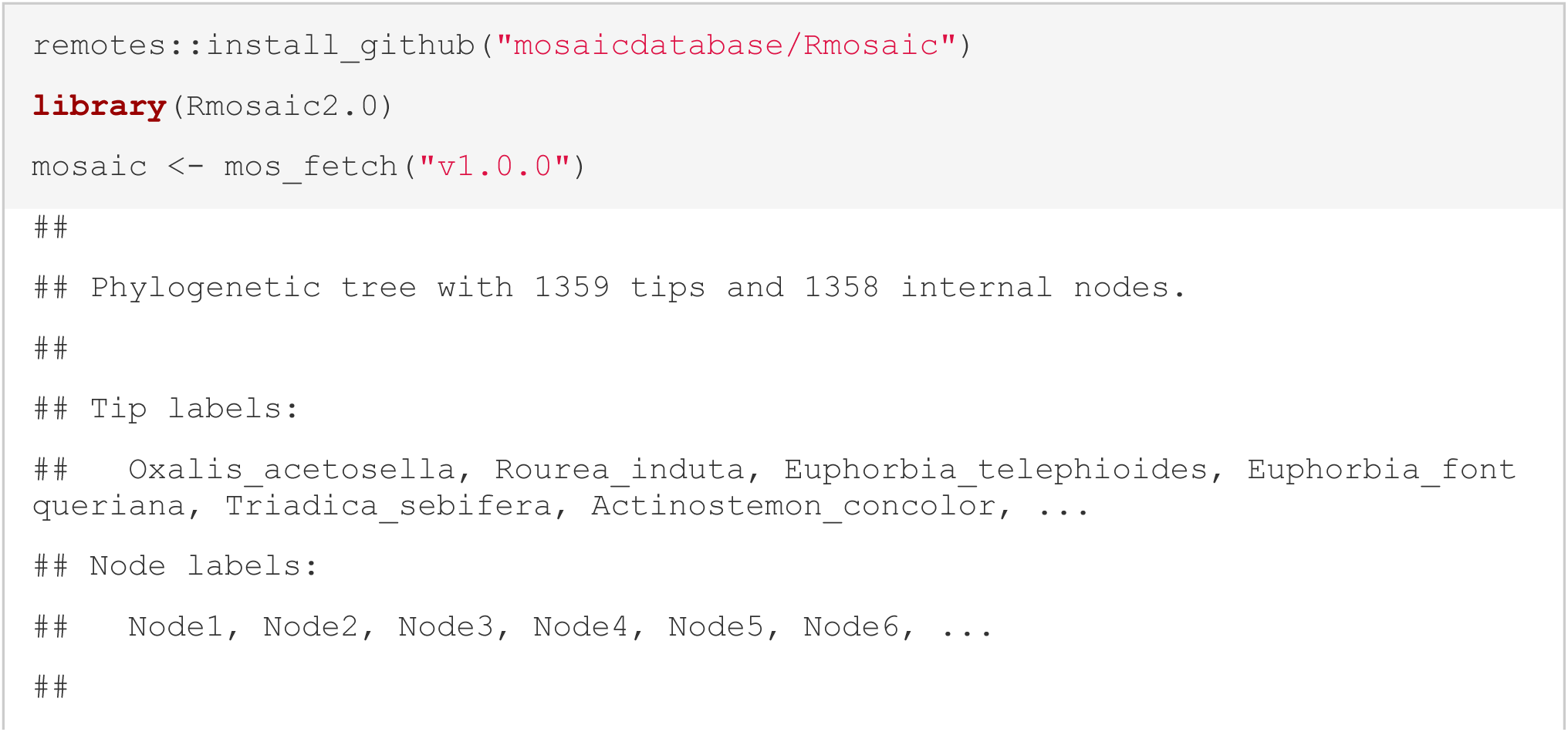
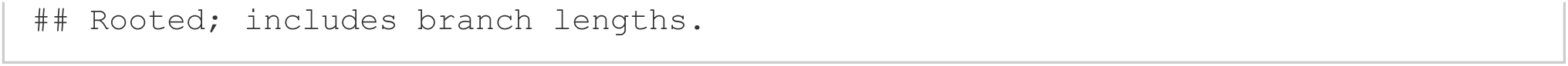

`Generating Dataframe`

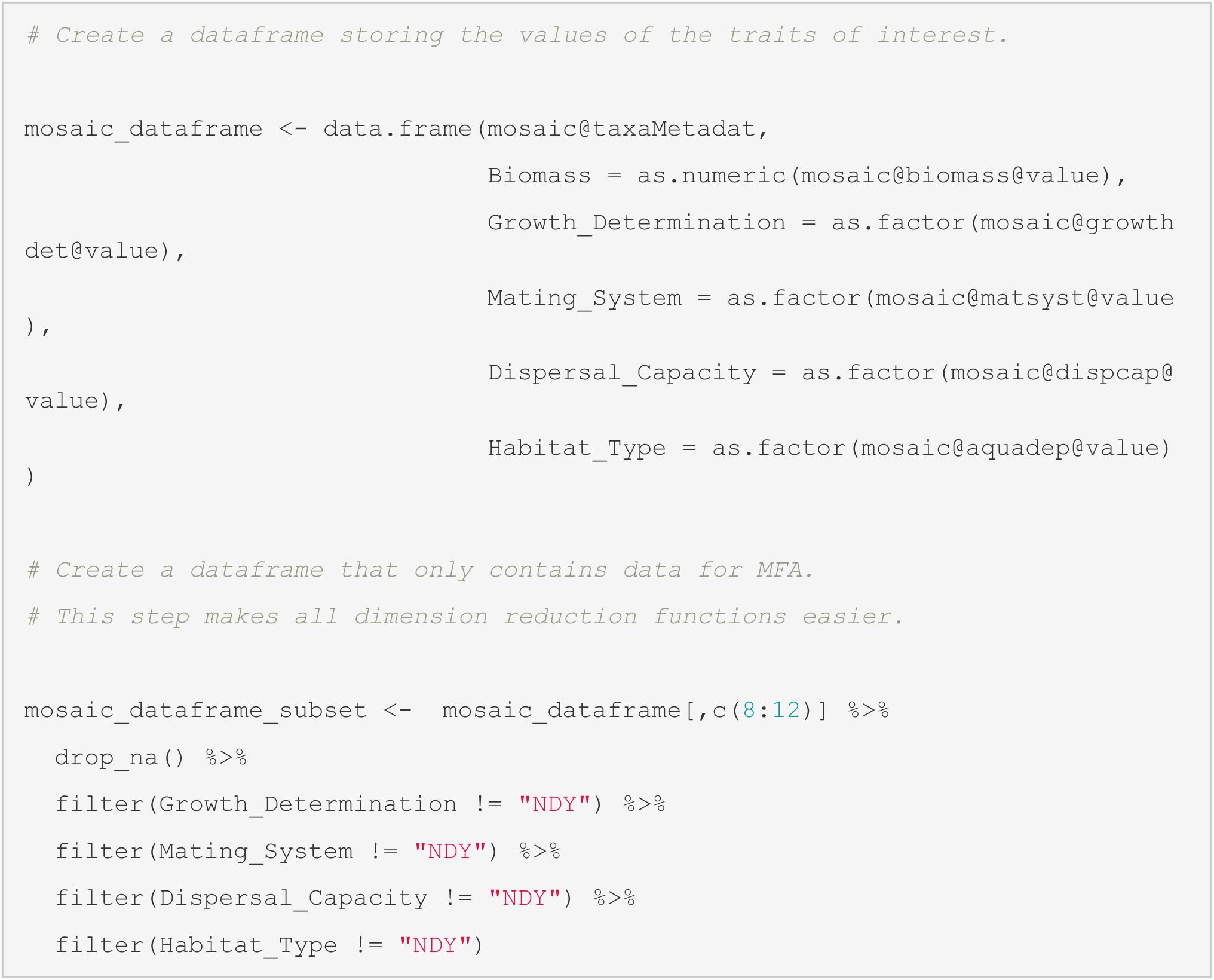

`Perform MFA`

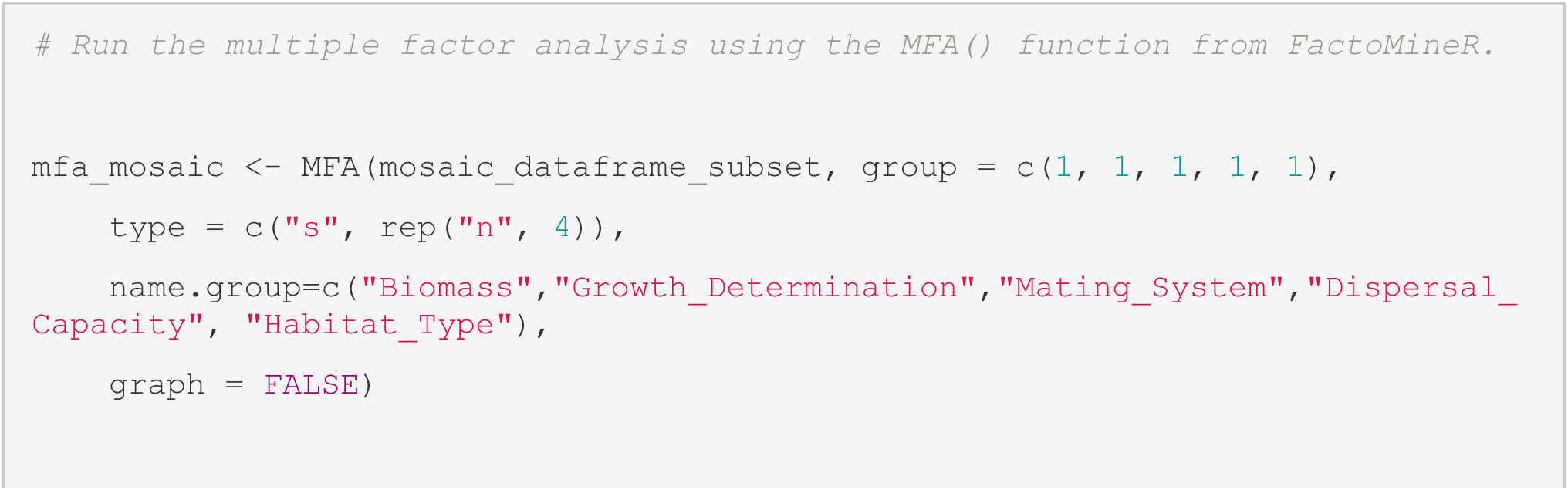
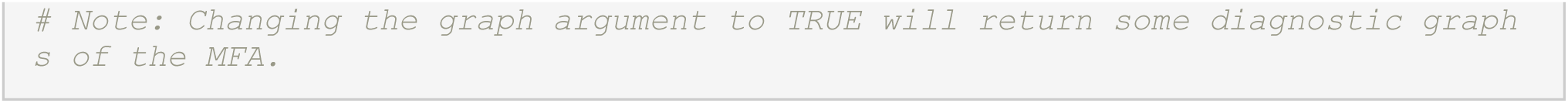

`Explore MFA`

In this hypothetical example, we are interested in what traits covary with the primary axes of variation in the MFA (i.e., Dim1 and Dim2). To visualize these relationships, we can use the in-built fviz functions in the factoextra package to illustrate (1) position of individuals, (2) average effect of trait level (i.e. monogamous) on individual position within the 2 dimensions and (3) average effect of trait type (i.e. mating system).

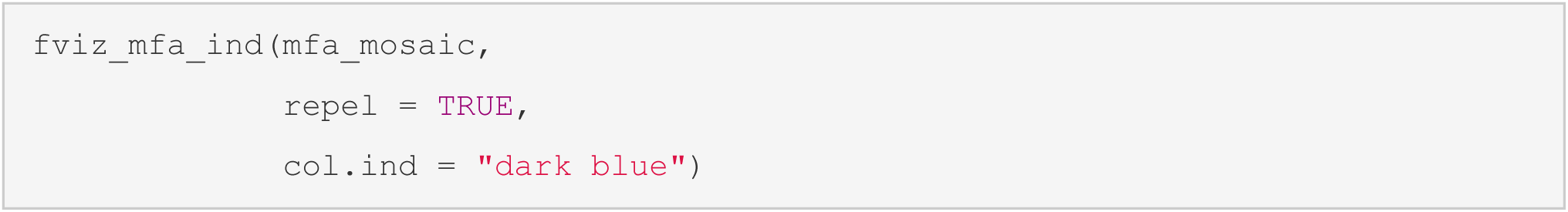

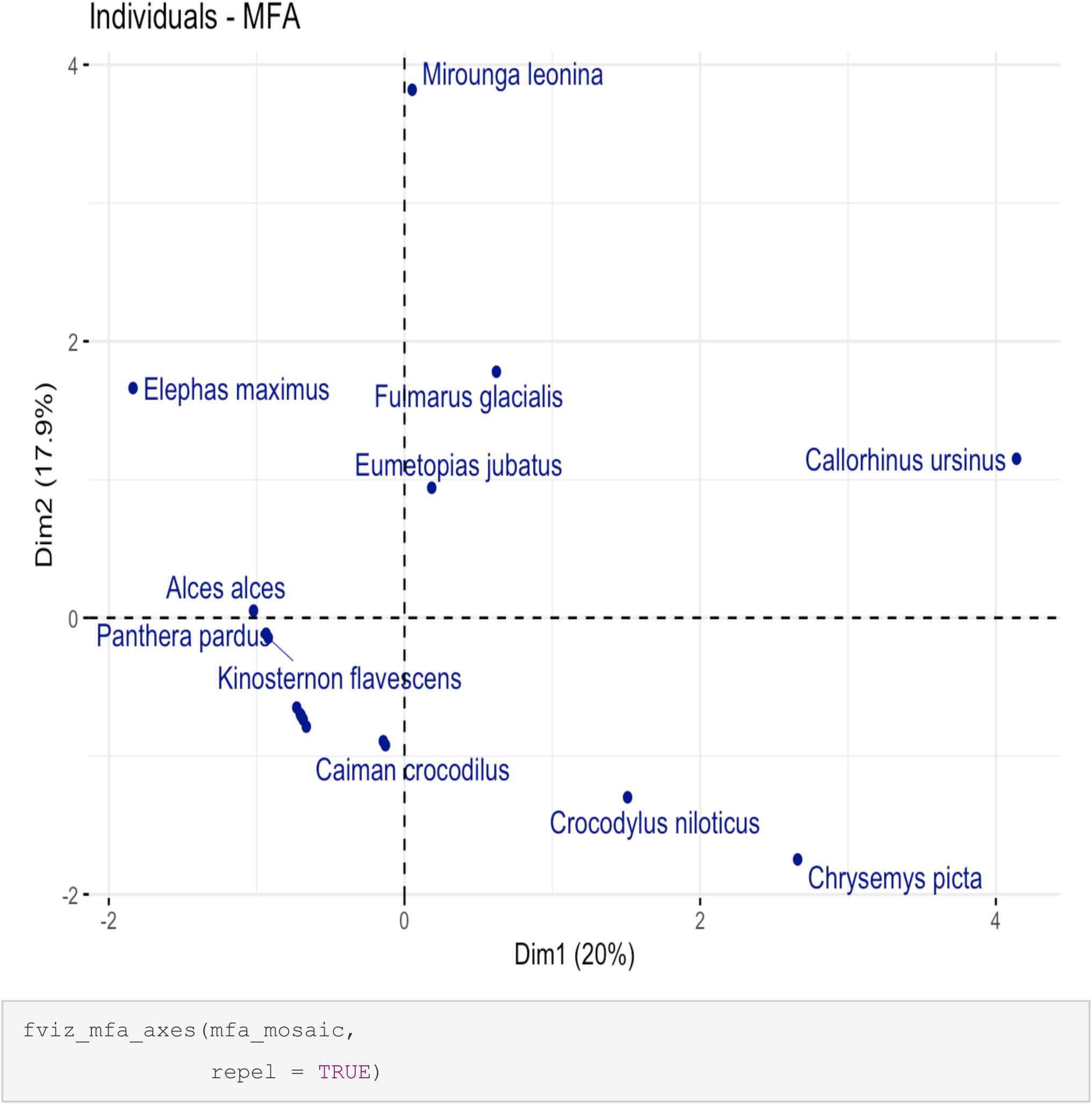

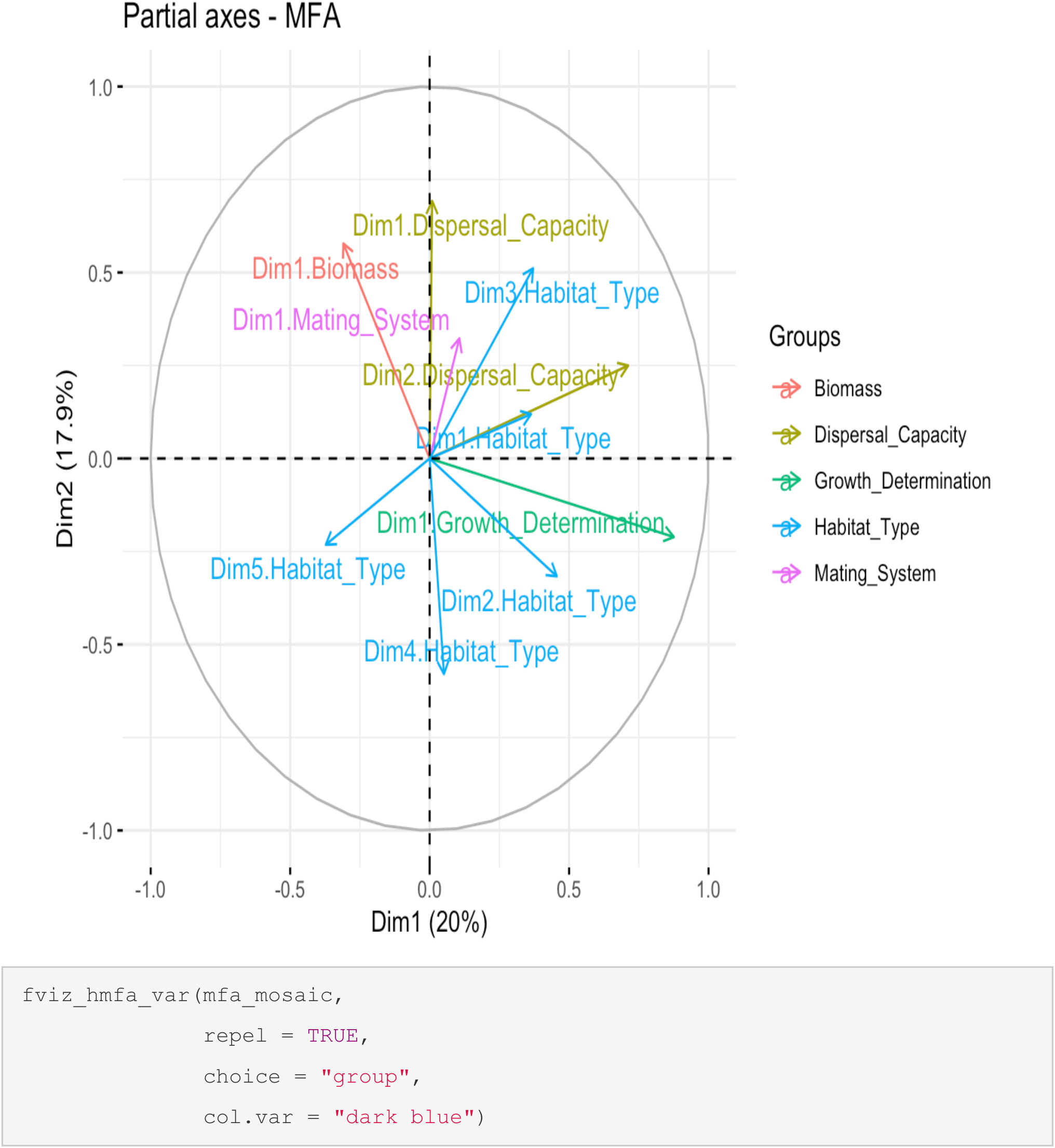

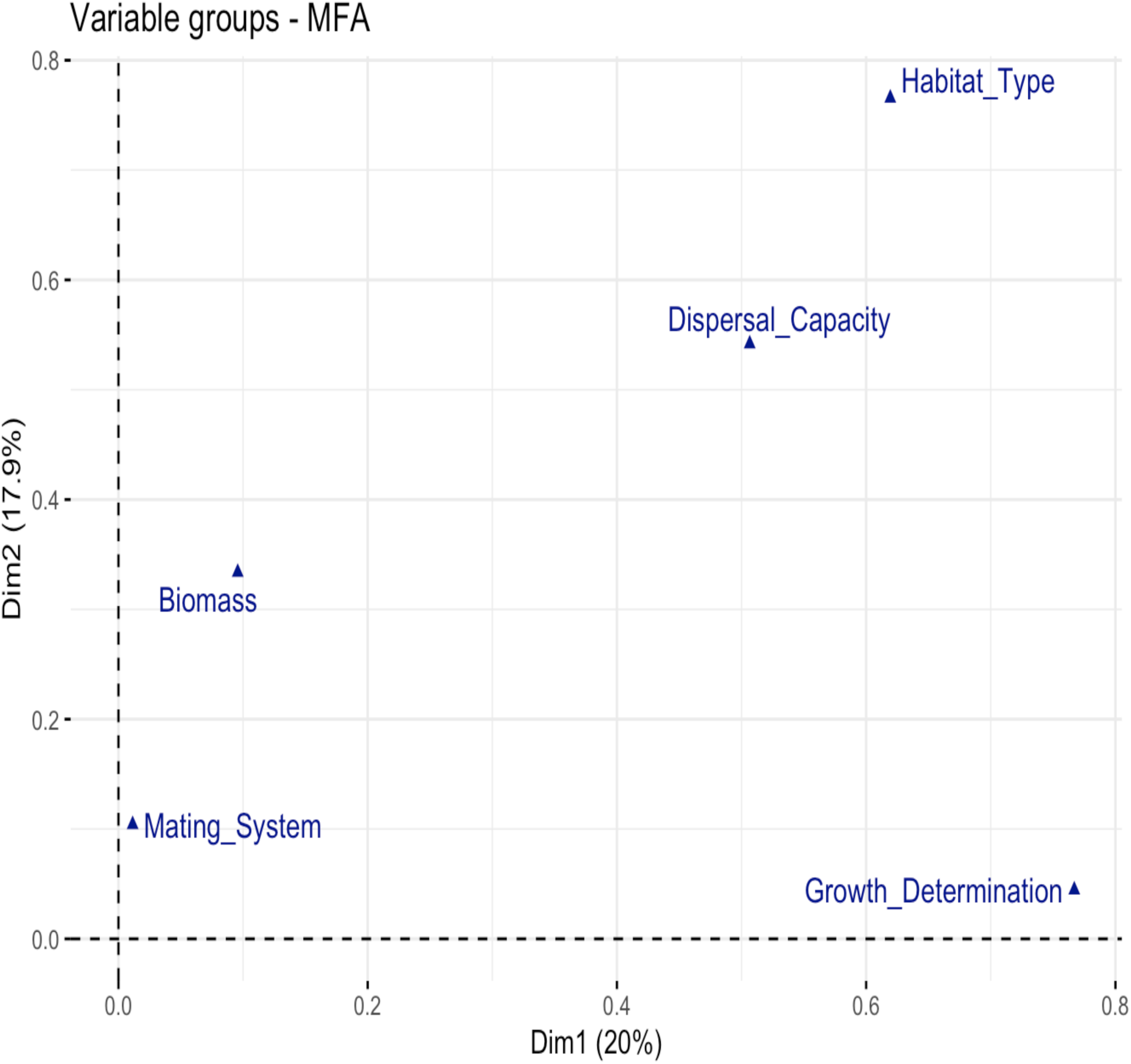

### Climate Data + MOSAIC

http://mosaicdatabase.web.ox.ac.uk

`Updated 14 March 2022`

#### Vignette #5 – Climate Data + MOSAIC

MOSAIC was built with the intention of its use in comparative biodemography - to strengthen the map of which functional traits predict vital rates and which do not. Traits in the MOSAIC database (factorial and numeric variables) are formatted to be easily accessed and used for regression-based analyses, among other types of analysis. MOSAIC was also designed with the analysis of demographic metrics and traits in their environmental contexts. To this end, MOSAIC includes a separate object (climate) which we will access and make use of in the following exercise.

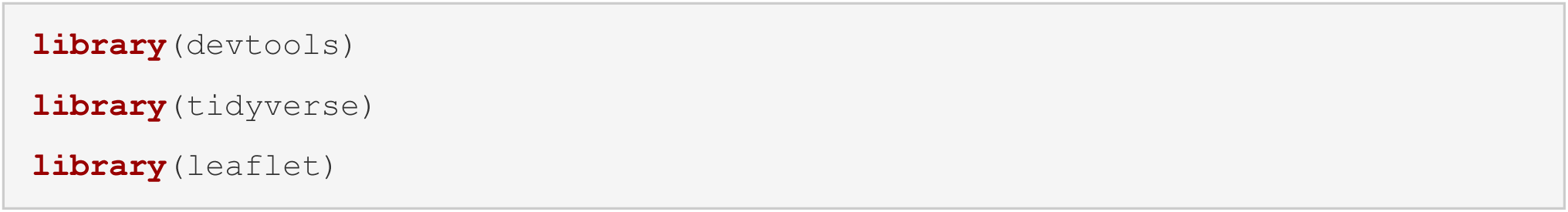

`Accessing MOSAIC`

`Download MOSAIC from the mosaic portal. For more information on the basics of downloading MOSAIC and navigating the data structure, see: Vignette #1:`

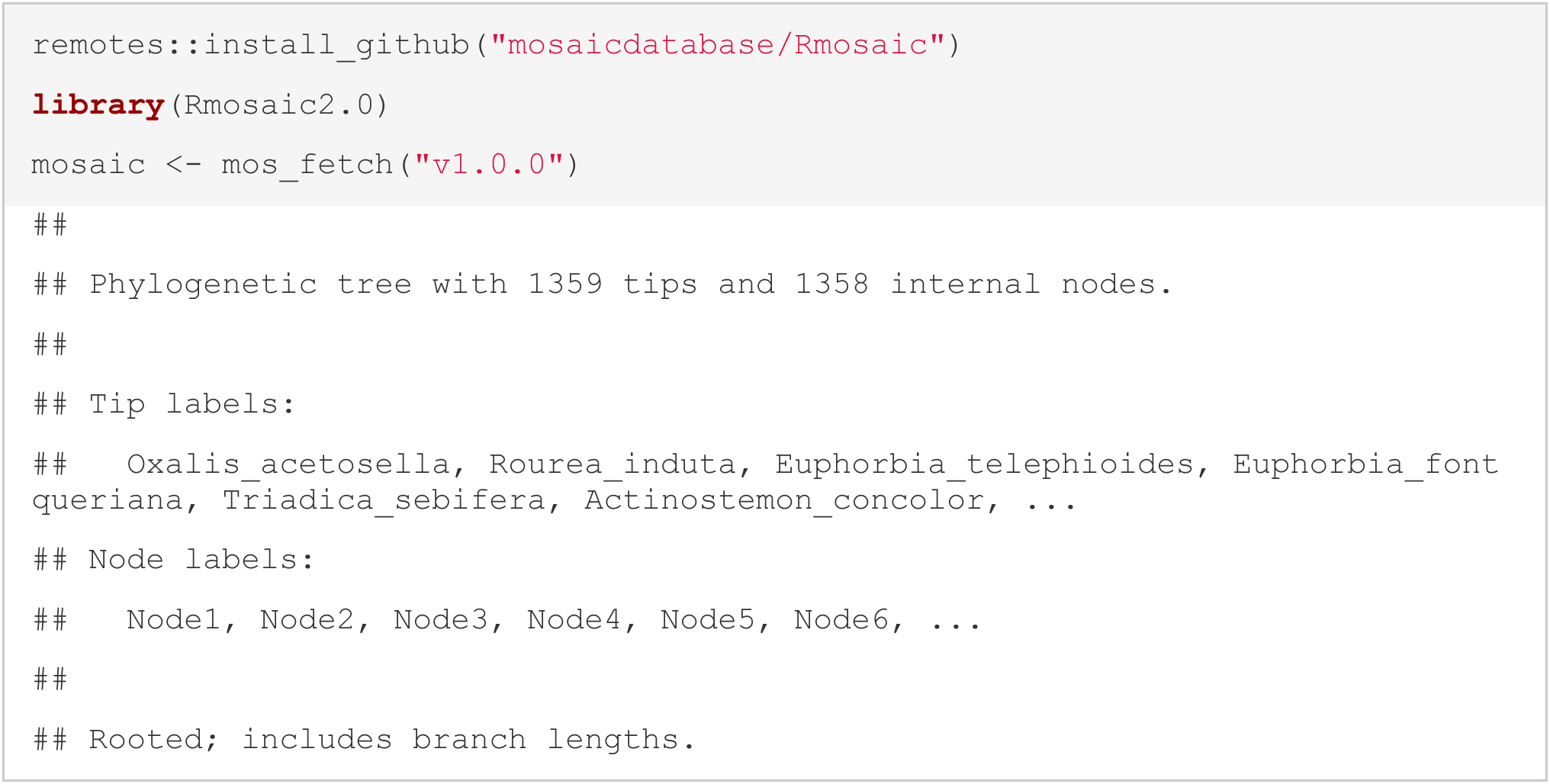

`Selecting Target Matrices`

`To get this exercise started, we first need to select which part of MOSAIC we are interested in. For simplicity, we are here focusing on mammal biomass as was done in Vignette #2:`

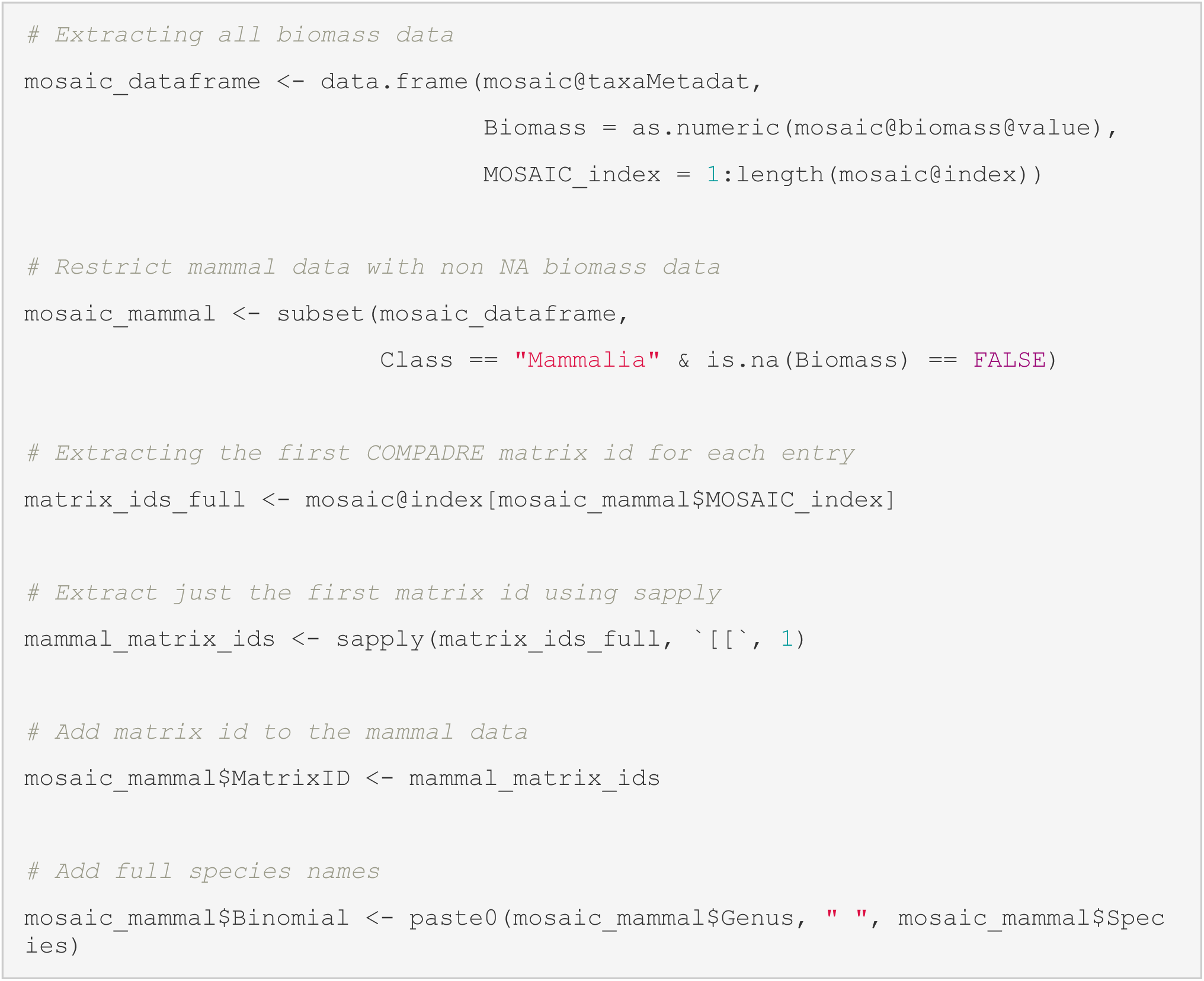

`We now have a data frame containing MOSAIC IDs, taxonomic information, and biomass values for all mammals within MOSAIC. The corresponding matrix IDs are:`

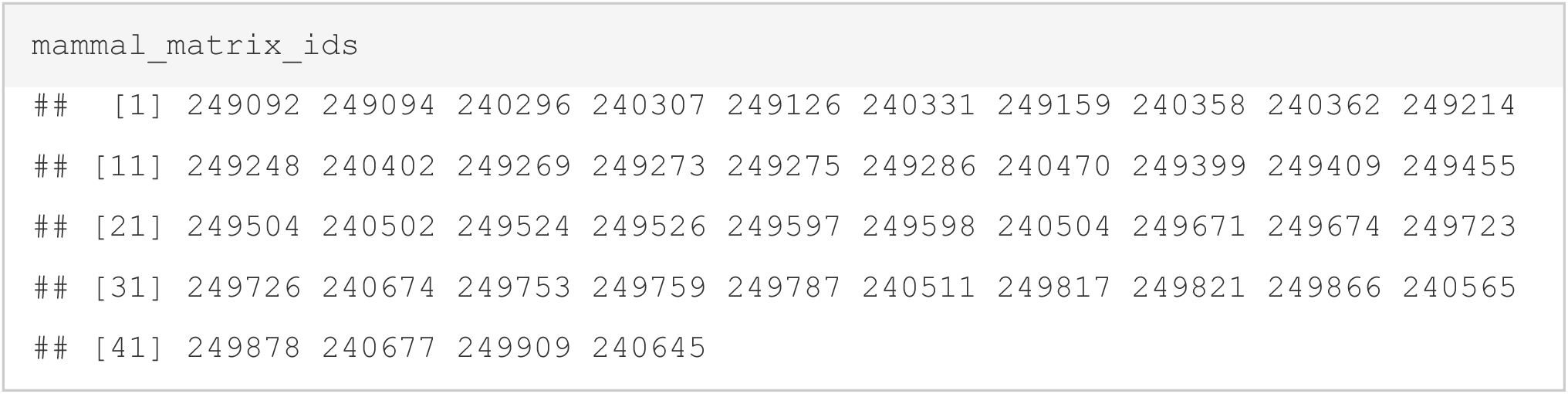

`Accessing the Climate Data`

With our matrix IDs at hand for analysis, we are now ready to tap into the climate data pool contained within MOSAIC. The climate data hereafter, has been obtained using the R-package KrigR and climate data is reported as mean and standard deviation of monthly time-series belonging to the location and study duration for each matrix contained within MOSAIC.

`Climate Data in MOSAIC`

`The climate data within MOSAIC is stored in the climate object:`

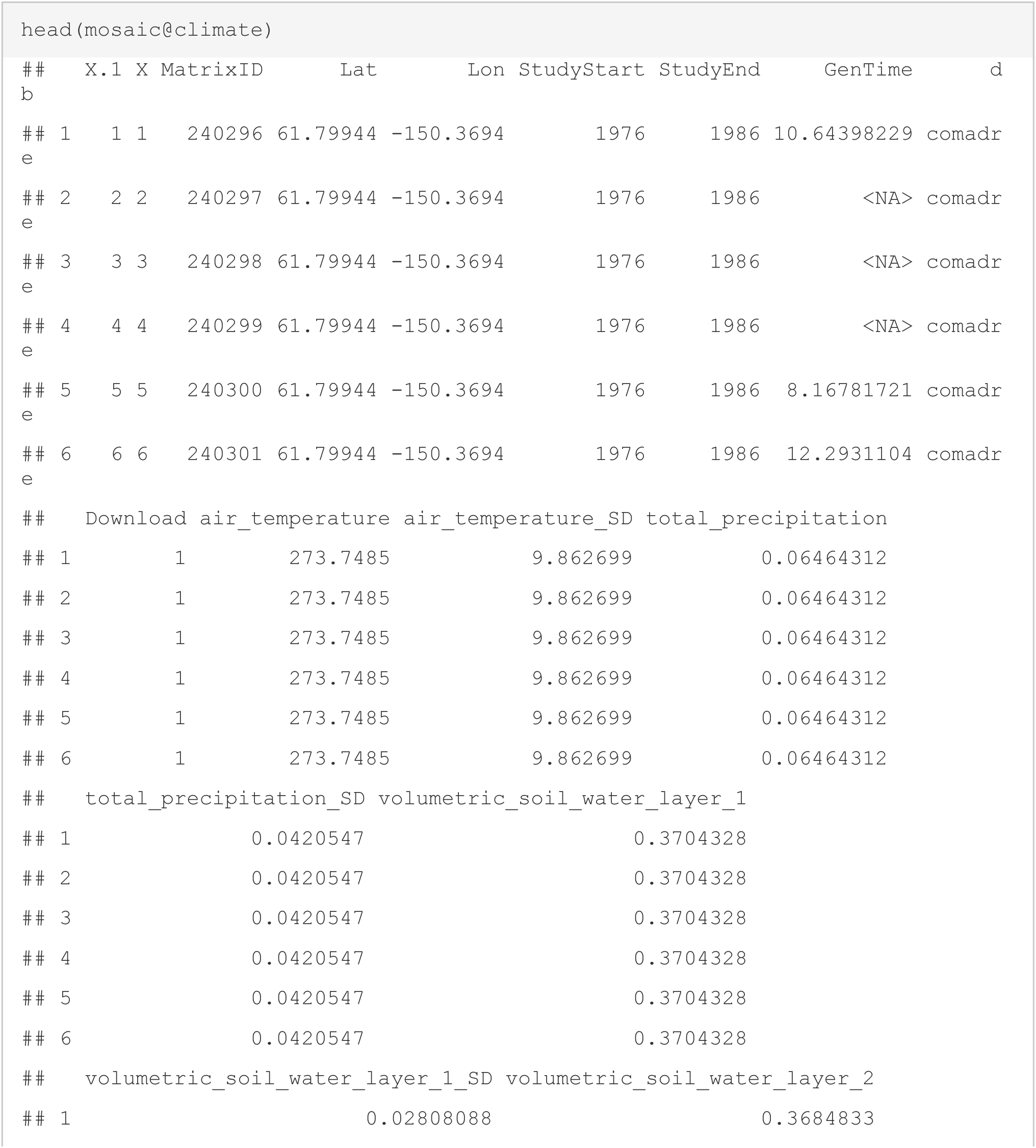
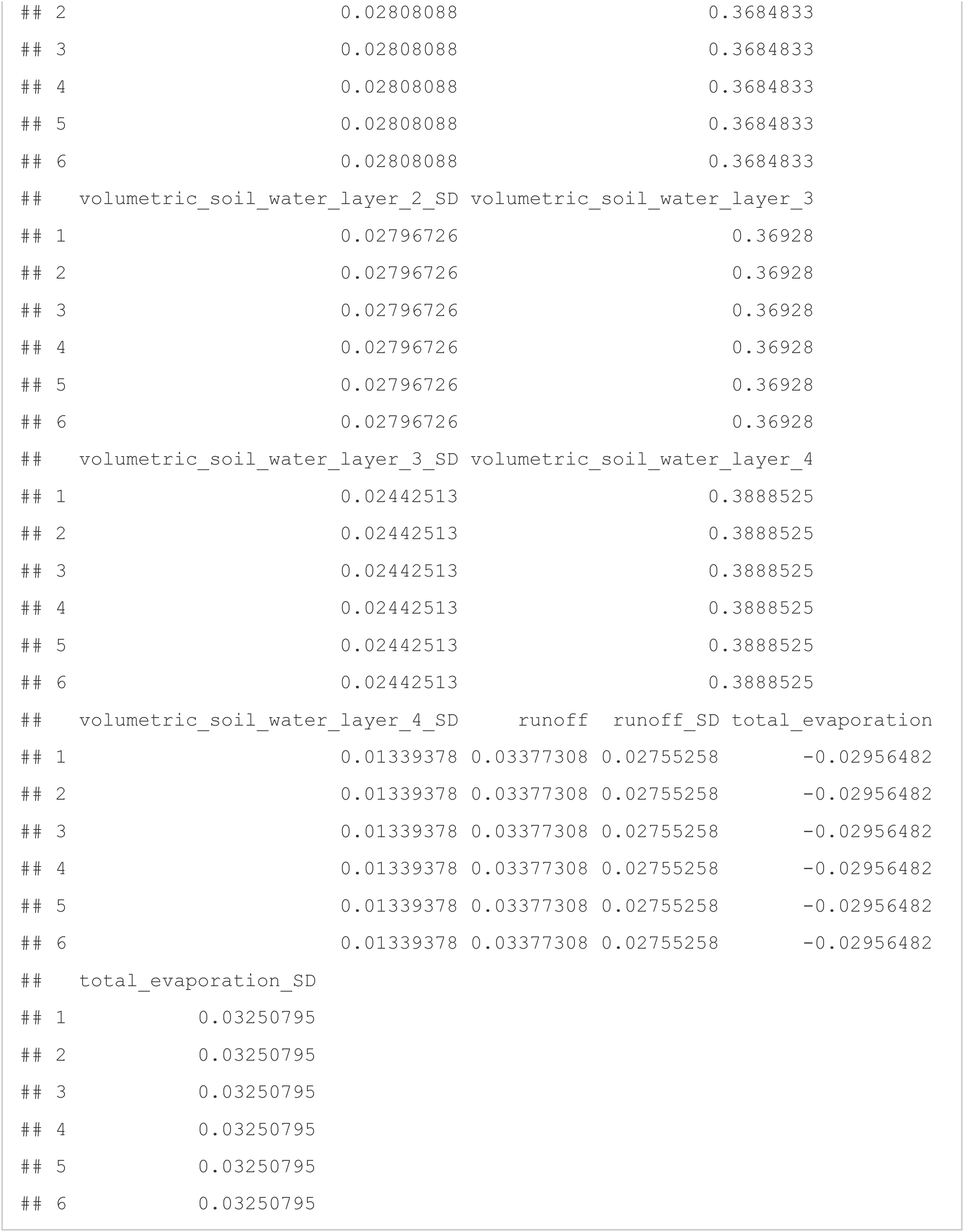

As you can see, there are NA values for some matrix IDs in the above output. This results from either (1) missing geolocations of the underlying matrix models making extraction of environmental parameters impossible, (2) study durations of matrix models pre-dating climate data availability at monthly scale within ERA5-Land, or (3) geolocation of matrix models falling outside of land masses.

`Extracting Climate Data`

Since we are only interested in a subset of matrix models, we do not require the full climate data object and so we can refer to only the relevant subset of the climate object by matching MatrixID values:

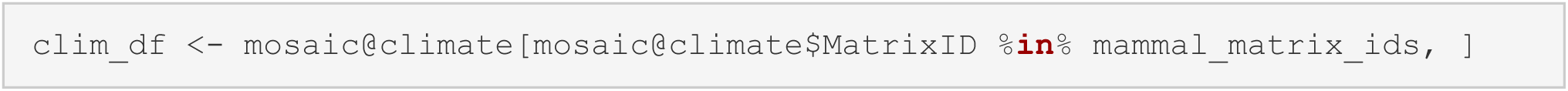

`With the subsetted climate data, we can now create a data frame holding all necessary data for some exploratory analyses:`

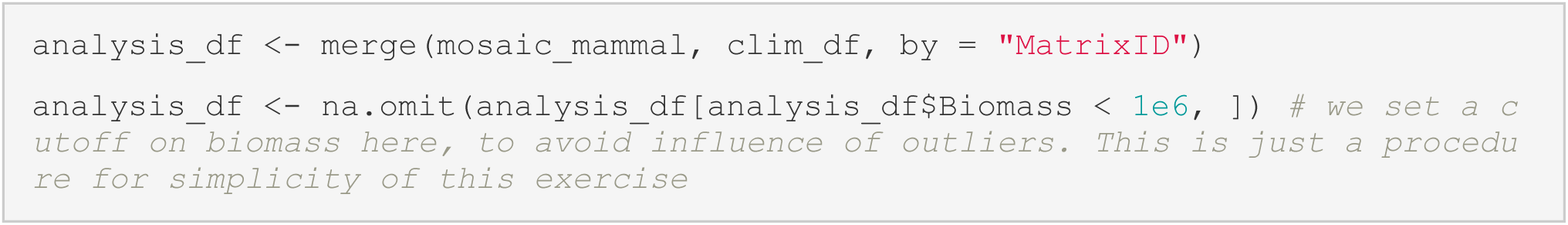

`Let’s plot the remaining data out in their geospatial context:`

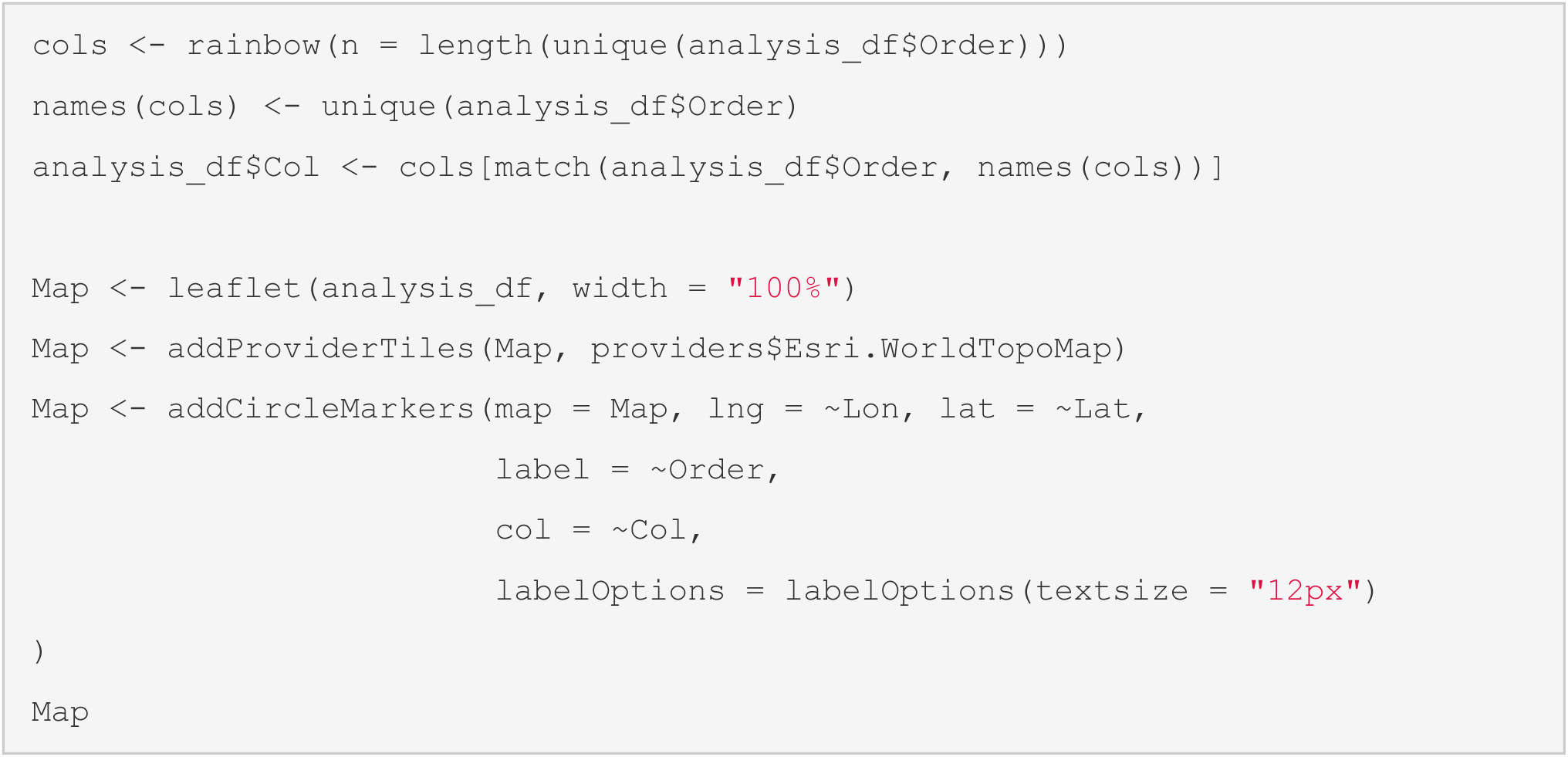

`Regression & Analysis`

Now we are ready to run analyses. The ones we are focussing on here are rudimentary at best and should simply serve as a demonstration of how easily one can make use of state-of-the-art ERA5-Land climate data with the MOSAIC data base.

`All Mammals`

`First, let’s assess the effect of temperature on biomass of all mammals:`

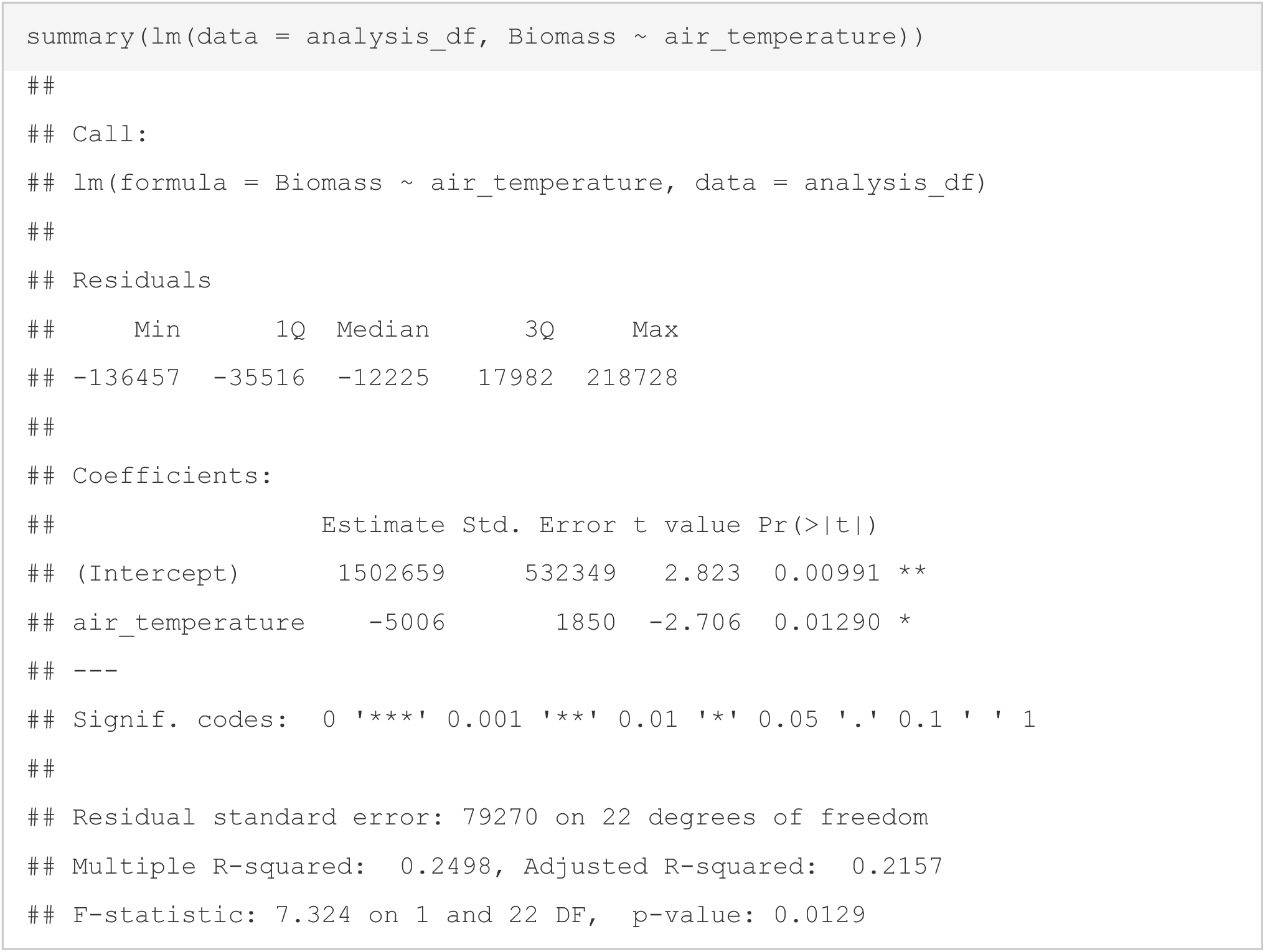

`According to this, mammals in colder regions are heavier. This is in support of Bergmann’s rule.`

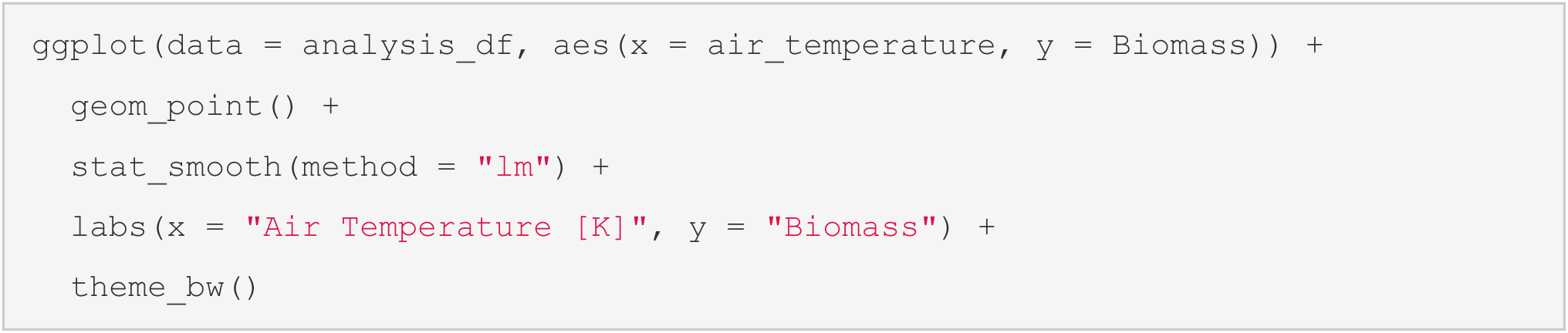

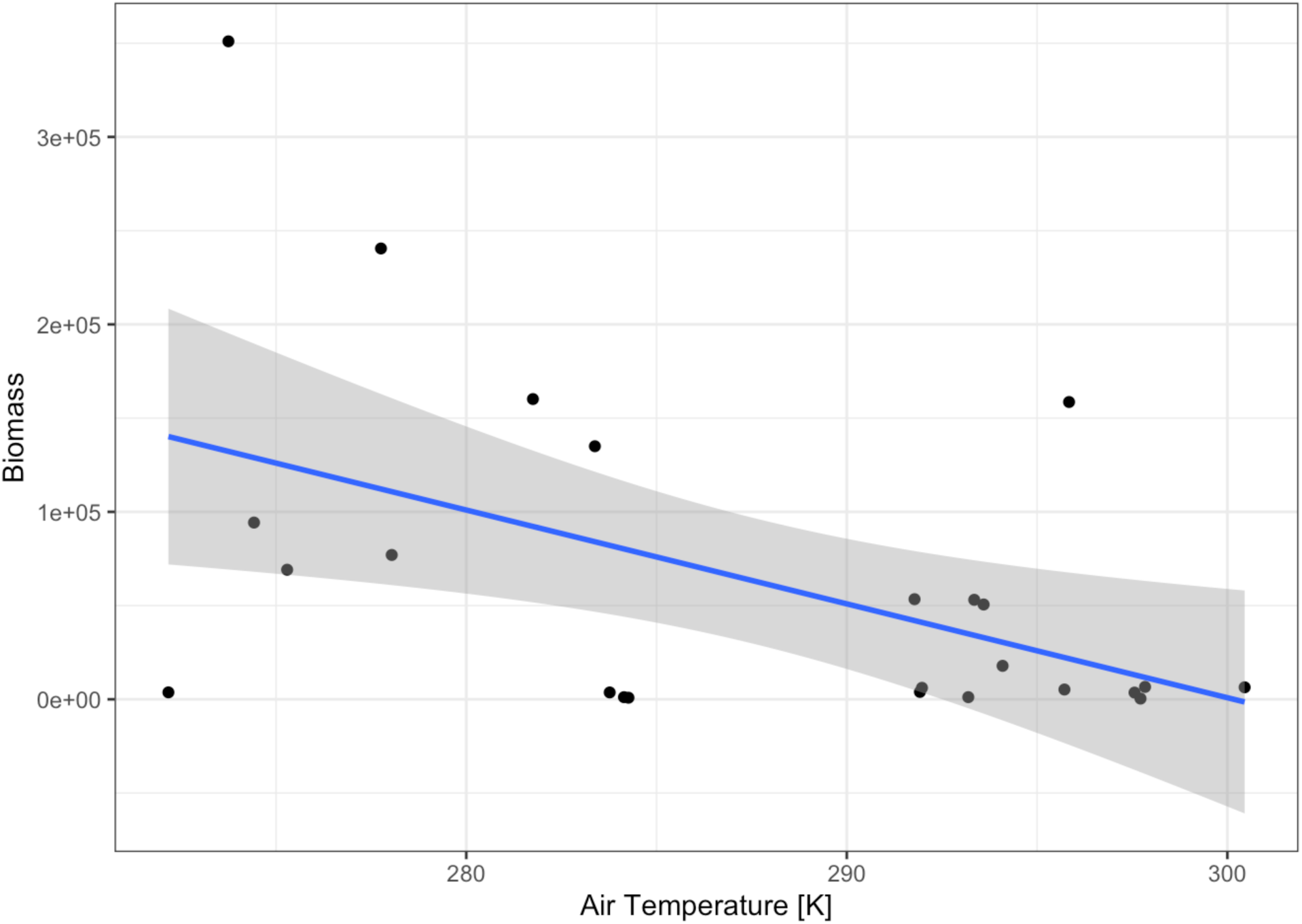

`Mammals by Order`

`Now let’s make use of the higher-resolution taxonomic information available in MOSAIC and assess the impact of air temperature on biomass by order of mammals:`

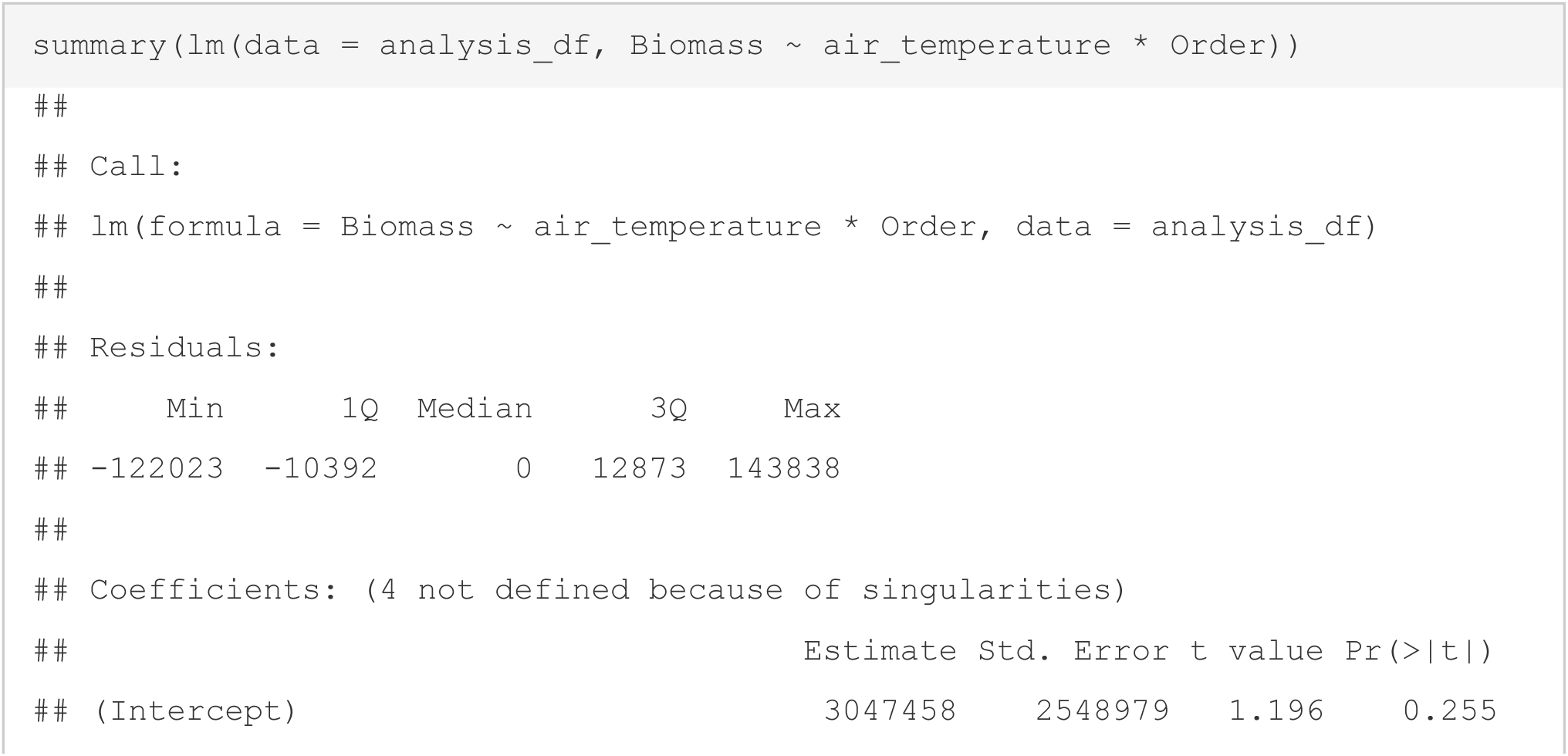
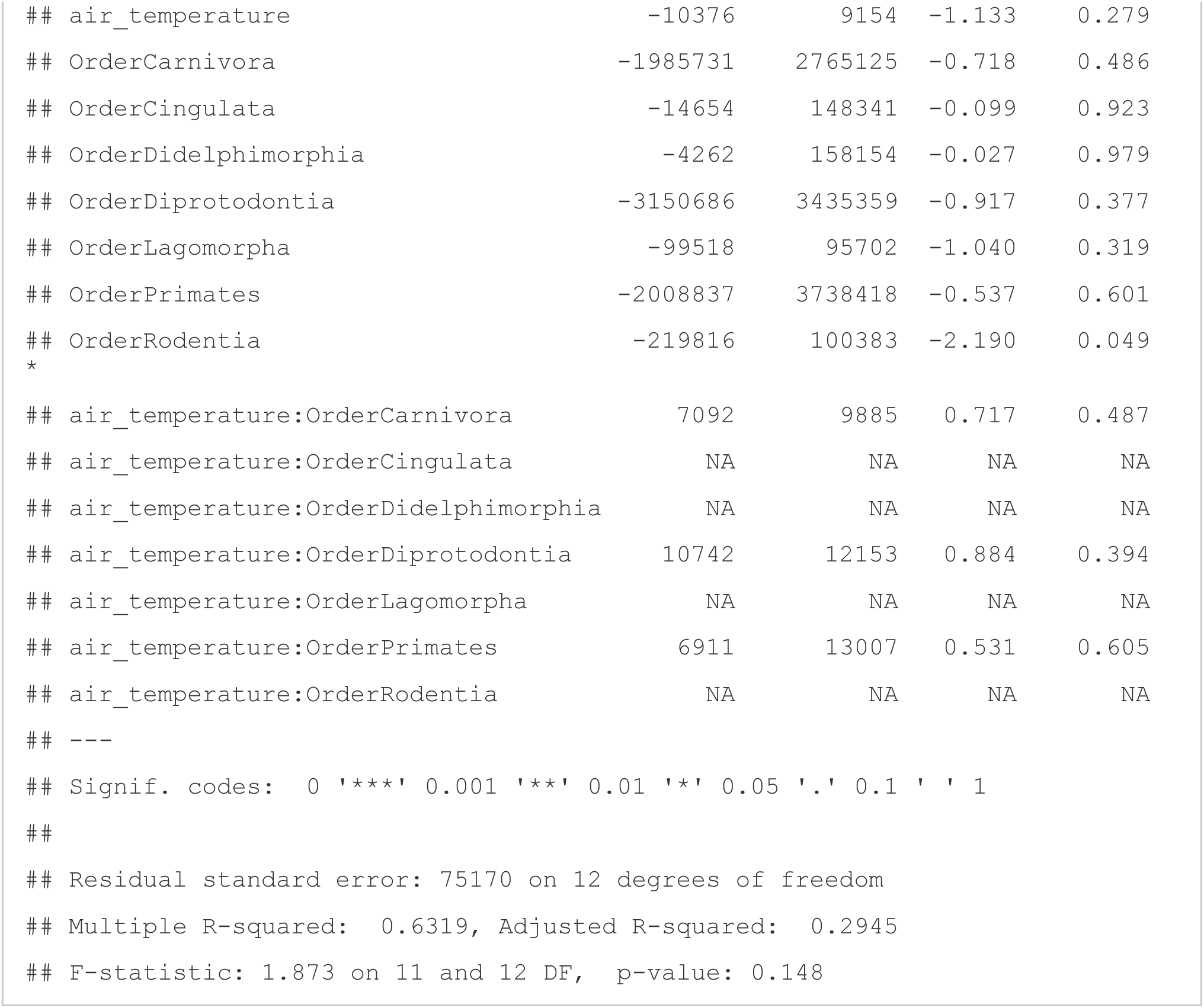

`Well, this is just a mess, but should serve to illustrate how many different analyses can easily be performed with MOSAIC and the in-built climate parameters.`

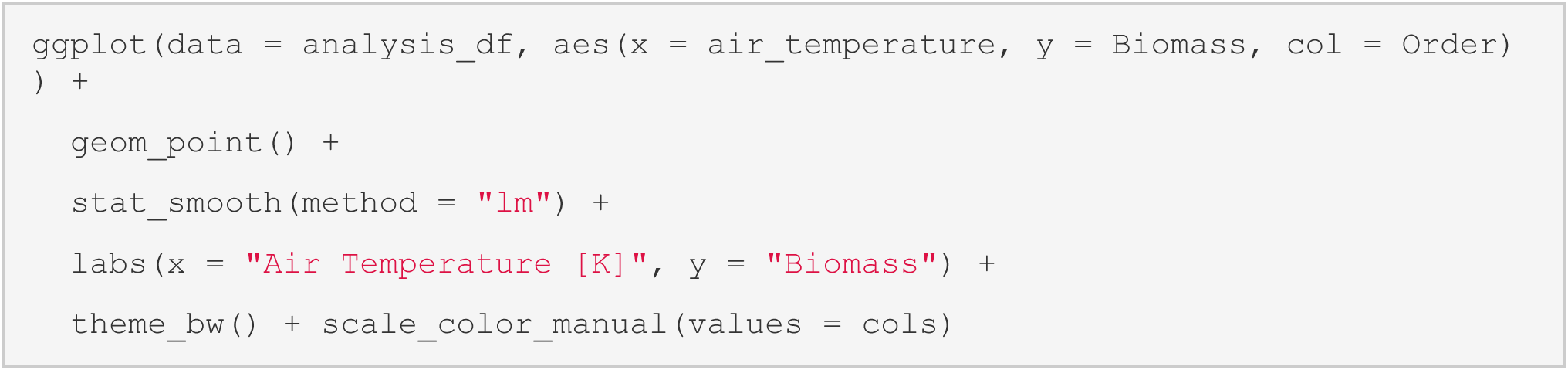

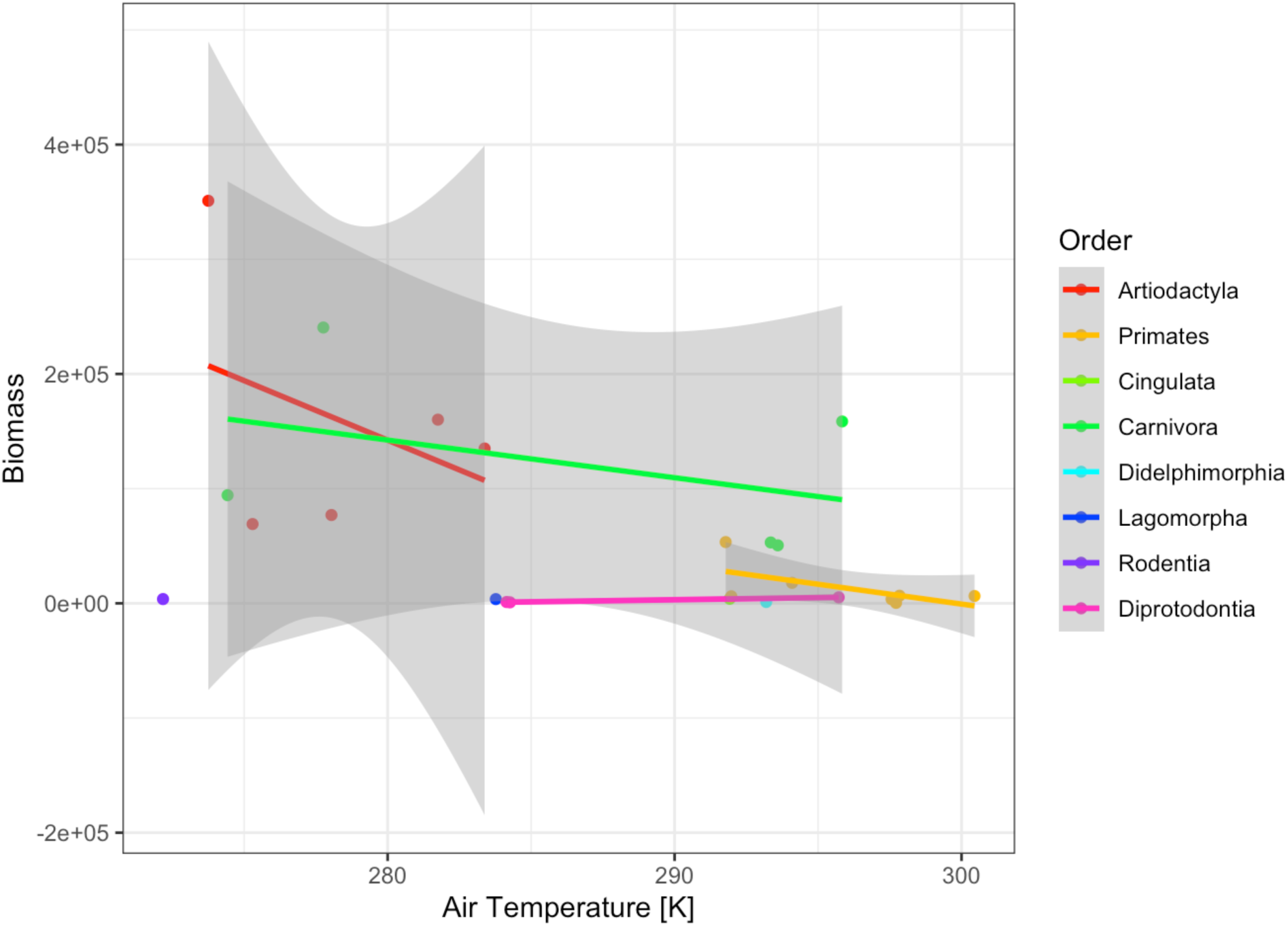

## S5: LogNormal Distribution of Mass and Height

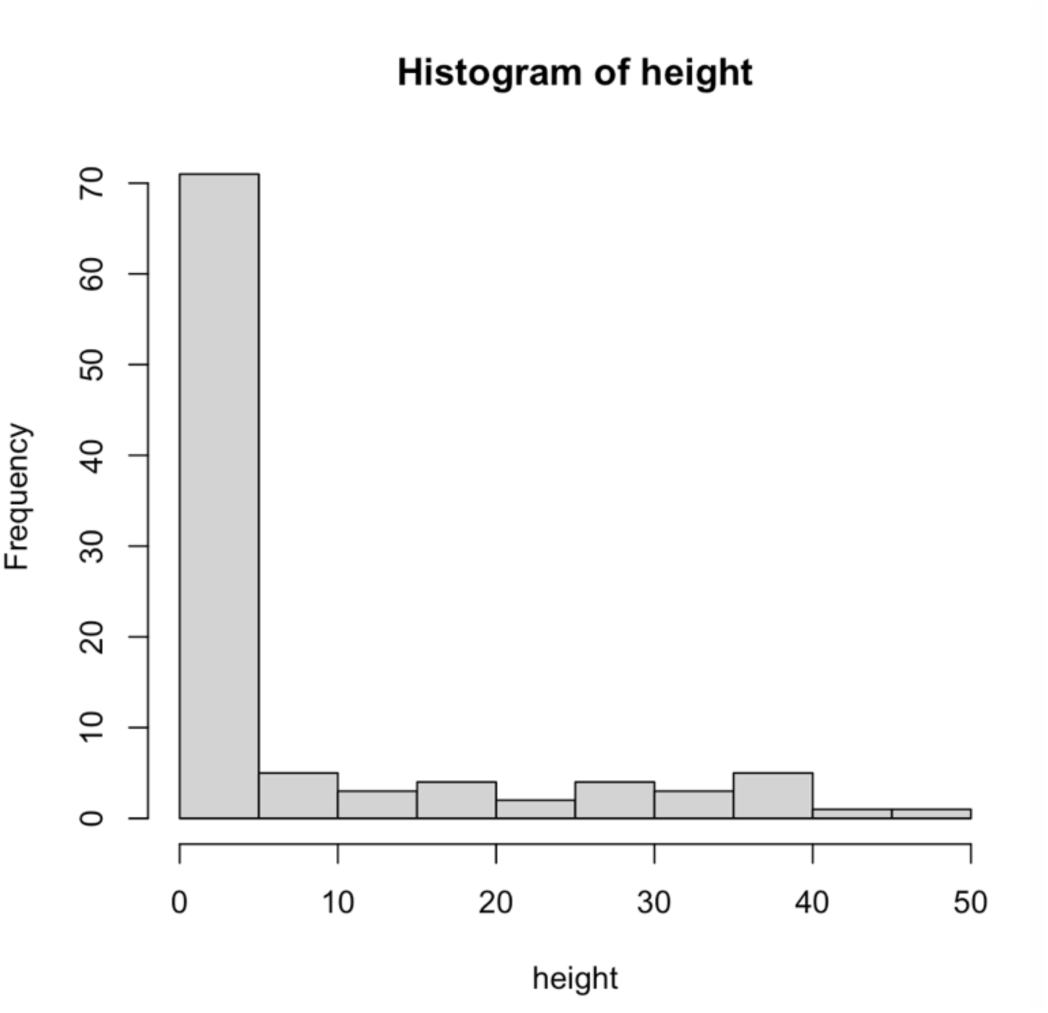

Raw heights from MOSAIC v1.0.0

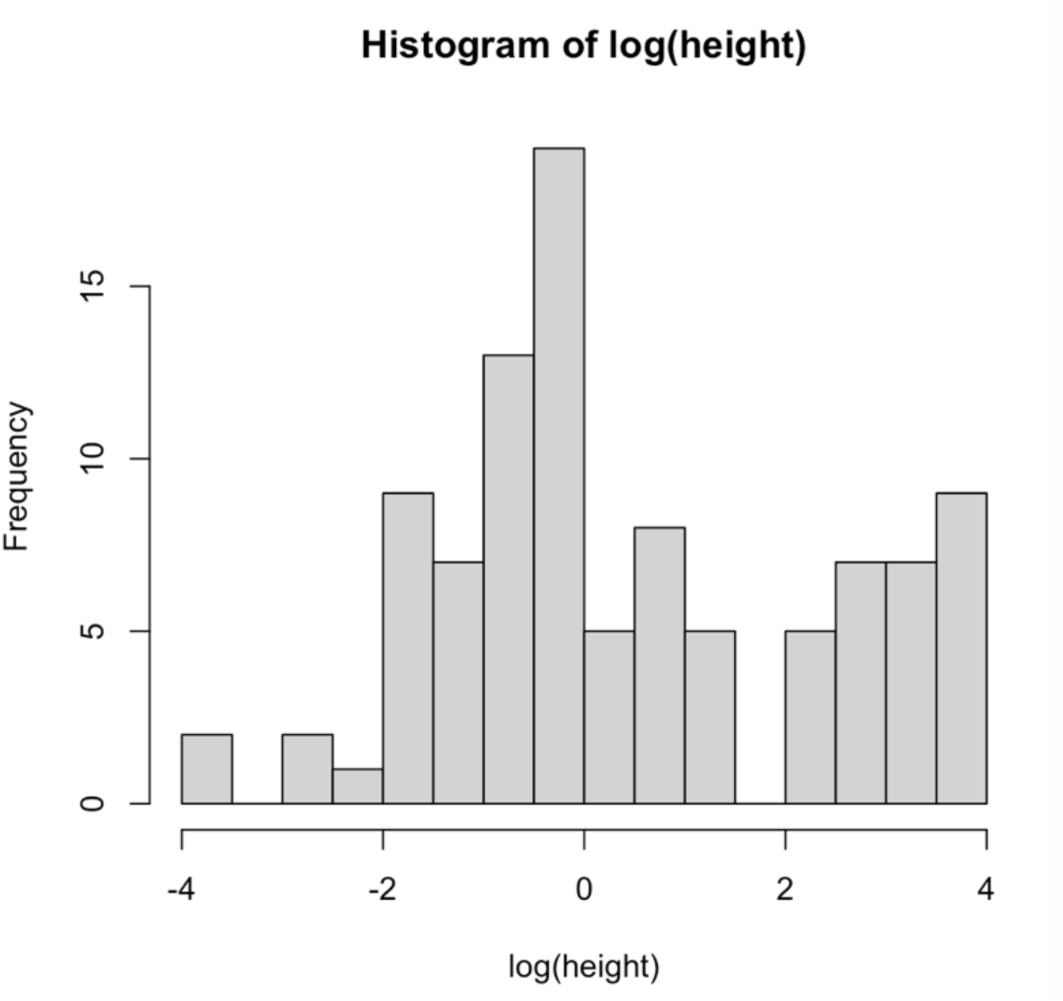

Log Heights from MOSAIC v.1.0.0

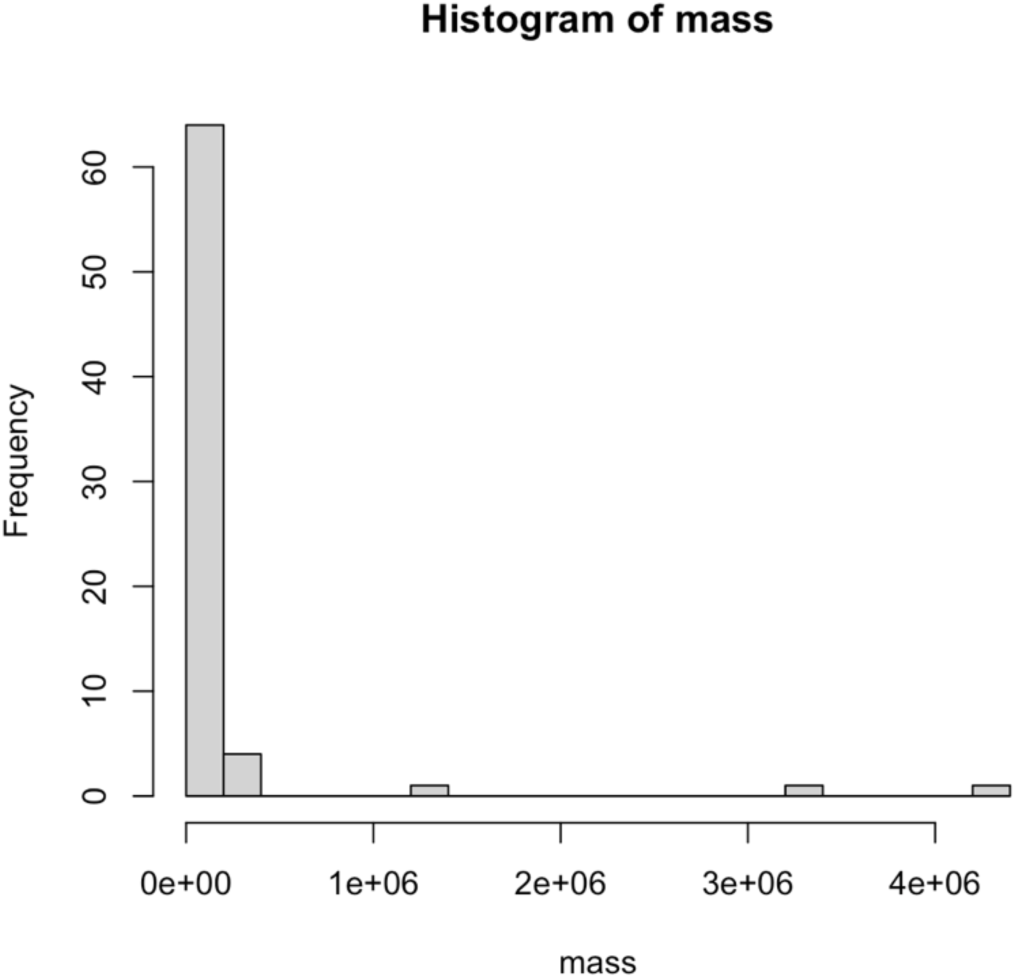

Raw mass from MOSAIV v1.0.0

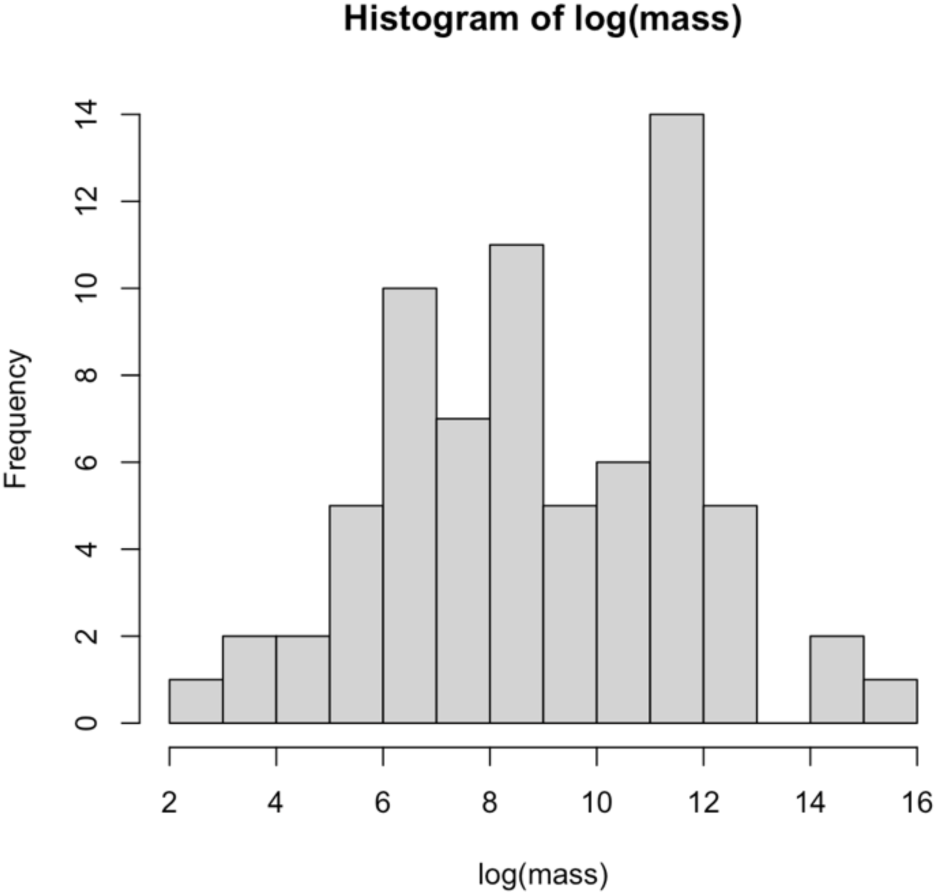

Log mass from MOSAIC v1.0.0

## References

Adler, P. B., Salguero-Gómez, R., Compagnoni, A., Hsu, J. S., Ray-Mukherjee, J., Mbeau-Ache, C., & Franco, M. (2014). Functional traits explain variation in plant life history strategies. Proceedings of the National Academy of Sciences of the United States of America, 111(27), 10019. doi: 10.1073/pnas.1410430111

Albert, C. H. (2015). Intraspecific trait variability matters. Journal of Vegetation Science, 26(1), 7–8. doi: 10.1111/jvs.12240

Albert, C.H., Thuiller, W., Yoccoz, N.G., Soudant, A., Boucher, F., Saccone, P. & Lavorel, S. (2010) Intraspecific functional variability: extent, structure and source of variation. The Journal of Ecology, 98, 604–613.

Bellier, E., Kéry, M., & Schaub, M. (2018). Relationships between vital rates and ecological traits in an avian community. Journal of Animal Ecology, 87(4), 1172–1181. doi: 10.1111/1365-2656.12826

Boettiger, C., & Temple Lang, D. (2012). Treebase: An R package for discovery, access and manipulation of online phylogenies. Methods in Ecology and Evolution, 3(6), 1060–1066. doi: 10.1111/j.2041-210X.2012.00247.x

Bolnick, D.I., Amarasekare, P., Araujo, M.S., Burger, R., Levine, J.M., Novak, M. et al. (2011). Why intraspecific trait variation matters in community ecology. Trend Ecol. Evol., 24, 183– 192.

Borghetti, M., Gentilesca, T., Colangelo, M., Ripullone, F., & Rita, A. (2020). Xylem Functional Traits as Indicators of Health in Mediterranean Forests. Current Forestry Reports, 6(3), 220– 236. doi: 10.1007/s40725-020-00124-5

Buckley, Y. M., & Puy, J. (2022). The macroecology of plant populations from local to global scales. New Phytologist, 233(3), 1038–1050. doi: 10.1111/nph.17749

Capdevila, P., Beger, M., Blomberg, S. P., Hereu, B., Linares, C., & Salguero-Gómez, R. (2020). Longevity, body dimension and reproductive mode drive differences in aquatic versus terrestrial life-history strategies. Functional Ecology, 34(8), 1613–1625. doi: 10.1111/1365-2435.13604

Carmona, C. P., de Bello, F., Azcarate, F. M., Mason, N. W. H., & Peco, B. (2019). Trait hierachies and intraspecific variability drive competitive interactions in Mediterranean annual plants. Journal of Ecology, 107, 2078–2089. https://doi.org/10.1111/1365-2745.13248

Caswell, H. (2001). Matrix population models (2nd editio). Sunderland, MA: Sinauer.

Chalmandrier, L., Hartig, F., Laughlin, D. C., Lischke, H., Pichler, M., Stouffer, D. B., & Pellissier, L. (2021). Linking functional traits and demography to model species-rich communities. Nature Communications, 12(1). doi: 10.1038/s41467-021-22630-1

Cheruvelil, K. S., & Soranno, P. A. (2018). Data-intensive ecological research is catalyzed by open science and team science. BioScience, 68(10), 813–822. doi: 10.1093/biosci/biy097

Conde, D. A., Staerk, J., Colchero, F., da Silva, R., Schöley, J., Maria Baden, H., … Vaupel, J. W. (2019). Data gaps and opportunities for comparative and conservation biology. Proceedings of the National Academy of Sciences of the United States of America, 116(19), 9658–9664. doi: 10.1073/pnas.1816367116

Consortium, E. P. (2012). An integrated encyclopedia of DNA elements in the human genome. Nature, 489(7414), 57–74. doi: 10.1038/nature11247

Crouse, D. T., Crowder, L. B., & Caswell, H. (1987). A stage-based population model for loggerhead sea turtles and implications for conservation. Ecology, 68(5), 1412–1423.

Csergő, A. M., Salguero-Gómez, R., Broennimann, O., Coutts, S. R., Guisan, A., Angert, A. L., … Buckley, Y. M. (2017). Less favourable climates constrain demographic strategies in plants. Ecology Letters, 20(8), 969–980. doi: 10.1111/ele.12794

Culina, A., Baglioni, M., Crowther, T. W., Visser, M. E., Woutersen-Windhouwer, S., & Manghi, P. (2018). Navigating the unfolding open data landscape in ecology and evolution. Nature Ecology and Evolution, 2(3), 420–426. doi: 10.1038/s41559-017-0458-2

Daskalova, G. N., Bowler, D., Myers-Smith, I. H., & Dornelas, M. (2021). Representation of global change drivers across biodiversity datasets. EcoEvoRxiv, 1–36. Retrieved from https://doi.org/10.32942/osf.io/db4s7

Daskalova, G. N., Myers-Smith, I. H., & Godlee, J. L. (2020). Rare and common vertebrates span a wide spectrum of population trends. Nature Communications, 11(1), 1–13. doi: 10.1038/s41467-020-17779-0

Davy, R., & Kusch, E. (2021). Reconciling high resolution climate datasets using KrigR. Environmental Research Letters.

De Magalhᾶes, J. P., & Costa, J. (2009). A database of vertebrate longevity records and their relation to other life-history traits. Journal of Evolutionary Biology, 22(8), 1770–1774. doi: 10.1111/j.1420-9101.2009.01783.x

Díaz, S., Kattge, J., Cornelissen, J. H. C., Wright, I. J., Lavorel, S., Dray, S., … Gorné, L. D. (2016). The global spectrum of plant form and function. Nature, 529(7585), 167–171. doi: 10.1038/nature16489

Diepenbroek, M., Glöckner, F. O., Grobe, P., Güntsch, A., Huber, R., König-Ries, B., … Triebel, D. (2014). Towards an Integrated Biodiversity and Ecological Research Data Management and Archiving Platform : The …. *Informatik 2014 – Big Data Komplexität Meistern. GI-Edition: Lecture Notes in Informatics (LNI) – Proceedings 232*, (November), 1711–1721. Retrieved from https://dl.gi.de/handle/20.500.12116/2782

Dornelas, M., Antão, L. H., Moyes, F., Bates, A. E., Magurran, A. E., Adam, D., … Zettler, M. L. (2018). BioTIME: A database of biodiversity time series for the Anthropocene. Global Ecology and Biogeography, 27(7), 760–786. doi: 10.1111/geb.12729

Easterling, M. R., Ellner, S. P., & Dixon, P. M. (2000). Size-specific sensitivity: Applying a new structured population model. Ecology, 81(3), 694–708.

Edwards, J. L., Lane, M. A., & Nielsen, E. S. (2000). Interoperability of biodiversity databases: Biodiversity information on every desktop. Science, 289(5488), 2312–2314. doi: 10.1126/science.289.5488.2312

Enquist, B. J., Condit, R., Peet, R. K., Schildhauer, M., & Thiers, B. (2016). The Botanical Information and Ecology Network (BIEN): Cyberinfrastructure for an integrated botanical information network to investigate the ecological impacts of global climate change on plant biodiversity. PeerJ Preprints.

Enquist, B. J., Feng, X., Boyle, B., Maitner, B., Newman, E. A., Jørgensen, P. M., … McGill, B. J. (2019). The commonness of rarity: Global and future distribution of rarity across land plants. Science Advances, 5(11), 1–14. doi: 10.1126/sciadv.aaz0414

Enquist, B. J., Norberg, J., Bonser, S. P., Violle, C., Webb, C. T., Henderson, A., … Savage, V. (2015). Scaling from traits to ecosystems: developing a general trait driver theory via integrating trait-based and metabolic scaling theories. In *Advances in ecological research* (pp. 249–318).

Farley, S. S., Dawson, A., Goring, S. J., & Williams, J. W. (2018). Situating ecology as a big-data science: Current advances, challenges, and solutions. BioScience, 68(8), 563–576. doi: 10.1093/biosci/biy068

Field, D., Sansone, S., Collis, A., Booth, T., Dukes, P., Gregurick, S. K., … Remington, K. (2009). ’Omics Data Sharing. Science, 326, 234–236.

Fortuna, M. A., Ortega, R., & Bascompote, J. (2014). The web of life. ArXiv. doi: 10.4324/9780203410134

Froese, R., & Pauly, D. (2010). *FishBase*. Retrieved from http://www.ices.dk/sites/pub/CMDoccuments/1992/L/1992_L10.pdf

Gallagher, R. V., Falster, D. S., Maitner, B. S., Salguero-Gómez, R., Vandvik, V., Pearse, W. D., … Enquist, B. J. (2020). Open Science principles for accelerating trait-based science across the Tree of Life. Nature Ecology and Evolution, 4(3), 294–303. doi: 10.1038/s41559-020-1109-6

Guerrero-Ramírez, N. R., Mommer, L., Freschet, G. T., Iversen, C. M., McCormack, M. L., Kattge, J., … Weigelt, A. (2021). Global root traits (GRooT) database. Global Ecology and Biogeography, 30(1), 25–37. doi: 10.1111/geb.13179

Healy, K., Guillerme, T., Finlay, S., Kane, A., Kelly, S. B. A., McClean, D., … Cooper, N. (2014). Ecology and mode-of-life explain lifespan variation in birds and mammals. Proceedings of the Royal Society B: Biological Sciences, 281(1784). doi: 10.1098/rspb.2014.0298

Hersbach, H., Bell, B., Berrisford, P., Hirahara, S., Horányi, A., Muñoz-Sabater, J., … Thépaut, J. M. (2020). The ERA5 global reanalysis. Quarterly Journal of the Royal Meteorological Society, 146(730), 1999–2049. doi: 10.1002/qj.3803

J.R. Wilmoth, K. Andreev, D. Jdanov, and D. A. G. (2007). Methods Protocol for the Human Mortality Database. Database, 2007(June 2000), 1–80.

Jasilioniene, A., Jdanov, D. A., Sobotka, T., Andreev, E. M., Zeman, K., & Shkolnikov, V. M. (2015). Methods Protocol for the Human Fertility Database. In *Max Planck Institute for Demographic Research*. Retrieved from http://www.humanfertility.org/Docs/methods.pdf

Jeliazkov, A., Mijatovic, D., Chantepie, S., Andrew, N., Arlettaz, R., Barbaro, L., … Chase, J. M. (2020). A global database for metacommunity ecology, integrating species, traits, environment and space. Scientific Data, 7(1), 1–15. doi: 10.1038/s41597-019-0344-7

Jetz, W., Thomas, G. H., Joy, J. B., Hartmann, K., & Mooers, A. O. (2012). The global diversity of birds in space and time. Nature, 491(7424), 444–448. doi: 10.1038/nature11631

Jones, K. E., Bielby, J., Cardillo, M., Fritz, S. A., O’Dell, J., Orme, C. D. L., … Purvis, A. (2009). PanTHERIA: a species-level database of life history, ecology, and geography of extant and recently extinct mammals. Ecology, 90(9), 2648–2648. doi: 10.1890/08-1494.1

Jongejans, E., Shea, K., Skarpaas, O., Kelly, D., & Ellner, S. P. (2011). Importance of individual and environmental variation for invasive species spread: A spatial integral projection model. Ecology, 92(1), 86–97. doi: 10.1890/09-2226.1

Kattge, J., Bönisch, G., Díaz, S., Lavorel, S., Prentice, I. C., Leadley, P., … Wirth, C. (2020). TRY plant trait database – enhanced coverage and open access. Global Change Biology, 26(1), 119–188. doi: 10.1111/gcb.14904

Kissling, W. D., Walls, R., Bowser, A., Jones, M. O., Kattge, J., Agosti, D., … Guralnick, R. P. (2018). Towards global data products of Essential Biodiversity Variables on species traits. Nature Ecology and Evolution, 2(10), 1531–1540. doi: 10.1038/s41559-018-0667-3

Laughlin, D. C. (2018). Rugged fitness landscapes and Darwinian demons in trait-based ecology. New Phytologist, 217(2), 501–503. doi: 10.1111/nph.14908

Laughlin, D. C., Gremer, J. R., Adler, P. B., Mitchell, R. M., & Moore, M. M. (2020). The Net Effect of Functional Traits on Fitness. Trends in Ecology and Evolution, 35(11), 1037–1047. doi: 10.1016/j.tree.2020.07.010

Levin, S. C., Crandall, R. M., Pokoski, T., Stein, C., & Knight, T. M. (2020). Phylogenetic and functional distinctiveness explain alien plant population responses to competition: Phylogeny and traits explain dominance. Proceedings of the Royal Society B: Biological Sciences, 287(1930). doi: 10.1098/rspb.2020.1070rspb20201070

Lintulaakso, K. (2013). MammalBase—database of recent mammals.

Madin, J. S., Anderson, K. D., Andreasen, M. H., Bridge, T. C. L., Cairns, S. D., Connolly, S. R., … Baird, A. H. (2016). The Coral Trait Database, a curated database of trait information for coral species from the global oceans. Scientific Data, 4, 170174. doi: 10.1038/sdata.2017.174

Maitner, B. S., Boyle, B., Casler, N., Condit, R., Donoghue, J., Durán, S. M., … Enquist, B. J. (2018). The bien r package: A tool to access the Botanical Information and Ecology Network (BIEN) database. Methods in Ecology and Evolution, 9(2), 373–379. doi: 10.1111/2041-210X.12861

Maldonado, C., Molina, C. I., Zizka, A., Persson, C., Taylor, C. M., Albán, J., … Antonelli, A. (2015). Estimating species diversity and distribution in the era of Big Data: To what extent can we trust public databases? Global Ecology and Biogeography, 24(8), 973–984. doi: 10.1111/geb.12326

Marx, V. (2013). The big challenges of big data. Nature, 498, 255–260.

Maurer, S. M., Firestone, R. B., & Scriver, C. R. (2000). Science’s neglected legacy: Large, sophisticated databases cannot be left to chance and improvisation. Nature, 405(6783), 117– 120. doi: 10.1038/35012169

McElreath, R. (2021) Statistical Rethinking. London, UK: Chapman and Hall/CRC.

Messier, J., McGill, B. J., & Lechowicz, M. J. (2010). How do traits vary across ecological scales? A case for trait-based ecology. Ecology Letters, 13(7), 838–848. doi: 10.1111/j.1461-0248.2010.01476.x

Metcalf, C. J. E., Graham, A. L., Martinez-Bakker, M., & Childs, D. Z. (2016). Opportunities and challenges of Integral Projection Models for modelling host-parasite dynamics. Journal of Animal Ecology, 85(2), 343–355. doi: 10.1111/1365-2656.12456

Michener, W. K. (2006). Meta-information concepts for ecological data management. Ecological Informatics, 1(1), 3–7. doi: 10.1016/j.ecoinf.2005.08.004

Middleton, O., Svensson, H., Scharlemann, J. P. W., Faurby, S., & Sandom, C. (2021). CarniDIET 1.0: A database of terrestrial carnivorous mammal diets. Global Ecology and Biogeography, 30(6), 1175–1182. doi: 10.1111/geb.13296

Monnet, A. C., Cilleros, K., Médail, F., Albassatneh, M. C., Arroyo, J., Bacchetta, G., … Leriche, A. (2021). WOODIV, a database of occurrences, functional traits, and phylogenetic data for all Euro-Mediterranean trees. Scientific Data, 8(1), 1–11. doi: 10.1038/s41597-021-00873-3

Morris, W. F., Pfister, C. A., Tuljapurkar, S., Haridas, C. V., Boggs, C. L., Boyce, M. S., … Menges, E. S. (2008). Longevity can buffer plant and animal populations against chnaging climatic variability. Ecology, 89(1), 19–25. Retrieved from http://www.esajournals.org/doi/pdf/10.1890/07-0774.1

Myhrvold, N. P., Baldridge, E., Chan, B., Sivam, D., Freeman, D. L., & Ernest, S. K. M. (2015). An amniote life-history database to perform comparative analyses with birds, mammals, and reptiles. Ecology, 96(11). doi: 10.1890/15-0846r.1

Nadrowski, K., Ratcliffe, S., Bönisch, G., Bruelheide, H., Kattge, J., Liu, X., … Wirth, C. (2013). Harmonizing, annotating and sharing data in biodiversity-ecosystem functioning research. Methods in Ecology and Evolution, 4(2), 201–205. doi: 10.1111/2041-210x.12009

Oliveira, B. F., São-Pedro, V. A., Santos-Barrera, G., Penone, C., & Costa, G. C. (2017). AmphiBIO, a global database for amphibian ecological traits. Scientific Data, 4, 1–7. doi: 10.1038/sdata.2017.123

Ozgul, A., Childs, D. Z., Oli, M. K., Armitage, K. B., Blumstein, D. T., Olson, L. E., … Coulson, T. (2010). Coupled dynamics of body mass and population growth in response to environmental change. Nature, 466(7305), 482–485. doi: 10.1038/nature09210

Paniw, M., Maag, N., Cozzi, G., Clutton-Brock, T., & Ozgul, A. (2019). Life history responses of meerkats to seasonal changes in extreme environments. Science, 363(6427), 631–635. doi: 10.1126/science.aau5905

Pistón, N., de Bello, F., Dias, A. T. C., Götzenberger, L., Rosado, B. H. P., de Mattos, E. A., … Carmona, C. P. (2019). Multidimensional ecological analyses demonstrate how interactions between functional traits shape fitness and life history strategies. Journal of Ecology, 107(5), 2317–2328. doi: 10.1111/1365-2745.13190

Poorter, L., Wright, S.J., Paz, H., Ackerly, D.D., Condit, R., Ibarra-Manriques, G. et al. (2008) Are functional traits good predictors of demographic rates? Evidence from five Neotropical forests. Ecology, 89, 1908–1920.

Reich, P. B., Wright, I. J., Cavender-Bares, J., Craine, J. M., Oleksyn, J., Westoby, M., & Walters, M. B. (2003). The evolution of plant functional variation: Traits, spectra, and strategies. International Journal of Plant Sciences, 164(SUPPL. 3). doi: 10.1086/374368

Reichman, O. J., Jones, M. B., & Schildhauer, M. P. (2011). Challenges and opportunities of open data in ecology. Science, 331(6018), 703–705. doi: 10.1126/science.1197962

Roper, M., Capdevila, P., & Salguero-Gómez, R. (2021). Senescence: Why and where selection gradients might not decline with age. Proceedings of the Royal Society B: Biological Sciences, 288(1955). doi: 10.1098/rspb.2021.0851

Rowcliffe, J. M., Jansen, P. A., Kays, R., Kranstauber, B., & Carbone, C. (2016). Wildlife speed cameras: measuring animal travel speed and day range using camera traps. Remote Sensing in Ecology and Conservation, 2(2), 84–94. doi: 10.1002/rse2.17

Salguero-Gómez, R., Jackson, J., & Gascoigne, S.J.L. (2021). Four key challenges in the open data revolution. Journal of Animal Ecology, 90(*9*), 2000-2004. doi: 10.1111/1365-2656.13567

Salguero-Gómez, R., Jones, O. R., Archer, C. R., Bein, C., de Buhr, H., Farack, C., … Vaupel, J. W. (2016). COMADRE: A global data base of animal demography. Journal of Animal Ecology. doi: 10.1111/1365-2656.12482

Salguero-Gómez, R., Jones, O. R., Archer, C. R., Buckley, Y. M., Che-Castaldo, J., Caswell, H., … Vaupel, J. W. (2015). The compadre Plant Matrix Database: An open online repository for plant demography. Journal of Ecology. doi: 10.1111/1365-2745.12334

Salguero-Gómez, R., & Laughlin, D. C. (2021). Not all traits are functional: the Panglossian paradigm. *Authorea*, (December 2021). doi: 10.22541/au.163940711.10447233/v1

Salguero-Gómez, R., Violle, C., Gimenez, O., & Childs, D. (2018). Delivering the promises of trait-based approaches to the needs of demographic approaches, and vice versa. Functional Ecology, 32(6), 1424–1435. doi: 10.1111/1365-2435.13148

Santini, L., Isaac, N. J. B., & Ficetola, G. F. (2018). TetraDENSITY: A database of population density estimates in terrestrial vertebrates. Global Ecology and Biogeography, 27(7), 787– 791. doi: 10.1111/geb.12756

Schneider, F. D., Fichtmueller, D., Gossner, M. M., Güntsch, A., Jochum, M., König-Ries, B., … Simons, N. K. (2019). Towards an ecological trait-data standard. Methods in Ecology and Evolution, 10(12), 2006–2019. doi: 10.1111/2041-210X.13288

Siefert, A., Violle, C., Chalmandrier, L., Albert, C. H., Taudiere, A., Fajardo, A., … Wardle, D. A. (2015). A global meta-analysis of the relative extent of intraspecific trait variation in plant communities. Ecology Letters, 18(12), 1406–1419. doi: 10.1111/ele.12508

Smallegange, I. M., Deere, J. A., & Coulson, T. (2014). Correlative changes in life-history variables in response to environmental change in a model organism. American Naturalist, 183(6), 784–797. doi: 10.1086/675817

Stenseth, N. C., & Mysterud, A. (2002). Climate, changing phenology, and other life history traits: Nonlinearity and match-mismatch to the environment. Proceedings of the National Academy of Sciences of the United States of America, 99(21), 13379–13381. doi: 10.1073/pnas.212519399

Swenson, N. G., Worthy, S. J., Eubanks, D., Iida, Y., Monks, L., Petprakob, K., … Zambrano, J. (2020). A reframing of trait–demographic rate analyses for ecology and evolutionary biology. International Journal of Plant Sciences, 181(1), 33–43. doi: 10.1086/706189

Terry, J. C. D., O’Sullivan, J. D., & Rossberg, A. G. (2022). No pervasive relationship between species size and local abundance trends. Nature Ecology and Evolution, 6(2), 140–144. doi: 10.1038/s41559-021-01624-8

Troia, M. J., & McManamay, R. A. (2016). Filling in the GAPS: evaluating completeness and coverage of open-access biodiversity databases in the United States. Ecology and Evolution, 6(14), 4654–4669. doi: 10.1002/ece3.2225

Violle, C., Enquist, B. J., McGill, B. J., Jiang, L., Albert, C. H., Hulshof, C., … Messier, J. (2012). The return of the variance: Intraspecific variability in community ecology. Trends in Ecology & Evolution, 27, 244–252.

Violle, C., Navas, M.-L., Vile, D., Kazakou, E., Fortunel, C., Hummel, I., & Garnier, E. (2007). Let the concept of trait be functional! Oikos, 116(5), 882–892. doi: 10.1111/j.2007.0030-1299.15559.x

Vitousek, M. N., Johnson, M. A., & Husak, J. F. (2018). Illuminating endocrine evolution: The power and potential of large-scale comparative analyses. Integrative and Comparative Biology, 58(4), 712–719. doi: 10.1093/icb/icy098

Whitlock, M. C. (2011). Data archiving in ecology and evolution: Best practices. Trends in Ecology and Evolution, 26(2), 61–65. doi: 10.1016/j.tree.2010.11.006

Wieczorek, J., Bloom, D., Guralnick, R., Blum, S., Döring, M., Giovanni, R., … Vieglais, D. (2012). Darwin core: An evolving community-developed biodiversity data standard. PLoS ONE, 7(1). doi: 10.1371/journal.pone.0029715

Wilkes, M. A., Edwards, F., Jones, J. I., Murphy, J. F., England, J., Friberg, N., … Brown, L. E. (2020). Trait-based ecology at large scales: Assessing functional trait correlations, phylogenetic constraints and spatial variability using open data. Global Change Biology, 26(12), 7255–7267. doi: 10.1111/gcb.15344

Wilkinson, M. D., Dumontier, M., Aalbersberg, Ij. J., Appleton, G., Axton, M., Baak, A., … Mons, B. (2016). Comment: The FAIR Guiding Principles for scientific data management and stewardship. Scientific Data, 3, 1–9. doi: 10.1038/sdata.2016.18

Williams, N. F., McRae, L., Freeman, R., Capdevila, P., & Clements, C. F. (2021). Scaling the extinction vortex: Body size as a predictor of population dynamics close to extinction events. Ecology and Evolution, 11(11), 7069–7079. doi: 10.1002/ece3.7555

Yang, J., Cao, M., & Swenson, N. G. (2018). Why Functional Traits Do Not Predict Tree Demographic Rates. Trends in Ecology and Evolution, 33(5), 326–336. doi: 10.1016/j.tree.2018.03.003

