## Supplementary material for "MOSAIC: A Unified Trait Database to Complement Structured Population Models": Online Supplemental Material

#### SUPPLEMENTARY ONLINE MATERIALS

S1: MOSAIC User Guide

### USER GUIDE TO THE MOSAIC LIFE HISTORY DATABASE

A Companion to COM(P)ADRE and PADRINO/A demographic databases

*Working document – 06 February 2022*

Table of Contents

User Guide Information

User Guide

Appendix

References

Table of Contents

**Introduction**
General Instructions
Database Organization Page 6
Database Design Page 6
The meanings of NA in MOSAIC Page 6
Disclaimer Page 6
What is new in this version? Page 7

Format of the User Guide
Format Diagram Page 8
Variables in MOSAIC Page 9

**Metadata**
A. Species Name/Taxonomy
1. **A1** Species Accepted Page 10
2. **A2** Kingdom Page 11

B. Study Information
3. **B1** Author Page 12
4. **B2** Journal Name Page 13
5. **B3** Year Publication Page 14
6. **B4** DOI/ISBN Page 15

**Primary User Guide/Variables**
* Applicable to plants
** Applicable to animals

A. Morphometry & Growth
1. **A1** Biomass Page 17
2. **A2** Height Page 18
3. **A3** Growth Determination Page 19
4. **A4** Regeneration Page 20
5. **A5** Sexual Dimorphism Page 21

B. Reproductive traits
6. **B1** Mating System** Page 22
7. **B2** Hermaphrodism Page 23
8. **B3** Protogyny/Protandry** Page 24

C. Movement traits
9. **C1** Dispersal Capability Page 25
10. **C2** Type of Dispersal Page 26
11. **C3** Mode of Dispersal Page 27
12. **C4** Dispersal Class Page 28
13. **C5** Volancy** Page 29

**What is new in this version?**

Version 1.0.0

- The first version of the database. No updates.

---

DIAGRAM OF THE MOSAIC DATABASE ARCHITECTURE

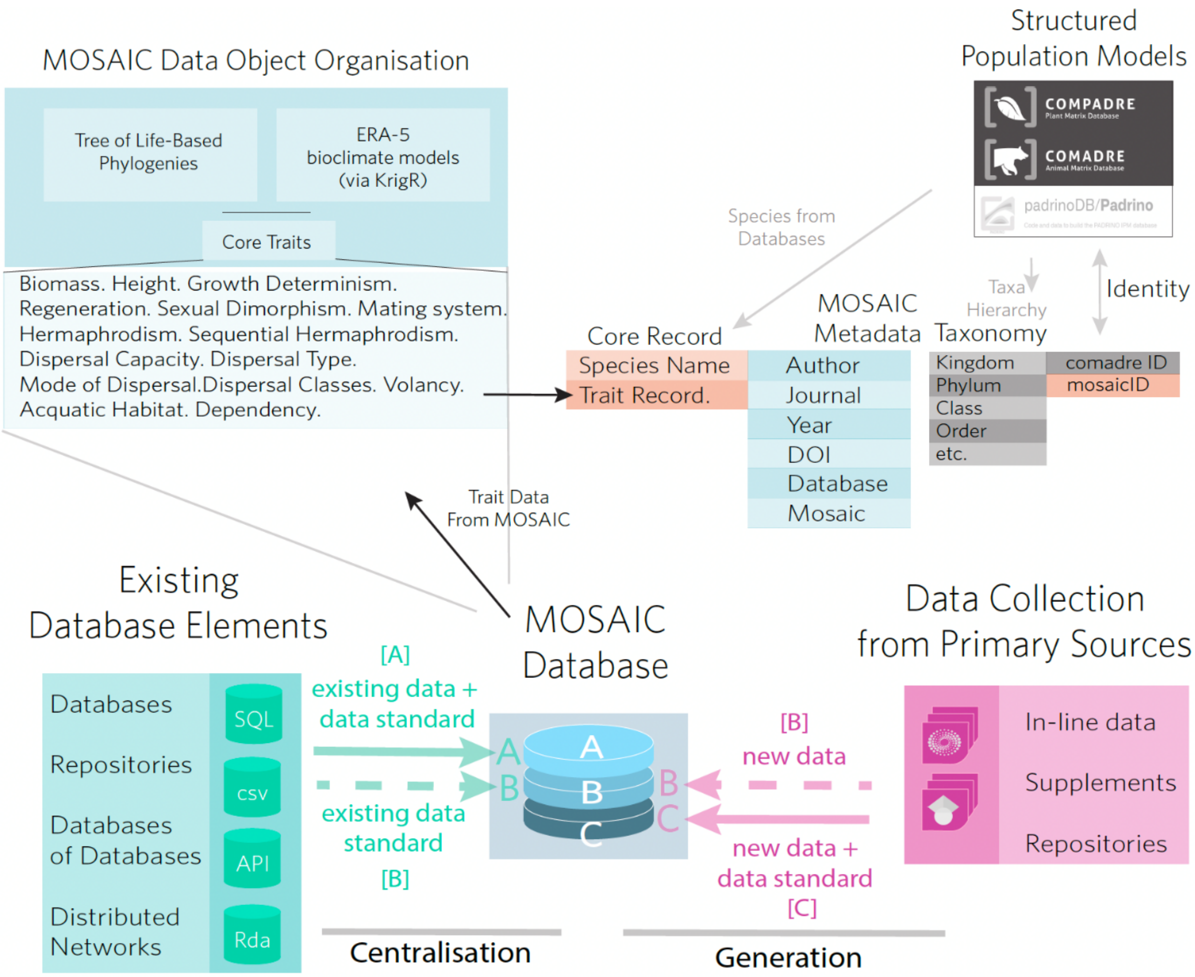

**Variables in MOSAIC**

The MOSAIC database is constructed of objects containing life history information across variables and organized into themes to aid in navigation. The metadata object is the object containing information about every study for which data is stored in the MOSAIC database. Every variable containing a value in the MOSAIC database will have corresponding metadata.

Optional clearance of working space.

```
rm(list = ls()) # Clear your environment
if(!is.null(dev.list())) dev.off() # Clear plots/graphics
cat("\014") # Clear console
```

##### Downloading MOSAIC

MOSAIC can be downloaded as an S4 object by running the below code in R. S4 data objects in R are an object oriented system in the R language that allow control of constituent data fields Similar to S3 objects (which use the “$” operator). S4 are comprised of objects that can be searched with the “@” operator or slots, discussed in more detail below.

```
library(devtools) # Compulsory package to pull down packages from GitHub. Ins
tall if necessary.
install_github("mosaicdatabase/Rmosaic")
source_url("https://raw.githubusercontent.com/mosaicdatabase/mosaicdatabase/m
ain/mosaic_fetch.R") # Link to GitHub repo
mosaic <- mos_fetch("v1.0.0") # Download version 1.0.0 (active version Feb 20
22)
library(Rmosaic)
```

##### Basics of manually navigating MOSAIC

Mosaic traits can be searched using the “@” operator. Attribute names searched this way are analogous to the columns of a dataframe in a relational database structure.

Once downloaded, you should be able to type statements mosaicdatabase@[insertfield] (where [insertfield] is a particular trait). If you are working in Rstudio, after the "@" a drop-down of the slots (traits) should autopopulate.

```
# Three examples of querying traits (easiest for navigation)
mosaic@biomass
mosaic@height
mosaic@volancy
```

The species corresponding with each index can be queried by prompting:

```
# Three examples of querying traits (easiest for navigation)
mosaic@species
```

Data in mosaic can also be access using slots. Slots are the recommended mode of searching the database - though it has the disadvantage of not enabling the autopopulation of the attributes contained in the database (traits must be spelled out manually).

```
# You can also search these by slot (recommended)
slot(mosaic, "biomass")
slot(mosaic, "height")
slot(mosaic, "volancy")
```

Within each trait object, there are eight fields in mosaic. The first of field is called "values" and contains the data. Values are unitless values (either numeric or factorial) that are reported in units described in the **Mosaic User Guide** http://mosaicdatabase.web.ox.ac.uk/user-guide. The metadata is organised into additional attributes, reflecting the individual elements of the metadata for a given record, including the authors, journal, year of publication, databases from which data are sourced (if applicable)

mosaic@metaTaxa maps the complete taxanomic classification structure of a species - from Kingdom to species for taxonomic clustering.

The other six attributes - "author", "year", "journal,"doi","database", and"mosaic" - are metadata corresponding with each value record. For instance:

```
mosaic@species[[2]] # For this species, let us look at volancy (flight capaci
ty) value
mosaic@volancy@value[[2]] # or, equivalently:
slot(slot(mosaic, "volancy"), "value")[[2]]
```

Corresponds with the following metadata

```
mosaic@volancy@author[[2]] # for "Acinonyx jubatus"
```

```
slot(slot(mosaic, "volancy"), "author")[[2]] # Author of the source publicati
on
slot(slot(mosaic, "volancy"), "year")[[2]] # And year of the source publicati
on
slot(slot(mosaic, "volancy"), "journal")[[2]] # The journal of the source pub
lication
#etc.
```

##### Using MOSAIC functions to quickly access files

A series of convenience functions can be sourced from the MOSAIC GitHub page to facilitate navigating and working with the mosaic database that can be accessed by running the follwing script.

```
source_url("https://raw.githubusercontent.com/mosaicdatabase/mosaicdatabase/m
ain/navMosaic_46.R")
```

Below we highlight some of the basic queries for which the mosaic functions can assist.

##### Is a species included in Mosaic?

```
spp_check("Fritillaria biflora")
## [1] TRUE
spp_check("Pagophilus groenlandicus")
## [1] FALSE
```

##### Can I see all records for a given trait?

```
traitAllSpp("biomass") # Only the first five records are shown for space
## [1] "NDY"    "50578"  "ND"     "52500"  "351000" "62000"
```

##### Can I see an overview of all records for a given species?

```
singSppTraitSummary("Aepyceros melampus")
##   biomass height   growthdet regen   dimorph        matsyst     hermaph se
qherm
## 1   52500    NDY Determinate   NDY Dimorphic Non-monogamous Gonochorous
NDY
##           dispcap disptype modedisp dispclass    volancy
## 1 Natal Dispersal   Active   Motile     Adult Non-volant
```

```
##                                  aquadep
## 1 Terrestrial, Water Habitat Independent
```

Can I see all records for more than one species?

```
sppAllTrait(c("Acinonyx jubatus", # you can also pass lists or dataframes to
this command
               "Acropora downingi",
               "Aepyceros melampus",
               "Alces alces",
               "Alligator mississippiensis"))
##                      sppnames biomass height   growthdet        regen   dim
orph
## 1           Acinonyx jubatus   50578    NDY Determinate          NDY Dimor
phic
## 2          Acropora downingi      ND    NDY         NDY Regenerative
NDY
## 3         Aepyceros melampus   52500    NDY Determinate          NDY Dimor
phic
## 4                Alces alces  351000    NDY Determinate          NDY Dimor
phic
## 5 Alligator mississippiensis   62000    NDY Determinate Regenerative
NDY
##          matsyst        hermaph seqherm         dispcap disptype       mod
edisp
## 1 Non-monogamous    Gonochorous     NDY Natal Dispersal   Active         M
otile
## 2            NDY Hermaphroditic     NDY Natal Dispersal  Passive Water cur
rents
## 3 Non-monogamous    Gonochorous     NDY Natal Dispersal   Active         M
otile
## 4 Non-monogamous    Gonochorous     NDY Natal Dispersal   Active         M
otile
## 5 Non-monogamous    Gonochorous     NDY Natal Dispersal   Active         M
otile
##           dispclass    volancy                                      aqua
dep
## 1           Juvenile Non-volant        Terrestrial, Water Habitat Independ
ent
## 2           Juvenile Non-volant                                      Mar
ine
```

```
## 3              Adult Non-volant        Terrestrial, Water Habitat Independ
ent
## 4           Juvenile Non-volant Terrestrial, Facultative Freshwater Depend
ent
## 5 Adult and Juvenile Non-volant  Terrestrial, Obligative Freshwater Depend
ent
```

Can I get a breakdown of counts/frequency of trait values?

```
traitFrequency("volancy")
##             counts  freq
## NDY           1329    NA
## Non-volant      82 0.882
## Semi-volant      1 0.011
## Volant          10 0.108
traitFrequency("growthdet")
##               counts  freq
## Determinate       31 0.596
## Indeterminate     21 0.404
## NDY             1370    NA
traitFrequency("hermaph")
##                               counts  freq
## Dioecious                         24 0.108
## Gonochorous                      106 0.475
## Hermaphroditic                    72 0.323
## Hermaphroditic & Gonochorous       1 0.004
## Monoecious                        20 0.090
## NDY                             1199    NA
```

Can I get all metadata for one or more traits?

```
metadata("volancy", 14)
##              author year                    journal database mosaic
## 1 Campos, Z. et al. 2006 The Herpetological Journal      NDY    NDY
multiMetaRecords("volancy", c(14:20))
##                                      author year
```

```
## 1                Ekerna, L. S. & Cords, M. 2007
## 2                  Zimmerman, S. J. et al. 2019
## 3 Jack, K. M., Sheller, C. & Fedigan, L. M. 2012
## 4                                      DFO 2013
## 5                     Torres, R. T. et al. 2017
## 6                                      NDY  NDY
## 7                        Campos, Z. et al. 2006
##
journal
## 1                                                                             A
nimal Behaviour
## 2
The Condor
## 3                                                              American Journal
of Primatology
## 4 Canadian Science Advisory Secretariat Central and Arctic Region Science
Advisory Report
## 5
Oryx
## 6
NDY
## 7                                                                   The Herpeto
logical Journal
##   database mosaic
## 1      NDY    NDY
## 2      NDY    NDY
## 3      NDY    NDY
## 4      NDY    NDY
## 5      NDY    NDY
## 6      NDY    NDY
## 7      NDY    NDY
```

Basic Regression using MOSAIC

http://mosaicdatabase.web.ox.ac.uk

Updated 22 February 2022

### Vignette #2 - Basic Regression Using MOSAIC & COMPADRE

MOSAIC was built with the intention of its use in comparative biodemography - to strengthen the map of which functional traits predict vital rates and which do not. Traits in the MOSAIC database (factorial and numeric variables) are formatted to be easily accessed and used for regression-based analyses, among other types of analysis. MOSAIC was also designed for quick integration with the COMADRE, COMPADRE, and PADRINO structural population model databases, as outlined in the model exercise below.

```
library(devtools)
library(tidyverse)
library(Rcompadre) # package to access matrix population models
library(Rage) # package to perform life-history calculations
```

#### Accessing MOSAIC

Download MOSAIC from the mosaic portal. For more information on the basics of downloading MOSAIC and navigating the data structure, see: Vignette #1:

```
library(devtools) # Compulsory package to pull down packages from GitHub. Ins
tall if necessary.
install_github("mosaicdatabase/Rmosaic") # library of navigation-aiding funct
ions
source_url("https://raw.githubusercontent.com/mosaicdatabase/mosaicdatabase/m
ain/mosaic_fetch.R") # Link to GitHub repo
mosaic <- mos_fetch("v1.0.0") # Download version 1.0.0 (active version Feb 20
22)
library(Rmosaic)
```

#### Regressing Biomass on Generation Time

In this exercise, the simple bivariate relationship of biomass on generation time is explored through basic regression. Bare in mind, that the below offer the building blocks for more complex multiple regression exercises and other forms of analysis – such as geospatial interpolation and ordination-based practices (see Vignette #4 for a brief vignette of PVA using mosaic).

#1: Extract biomass data for mammals

Extract the biomass values for mammals in the MOSAIC dataset.

```
# Extracting all biomass data

mosaic_dataframe <- data.frame(mosaic@taxaMetadat,
                               Biomass = as.numeric(mosaic@biomass@value),
                               MOSAIC_index = 1:length(mosaic@index))

# Restrict mammal data with non NA biomass data

mosaic_mammal <- subset(mosaic_dataframe,
                        Class == "Mammalia" & is.na(Biomass) == F)

# Extracting the first COMPADRE matrix id for each entry

matrix_ids_full <- mosaic@index[mosaic_mammal$MOSAIC_index]

# Extract just the first matrix id using sapply

mammal_matrix_ids <- sapply(matrix_ids_full, `[[`, 1)

# Add matrix id to the mammal data

mosaic_mammal$MatrixID <- mammal_matrix_ids

# Add full species names

mosaic_mammal$Binomial <- paste0(mosaic_mammal$Genus, " ", mosaic_mammal$Species)
```

#2: Download COMADRE matrix population models

A cdb_fetch() function is used to clone the Comadre database <www.compadre-db.org> - which is an s3 copy of a SQL-based relational database - into an S4 data object locally manipulable in R.

Using ids associated with each matrix population model, we will subset the Comadre database of matrix population models for the animals. First we use the Rcompadre package to ‘fetch’ the Comadre database.

```
# Download the most recent database
comadre <- cdb_fetch("comadre")
# Rcompadre function flags potential issues with matrices, including NAs and ergodicity
comadre_flag <- cdb_flag(comadre)
# Remove matrices with NAs and those that are non-ergodic (which throw errors in generation time)
comadre_correct <- subset(comadre_flag,
                          check_NA_A == FALSE &
                          check_ergodic == TRUE)
```

The subsets of mosaic and Compadre can be overlapped in one line of code.

```
# Subset to mammal matrices
comadre_biomass <- subset(comadre_correct,
                          MatrixID %in% mammal_matrix_ids)
```

3. Calculating generation time

Rcompadre enables sub-setting of a matrix population model (life table) into constituent matrices representing the survival and fecundity/reproductive components (U and F matrices, respectively). Functions in the Rage package often require the specification of both the U and the F matrix, not the combined A matrix. The Rage package contains a suite of common demographic calculations - including GENERATION TIME - which are used to simplify this exercise. Note that other common demographic derivations, including transient indices can be extracted using the identical procedure.

```
# Calculate Generation Time
comadre_biomass$Generation_Time <- mapply(Rage::gen_time,
                                          matU(comadre_biomass),
```

```
                                                    matF(comadre_biomass))
```

4. Explore the relationship of generation time data and biomass data

Matrix IDs can be used to bridge generation time from comaadre with mosaic records.

```
# Extract matrix ID and generation time columns (excluding other matrix metad
ata)

comadre_biomass <- as.data.frame(comadre_biomass)

comadre_biomass <- comadre_biomass[,c("MatrixID", "Generation_Time")]

# Join the datasets, using Matrix IDs to crosswalk the data

mosaic_biomass <- merge(mosaic_mammal, comadre_biomass,
                        by = "MatrixID", all.x = TRUE)
# Remove infinity values or NA

mosaic_biomass <- mosaic_biomass[!(is.infinite(mosaic_biomass$Generation_Time
) |
                                     is.na(mosaic_biomass$Generation_Time)),]

# Log biomass and generation time

mosaic_biomass$log10_biomass <- log10(mosaic_biomass$Biomass)
mosaic_biomass$log10_gent <- log10(mosaic_biomass$Generation_Time)

# The plot

plot(mosaic_biomass$log10_biomass, mosaic_biomass$log10_gent,
     pch=16, cex=1.5, col=alpha("black", 0.5),
     xlab=expression(paste(log[10], "Biomass")),
     ylab=expression(paste(log[10], "Generation time"))
```

```
)

# Simple linear regression (note that we are skipping model fitting in this b
rief vignette)

lmBioGenT = lm(log10_gent~log10_biomass, data = mosaic_biomass)

summary(lmBioGenT)
##
## Call:
## lm(formula = log10_gent ~ log10_biomass, data = mosaic_biomass)
##
## Residuals:
##      Min       1Q   Median       3Q      Max
## -0.53325 -0.16107 -0.01597  0.19936  0.38382
##
## Coefficients:
##               Estimate Std. Error t value Pr(>|t|)
## (Intercept)    0.85966    0.25372   3.388  0.00253 **
## log10_biomass  0.02762    0.05728   0.482  0.63418
## ---
## Signif. codes:  0 '***' 0.001 '**' 0.01 '*' 0.05 '.' 0.1 ' ' 1
##
## Residual standard error: 0.2623 on 23 degrees of freedom
## Multiple R-squared:  0.01001,    Adjusted R-squared:  -0.03303
## F-statistic: 0.2326 on 1 and 23 DF,  p-value: 0.6342
abline(lmBioGenT, col="red", lty=2, lwd=2)
```

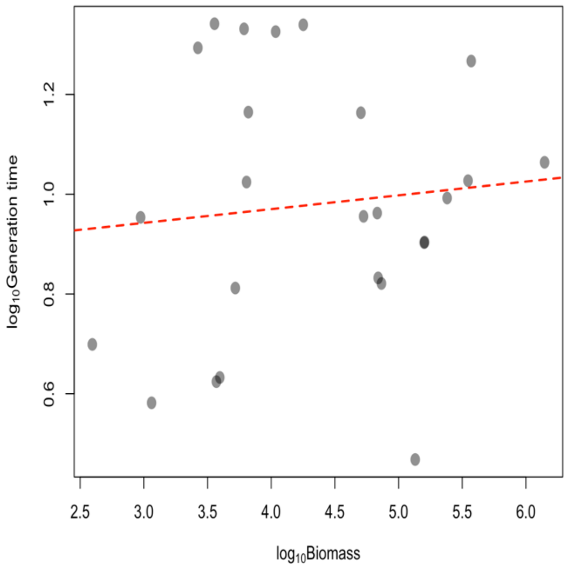
1.2
1.0
0.8
0.6
$log_{10}$Generation time
2.5
3.0
3.5
4.0
4.5
5.0
5.5
6.0
$log_{10}$Biomass

#### S5: LogNormal Distribution of Mass and Height

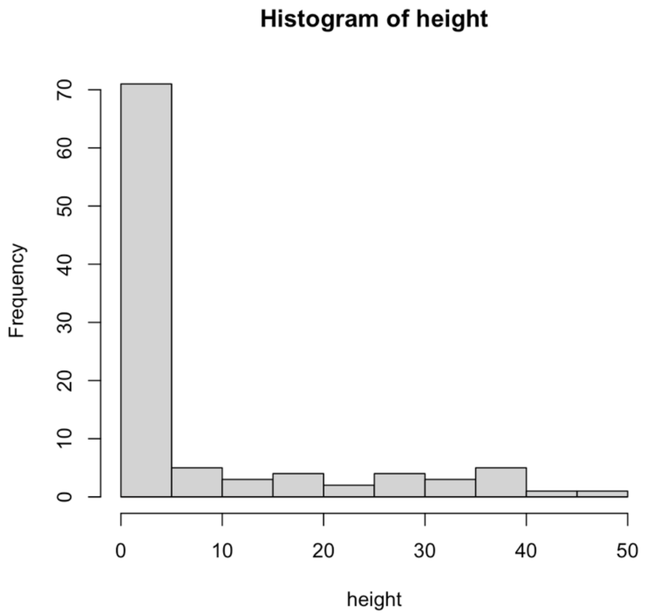

Raw heights from MOSAIC v1.0.0

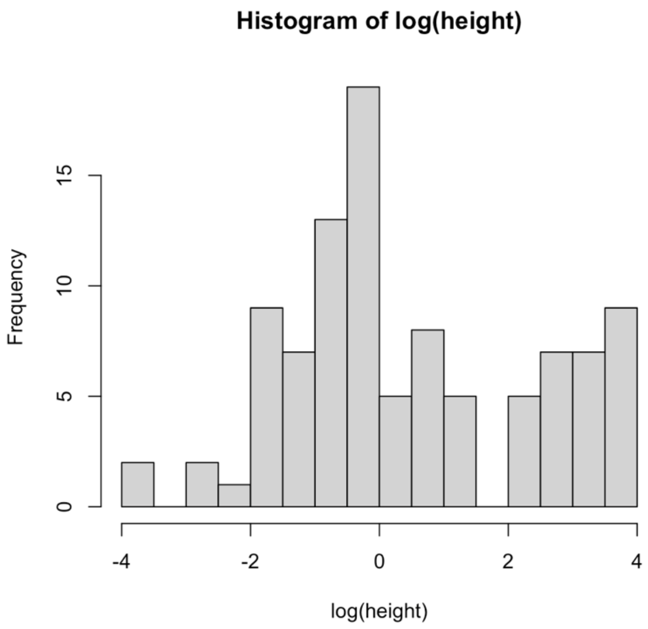

Log Heights from MOSAIC v.1.0.0

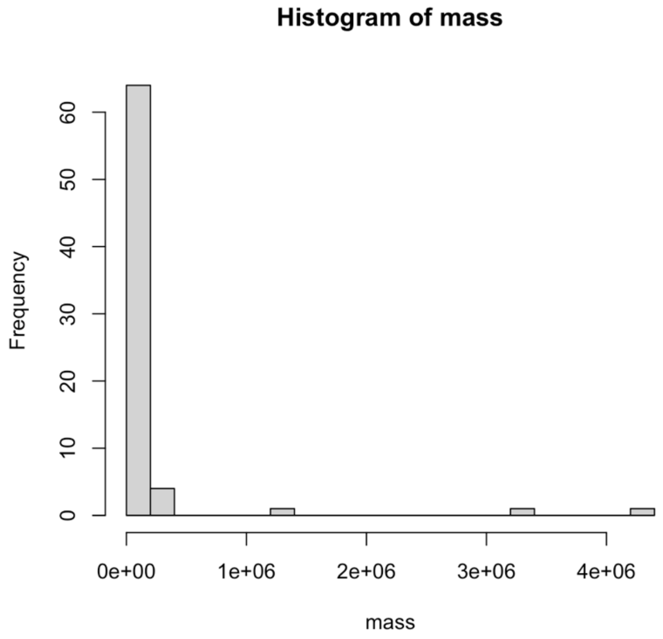

Raw mass from MOSAIV v1.0.0

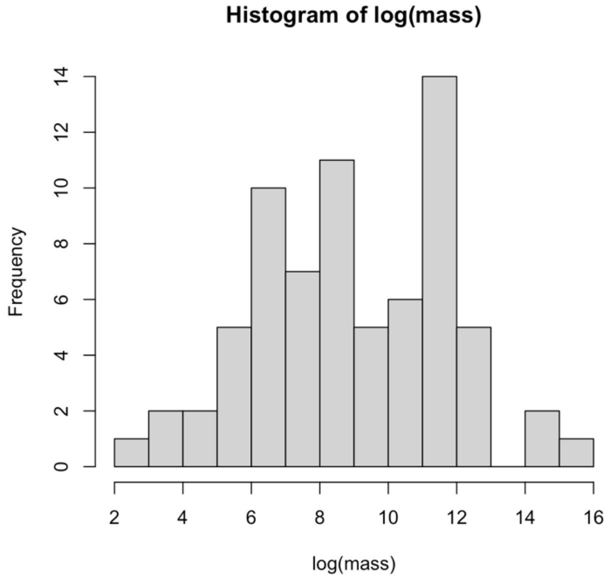

Log mass from MOSAIC v1.0.0
